# A chromosome-scale assembly of allotetraploid *Brassica juncea* (AABB) elucidates comparative architecture of the A and B genomes

**DOI:** 10.1101/681080

**Authors:** Kumar Paritosh, Satish Kumar Yadava, Priyansha Singh, Latika Bhayana, Arundhati Mukhopadhyay, Vibha Gupta, Naveen Chandra Bisht, Jianwei Zhang, David Kudrna, Dario Copetti, Rod A Wing, Vijay Bhaskar Reddy Lachagari, Akshay Kumar Pradhan, Deepak Pental

## Abstract

*Brassica juncea* (AABB; genome size ∼920 Mb), commonly referred to as mustard, is a natural allopolyploid of two diploid species – *B. rapa* (AA) and *B. nigra* (BB). We report a highly contiguous genome assembly of an oleiferous type of *B. juncea* variety Varuna, an archetypical Indian gene pool line of mustard, with ∼100x PacBio single-molecule real-time (SMRT) reads providing contigs with an N50 value of >5Mb. Assembled contigs were corrected and scaffolded with BioNano optical mapping. Three different linkage maps containing a large number of GBS markers were developed and used to anchor scaffolds/contigs to the 18 linkage groups of *B. juncea*. The resulting chromosome-scale assembly is a significant improvement over the previous draft assembly of *B. juncea* Tumida, a vegetable type of mustard. The assembled genome was characterized for transposons, centromeric repeats, gene content, and gene block associations. Both A and B genomes contain highly fragmented gene block arrangements. In comparison to the A genome, the B genome contains a significantly higher content of LTR/Gypsy retrotransposons, distinct centromeric repeats and a large number of *B. nigra* specific gene clusters that break the gene collinearity between the A and the B genomes. The genome assembly reported here will provide a fillip to the breeding work on oleiferous types of mustard that are grown extensively in the dry land areas of South Asia and elsewhere.

## Introduction

Genus Brassica contains some of the most important oilseed and vegetable crops that are grown worldwide. Nagaharu, based on cytogenetic studies, proposed a model on the relationship of six key species of the genus^1^. The model, known as ‘U’s triangle has three diploid species – *B. rapa* (AA, n=10), *B. nigra* (BB, n=8) and *B. oleracea* (CC, n=9) on the nodes and three allotetraploids – *B. juncea* (AABB, n=18), *B. napus* (AACC, n=19) and *B. carinata* (BBCC, n=17) at the coordinates. *B. rapa* was the first of the six species to be sequenced^2^ followed by *B. oleracea*^3^, *B. napus*^4^ and *B. juncea*^5^. Draft genomes of *B. rapa* (estimated genome size ∼ 485 Mb), *B. oleracea* (∼ 630 Mb) and *B. napus* (∼1130 Mb) were assembled with short-read, high-throughput Illumina technology with some limited Sanger sequencing^2–4^. A draft sequence of a vegetable type of *B. juncea* variety Tumida (genome size ∼922 Mb) was assembled^5^ using ∼176x Illumina reads, gap filling with 12x PacBio single-molecule real-time (SMRT) sequences and long-range scaffolding with BioNano optical mapping.

The long-read third generation technologies such as SMRT sequencing by PacBio and Nanopore sequencing combined with long-range scaffolding technologies have significantly improved genome assemblies providing higher levels of contiguity and more extensive coverage of repeat sequences, transposable elements, centromeric and telomeric regions^6,7^. Recent genome sequencing efforts have relied exclusively on high coverage SMRT sequencing, both for the de-novo genome assemblies and for improving earlier assembled draft genomes that used the second-generation technologies^8–13^. Amongst the Brassica species – assembly of *B. rapa* Chiifu has been improved by additional data from ∼57x SMRT sequencing, BioNano optical mapping, and Hi-C reads^14^. Highly contiguous de-novo assemblies have been generated for two new lines of *B. rapa* and *B. oleracea* by Nanopore long-read sequencing and optical mapping^15^.

We report here reference genome assembly of an oleiferous type of *B. juncea* – variety Varuna, an archetypical line of the Indian gene pool of mustard, with ∼100x SMRT coverage using PacBio Sequel chemistry and long-range scaffolding with BioNano optical mapping. To facilitate and validate the assembly of the two constituent genomes – A and B of *B. juncea,* a draft genome of an Indian gene pool line of *B. nigra* (variety Sangam) was assembled using Illumina shotgun sequencing and gap filling with Oxford Nanopore long-read sequences. Sequence assemblies were further validated with three new genetic maps containing a large number of GBS markers.

The assembled genome of *B. juncea* variety Varuna, consisting of 58 scaffolds and nine contigs assigned to 18 pseudochromosomes, is a significant improvement over the previous assembly of *B. juncea* variety Tumida^5^ – particularly, for the B genome. The Tumida assembly (Version 1.1) consisted of 10,581 scaffolds with N50 of ∼1.5Mb; in the Varuna assembly N50 of the scaffolds is ∼33.6 Mb. Further, the centromeric and telomeric regions were identified in most of the 18 pseudochromosomes. We compare the architecture of the A, B and C genomes and highlight both similar and unique features of the B genome vis-à-vis the A and C genomes.

## Results

### Genome assembly

To assemble the genome of *B. juncea* (AABB, genome size ∼922Mb^5^), three different experimental activities were initiated concurrently. In the first activity, high molecular weight DNA isolated from *B. juncea* variety Varuna was subjected to SMRT sequencing on the PacBio RSII platform. A total of 9,735,857 reads with an N50 value of ∼15.5 kb were obtained – providing ∼100x coverage of the genome (Supplementary Table 1). The reads were assembled into 1,253 contigs with a N50 value of ∼5.7 Mb using Canu assembler (Table 1). *B. juncea* genome was also sequenced on an Illumina HiSeq platform to obtain ∼40x coverage of the genome (Supplementary Table 2). Around 98.5% of the short-reads could be mapped to the SMRT sequencing based contigs, indicating that the long-read sequences had covered most of the genomic regions. In the second activity, a doubled haploid (DH) line of *B. nigra* (BB) variety Sangam, genome size ∼591 Mb^5^, was sequenced using Illumina HiSeq platform. Around 110x coverage was obtained by sequencing paired-end (PE) libraries of three different insert sizes (Supplementary Table 3a). An additional 2.8 Gb sequence data were obtained from three different mate-pair (MP) libraries (Supplementary Table 3b). The paired-end reads were assembled to obtain 57,941contigs with an N50 value of ∼34.6 kb. These contigs were scaffolded using ∼14x Oxford Nanopore long-read sequence data (Supplementary Table 3c) followed by mate-pair library data. These processes resulted in an assembly consisting of 18,996 scaffolds with an N50 value of ∼85.9 kb covering around 470.8 Mb of the *B. nigra* genome (Supplementary Table 4).

**Table 1.** Genome assembly statistics of *Brassica juncea* variety Varuna.

| *B. juncea* (AABB, 2n=18) |  |  |  |
| --- | --- | --- | --- |
| PacBio | ✓ | ✓ | ✓ |
| BioNano |  | ✓ | ✓ |
| Linkage map |  |  | ✓ |
| Total assembly size(bp) | 869,539,390 | - | - |
| Number of contigs | 1,253 | - | - |
| Longest contig (bp) | 35,843,917 | - | - |
| N50 contig length (bp) | 5,734,093 | - | - |
| Number of Scaffolds | - | 58 | 18 |
| Longest Scaffold(bp) | - | 72,092,522 | 73,851,810 |
| N50 scaffolds length(bp) | - | 33,612,814 | 58,609,396 |
| Scaffolds assigned to LGs | - | - | 54 +9 contigs |
| Unassigned contigs |  | 714 | 705 |
| Unassigned scaffolds | - | - | 4 |
| Length of assigned sequences(bp) | - | 847,778,585 | 841,233,580 |
| Length of Unassigned sequences(bp) | - | 36,400,777 | 35,468,516 |
| Missing bases (%) | 0 | 3.29 | 3.43 |

In the third activity, three genetic maps were developed – two for *B. juncea* and one for *B. nigra*. An earlier developed genetic map of *B. juncea* using an F_1_DH population derived from a cross of Varuna and Heera (an east European gene pool line) – named VH population^16^, which contained 833 intron polymorphic (IP), genic SSR and SNP markers^17^, was further enriched by an addition of 2,947 GBS markers using *Sph*I-*Mluc*I enzyme combination (Supplementary Table 5; Supplementary Fig. 1). A second F_1_DH population (named TuV) was developed in this study from a cross between Tumida and Varuna. A population of 119 individuals was first mapped with 524 IP and SSR markers that were common with the VH population – subsequently, 8,517 GBS markers, using *Hinf*I-*Hpy*CH4IV enzyme combination, were added for a more extensive coverage (Supplementary Table 6; Supplementary Fig. 2). A third F_1_DH mapping population was developed from a cross between *B. nigra* variety Sangam (an Indian gene pool line) and line 2782 (an east European type line). This map contained, besides a common set of IP and genic SSR markers with the VH population, an additional 2,723 GBS markers (Supplementary Table 7; Supplementary Fig. 3).

As we intended to use the *B. nigra* assembly to identify the contigs specific to the B genome in the *B. juncea* SMRT sequence assembly, we first developed a reference genome of *B. nigra*. The genetic map of *B. nigra* was used for assigning scaffolds to the eight LGs – 14,147 scaffolds could be assigned to the LGs to constitute eight pseudochromosomes (Supplementary Table 4; Supplementary Fig. 4); 4,849 scaffolds remained unassigned. To separate the A and B genome specific contigs of *B. juncea* – scaffolds of *B. nigra* were compared to the 1,253 SMRT based contigs of *B. juncea*. Of the 1,253 contigs – 691 were found to belong to the B genome. The remaining contigs were checked against the draft genome sequence of *B. rapa*^2^, and 505 contigs were found to be specific to the A genome; 57 contigs could not be specified to either of the two genomes. Around eighteen mis-assemblies were noticed – these had mostly arisen due to the coalescence of highly conserved syntenic regions. These mis-assemblies were confirmed by the high-density genetic map of VH population. Scaffolding was carried out by hierarchical mapping of the 1,253 SMRT contigs on three different optical maps. Two maps were developed with NLRS based labeling using *BssS*I and *BspQ*I enzymes and one with DLS labeling using DLE-1 enzyme (Supplementary Table 8; Supplementary Fig. 5). Corrections were made at each mapping stage, and corrected sequences were mapped again on the consensus maps. A total of 45 contigs were identified with mis-assemblies. These included the eighteen mis-assemblies that were identified earlier. These were curated by breaking the mis-assembled regions and aligning the edited contigs again to the consensus maps. The final assembly, after corrections and multiple rounds of scaffolding, was constituted of 58 scaffolds and 714 unscaffolded contigs/fragments. The total size of the assembled scaffolds was ∼847.8 Mb with an N50 value of ∼33.6 Mb (Table 1). The unscaffolded contigs/fragments constituted ∼36.4 Mb of the genome with an N50 value of ∼55.6 kb.

The VH genetic map was used to assign the scaffolds and contigs to 18 linkage groups of *B. juncea* and also to assess the quality of the assembly. Fifty-one of the 58 scaffolds and nine of the 714 contigs could be assigned to the 18 pseudochromosomes – which were designated as BjuVA01 - BjuVA10, BjuVB01 - BjuVB08^17^ (Figure 1; Supplementary Table 9). Three of the pseudochromosomes were covered with only one scaffold, four with two scaffolds, six with three and the rest with four or more scaffolds and a few contigs – the maximum number being ten for BjuA01. GBS markers, 2,947 in number that mapped on the VH population, were used to compare their position on the genetic map with their physical location on the pseudochromosomes. A very high correlation was observed between the marker order on the genetic map and the physical position of the GBS tags on the pseudochromosomes (Supplementary Fig. 6). To further validate the assembly, we used the TuV population genetic map, which contained 8,517 GBS markers assigned to the 18 LGs of *B. juncea*. The order of the dispersed GBS markers on the TuV genetic map was invariably found to be correlated with the order of their sequences on the pseudochromosomes (Supplementary Fig. 7). Based on the genetic marker data, two more scaffolds could be assigned to the pseudochromosomes (Supplementary Table 9). At the end of all the steps, from the overall genome assembly consisting of 58 scaffolds and 714 contigs/fragments – 54 scaffolds and nine contigs could be assigned to the 18 pseudochromosomes; four scaffolds and 705 contigs remained unassigned (Table 1). The size of the assigned sequences was calculated to be around 841.2 Mb (∼91.2% of the estimated genome content). About 35.4 Mb of the sequenced genome (∼3.7% of the total genome) remained unassigned. A comparison of the chromosomes level assembly of *B. juncea* Varuna genome with the assembly earlier reported for *B. juncea* Tumida (V1.1)^5^, and its improved version V1.5^18^ showed a high level of contiguity and lesser number of gaps in the Varuna assembly (Supplementary Table 10). The coverage achieved for the B genome in the present study was significantly higher – ∼507.7 Mb, as compared to the earlier reported coverage of ∼395.9 Mb in V1.1^5^ and ∼377.6 Mb in V1.5^18^ (Supplementary Table 10).

**Figure 1.**
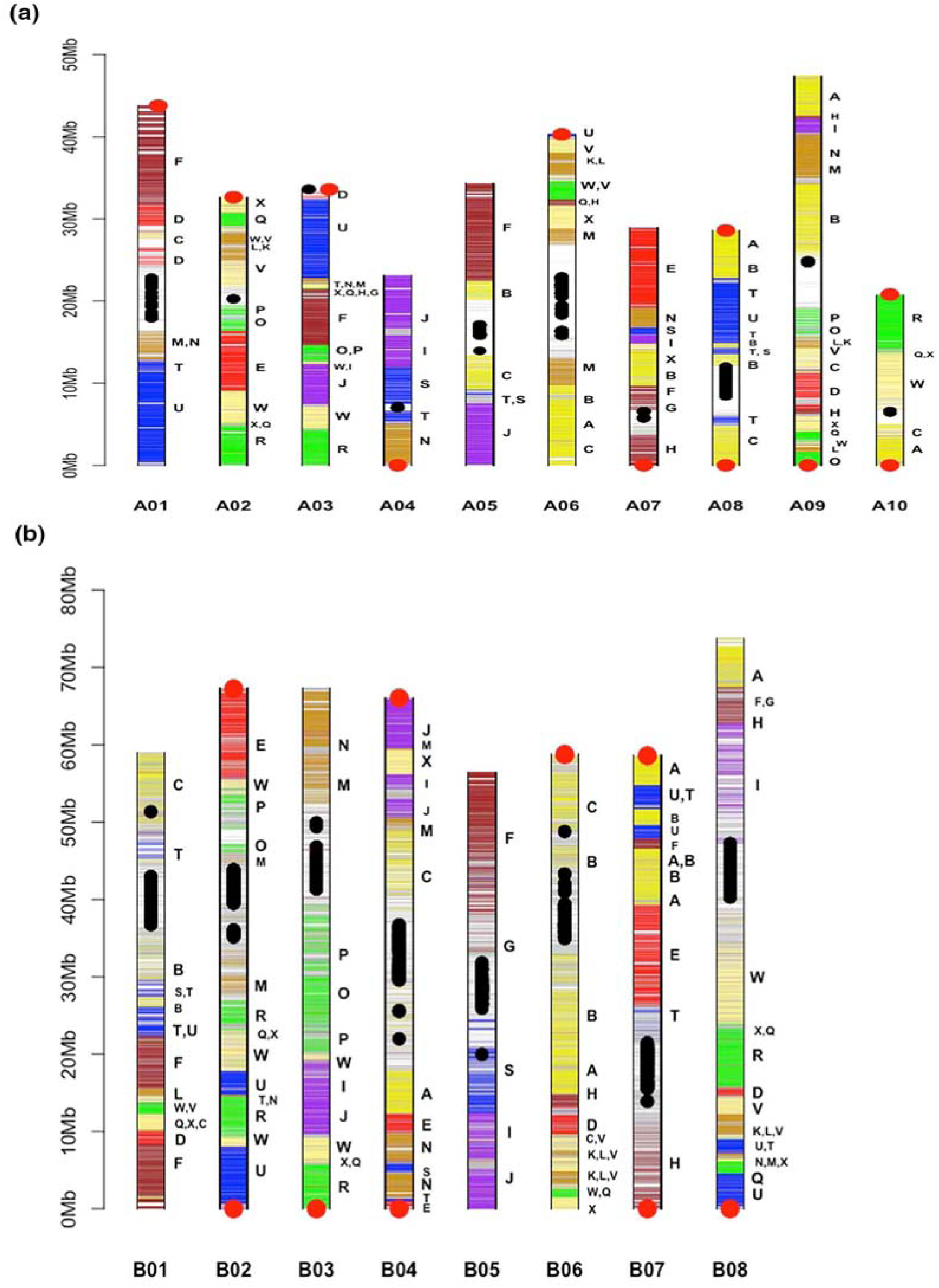
Graphic representation of *Brassica juncea* (i) A genome and (ii) B genome pseudochromosomes. Horizontal bars represent the predicted genes. Different regions of the pseudochromosomes have been assigned to gene blocks on the basis of synteny with *Arabidopsis thaliana* (At) gene blocks (A–X) as defined by Schranz et al., (2006) ^26^; same color code has been used for *B. juncea* gene blocks as used for the gene blocks in At. Centromeric repeats are represented with black dots; telomeric repeats, wherever found, are represented as red dots.

### Genome annotation

*B. juncea* Varuna genome assembly was annotated for three components – transposable elements, centromeres, and genes. Transposable elements (TEs) were identified by structure and similarity using a *de-novo* prediction approach (Methods for details). This exercise generated a database of 1,590 consensus repeat sequences that were merged with the *Arabidopsis thaliana* (At) TE database to annotate TEs in the assembled pseudochromosomes of the A and B genomes, separately. Repeats were broadly classified under three major categories – retrotransposons, DNA transposons, and other repeats and further subclassified (Supplementary Table 11). Around 385Mb (∼45.8%) of the assembled genome of Varuna was found to be constituted of TEs. The B genome had a higher repeat content ∼259Mb (∼51%) as compared to the A genome – 113Mb (∼33.9%). Retrotransposons – LTR/Copia and LTR/Gypsy were the predominant TEs present in the Varuna genome. Both these types showed significant expansion in the B genome. LTR/Copia constituted ∼4.4% (∼14Mb) of the A genome; in comparison – these elements constituted around ∼9% (46 Mb) of the B genome. The expansion of LTR/Gypsy was most pronounced in the B genome – ∼21.5% (∼109Mb) in comparison to ∼7.5% (∼25Mb) in the A genome (Supplementary Fig. 8, Supplementary Table 11).

Candidate centromeric regions were identified from the correlation plots between GBS markers on the genetic maps and the physical positions of their respective tags on the pseudochromosomes (Supplementary Fig. 6, 7); regions with significantly low recombination frequencies were marked as potential centromeric regions and analyzed for repeats that were absent from other parts of the pseudochromosomes (details in Methods). CentBr1 and CentBr2, identified as centromere-specific sequences in *B. rapa* (A genome) and *B. oleracea* (C genome) in earlier studies^2,3^ were found to be present in the A genome of *B. juncea* but absent from the centromeric regions of all the pseudochromosomes of the B genome. Three new A genome-specific centromeric repeats, and seven B genome-specific repeats were identified (Supplementary Table 12, 13; Supplementary Fig. 9, 10). Further, the centromeric/ pericentromeric regions of all the B genome pseudochromosomes contained a much higher content of TEs as compared to such areas in the A genome (Supplementary Fig. 8, Supplementary Table 14).

For gene annotation, we performed RNA-seq analysis on poly-A enriched RNA isolated from seedling and young inflorescence tissues of Varuna on a PacBio RSII platform. A total of 40,208 full-length high-quality sequences were obtained (details provided in Supplementary File 1). Genome sequences assigned to the 18 pseudochromosomes of Varuna were repeat-masked (see Methods) and analyzed for gene content with Augustus software, using 543 randomly selected full-length CDS obtained from the PacBio based RNA-seq analysis as the training dataset. A total of 105,354 genes were predicted of which 48,270 in the A genome and 57,084 in the B genome and assigned to the individual chromosomes (Supplementary Table 15). The predicted genes were validated by comparing these with the non-redundant proteins in the UniProt plant database. This analysis could validate around 93% of the predicted genes. Predicted genes were also confirmed by the Iso-seq transcriptome analysis and short-read RNA-seq data generated for *B. juncea* in some other studies^17,19,20^ (Supplementary File 1). A total of 82,008 genes were found to express in one or the other study – 38,232 (∼79%), 43,776 (∼76%) in the A and B genome, respectively. We also carried out RNA-seq of *B. nigra* Sangam using RNA isolated from the seedling and young inflorescence tissues on Illumina and Roche sequencing platforms (Supplementary File 2). For the *de-novo* prediction of the genes in the assembled *B. nigra* genome, the eight pseudochromosomes and unassigned scaffolds of *B. nigra* were used for gene prediction with Augustus software using the same training dataset as used for *B. juncea* gene prediction. A total of 46,227 genes were predicted, of which 35,981 were found to express in the RNA-seq carried out on *B. nigra* tissues (Supplementary File 2). All the expressed genes in *B. nigra* were found to be present in the annotated B genome of *B. juncea* Varuna.

A dataset was generated for *B. juncea* with information on the physical position of every predicted gene on the 18 pseudochromosomes – A01-A10, B01-B08 (Supplementary Table 15). This dataset also contains information on the expression status of each of the predicted gene and its closest ortholog in *B. rapa* Chiifu and the model species At based on protein similarity. Genes were marked based on 24 ancestral genomic blocks (A-X) defined for the At genome^21^ (Fig. 1, Supplementary Table 15). We carried out ortholog tagging of the *B. juncea* Varuna genes with the previously assembled *B. rapa* Chiifu (V1.5) and *B. juncea* Tumida (V1.1 and V1.5) assemblies; many mis-assembled regions in the previous assemblies could be identified (Supplementary Fig. 11). A total of 35,644 (88.3%) and 1802 (92.5%) of the contigs/scaffolds that remained unassigned in the previous *B. rapa* (V1.5) and *B. juncea* Tumida (V1.1, V1.5) assemblies, respectively, could be placed on the 18 pseudochromosomes of the *B. juncea* Varuna (Supplementary Table 16). Mis-assemblies were found to be more pronounced in the assembly of the B genome of *B. juncea* Tumida.

Orthologous gene groups between the A and B genomes of *B. juncea* were identified based on protein similarity using OrthoFinder^22^. A total of 19,404 orthologous groups were identified between the A and B genomes; 39,383 genes of A had orthologs in B, and 42,727 genes of B had orthologs in the A genome. Further, 5,942 A genome-specific and 22,138 B genome-specific genes were identified (Blastp E-value <1e^−05^, coverage >80%). No orthologs of these genes were identified in At. A total of 2,937 A genome-specific genes and 12,262 of B genome-specific genes were observed to express in the RNA-seq data of *B. juncea* and *B. nigra* (Supplementary Fig. 12, 13). We checked whether the genome-specific genes were dispersed or present in clusters. A total of 152 gene clusters with ≥ten species-specific genes were identified in the B genome, whereas only 38 such clusters were observed in the A genome.

### Comparative genome architecture of A, B and C genomes

Syntenic genome segments between At and the A and B genomes of *B. juncea* were identified using MCScanX. Both A and B genomes have, with a few exceptions, a minimum of three syntenic regions to each of the gene block (A – X) of At. This can be attributed to genome triplication, referred to as the ***b*** event, which is considered to be the defining characteristic of the species belonging to the tribe Brassiceae^23^. The A and B genomes contained 24,696 and 27,125 orthologs of At, respectively; these were used to compare the gene retention/loss patterns in the syntenic regions of the A and the B genomes. This analysis confirmed the earlier reports on the presence of three paleogenomes in the A genome – LF_A_ (Least fractionated), MF1_A_ (Moderately fractionated) and MF2_A_ (Most fractionated)^2,24,25^. A similar trend was obtained in the B genome (Supplementary Fig. 14; Supplementary Table 17) – hence the three paleogenomes constituting the B genome were designated LF_B_, MF1_B_, and MF2_B_. Both in the A and B genomes, a significantly higher number of At gene orthologs were retained in the LF paleogenomes as compared to the genes retained in the MF1 and MF2 paleogenomes (χ^2^ test, p <0.05; Supplementary Table 17).

To check the evolutionary relationship between At and constituent paleogenomes of the A and B genomes, a set of 1,482 genes were identified that are present as single-copy genes in At, spread over all the gene blocks (A–X), and as three orthologs in both the A and B genomes. The divergence between At–A and At–B orthologs and paralogs within the A (LF_A_, MF1_A_, and MF2_A_) and the B (LF_B_, MF1_B_ and MF2_B_) genomes were studied by calculating synonymous nucleotide substitutions (Ks values) in pairwise comparisons using the PAML package^26^. Divergence values between At–A and At–B orthologs were not significantly different (one-way ANOVA, p=0.05). No significant variation was found among Ks values calculated in pairwise comparisons of paralogs present in the A and the B genomes (one-way ANOVA, p=0.05). However, the values in the former category were significantly higher than the values in the paralog comparisons (Supplementary Table 18).

Pairwise gene comparison was carried out to find the relationship between the LF_A_, MF1_A_, MF2_A_, LF_B_, MF1_B_ and MF2_B_ paleogenomes. This analysis showed LF_A_ – LF_B_, MF1_A_ – MF1_B_, MF2_A_ – MF2_B_ to be the most related combinations (Supplementary Fig 15, Supplementary Table 18) and therefore, can be referred to as homoeologs. A neighbor-joining tree was constructed based on the Ks values obtained with 1,482 genes belonging to At and the three constituent paleogenomes of A and B and additionally the C genome (Supplementary Fig 16). The split between the Arabidopsis and the Brassiceae progenitor occurred first (∼17 MYA). Paleologs in LF, MF1, and MF2 show antiquity of ∼ 15 MYA, homoeologs between A and B diverged ∼ 8.5 MYA, and homoeologs between A and C diverged ∼ 4.5 MYA. However, the timing of the ***b*** event could not be discerned from the analysis.

Based on the extent of gene fractionation (Supplementary Table 17) and divergence analysis (Supplementary Table 18, Supplementary Fig 15), we assigned all the gene blocks identified in the A and B genomes (Fig 1, Supplementary Table 15) to the three-constituent paleo-genomes LF_A_, MF1_A_, MF2_A_ and LF_B_, LF1_B_, MF2_B_ (Fig. 2). As a contiguous genome assembly of the C genome has been reported recently^15^– we also assigned the gene blocks in the C genome to the three constituent paleogenomes LF_C_, MF1_C_, and MF2_C_ (Fig. 2). The A and C genome homoeologs, besides being less divergent, also show extensive similarity in the gene block arrangements as compared to the A and B genome homoeologs (Fig 3).

**Fig. 2.**
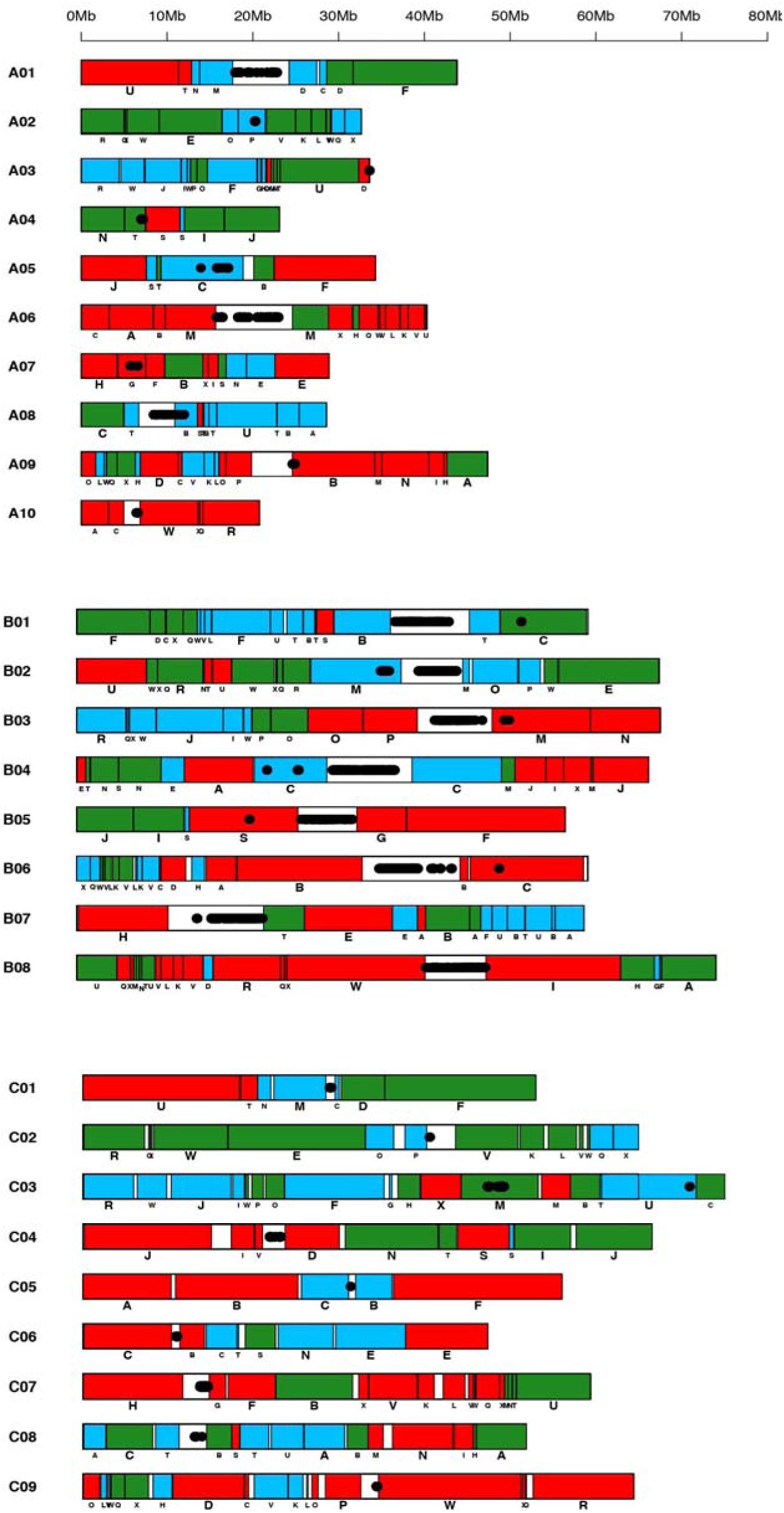
Gene block arrangements in the A and B genomes of *B. juncea* and C genome of B. *oleracea*. LF, MF1 and MF2 paleogenomes are represented by red, green and blue colors, respectively. A and C genomes show very similar block arrangements, whereas B genome shows more fragmentation of the gene blocks.

**Figure 3.**
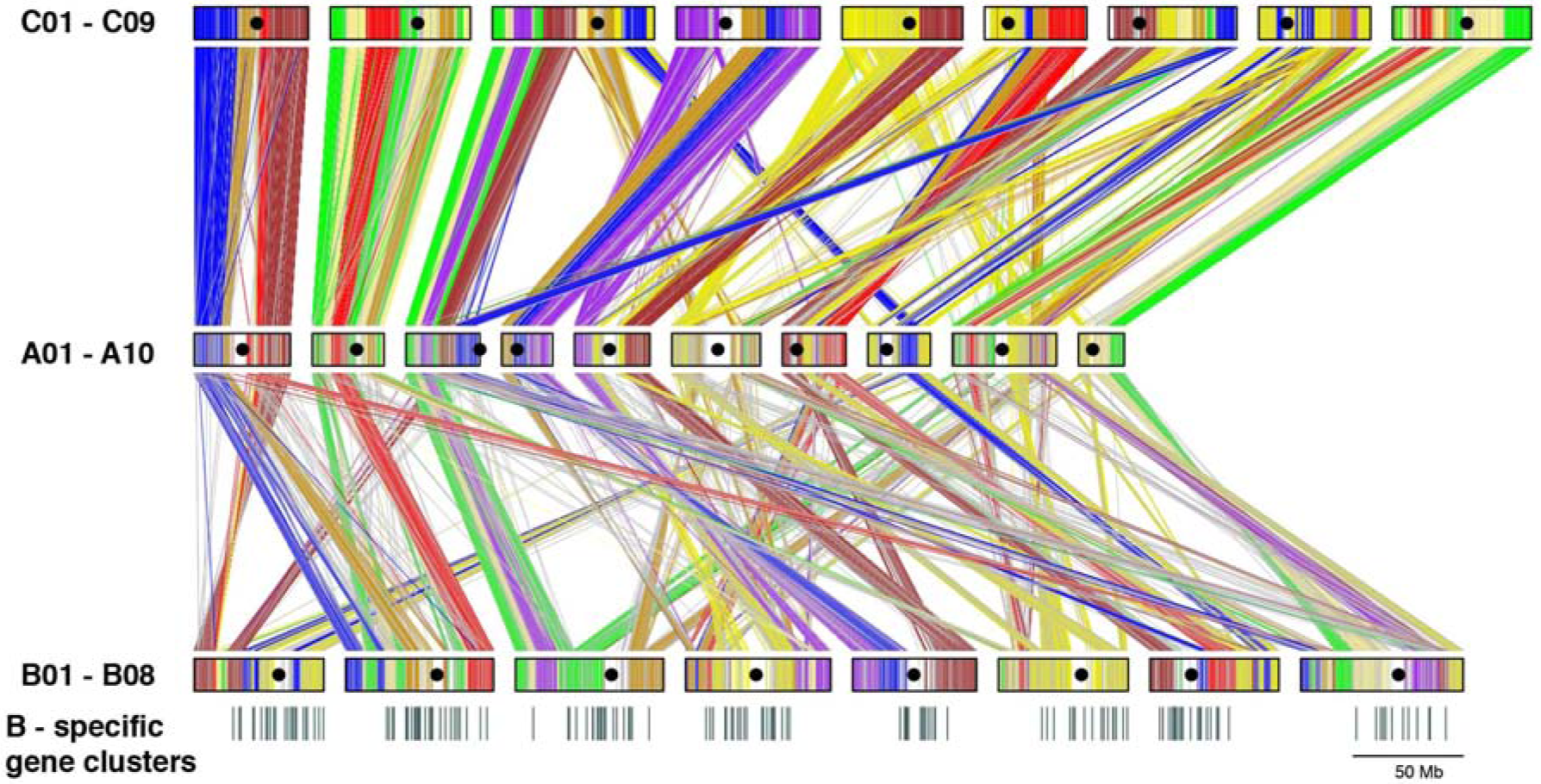
Segmental collinearity of the A, and B genomes of *Brassica juncea* and C genome of *B. oleracea*. Gene blocks syntenous with *A. thaliana* are color coded following Schranz et al., (2006)^26^. The A and C genomes have many similar gene block associations as compared to the A and B genomes. The B genome contains very high number of genome-specific genes many of which are present as clusters of ≥ 10 genes – such clusters have been represented as black bars below the B genome pseudochromosomes.

Gene block associations have been extensively studied in the selected taxa of the family Brassicaceae by *in situ* hybridizations^27^, genetic mapping^17,28,29^ and genome sequencing^30^ to understand the evolutionary processes. These studies have been summarized as – ‘presence and absence of ancestral genomic block associations in 35 species belonging to different tribes recognized in the family *Brassicaceae’*^30^ (Supplementary Table 19). More contiguous genome assemblies have shown the A, B, and C genomes to be even more fragmented than reported earlier^30^ with many intra-block fragmentations (Supplementary Table 20). We manually scanned the gene block associations and classified these under four categories – (a) Intra-paleogenome contiguous gene block associations (b) Intra-paleogenome non-contiguous gene block associations (c) inter-paleogenome non-contiguous gene block associations and (d) inter-paleogenome same-gene block associations (Supplementary Table 21).

The ancestral intra-paleogenome contiguous gene block associations – A–B, F–G, G– H, T–U, and W–X have been reported to be absent in the A and C genomes of *B. rapa, B. oleracea* and *B. napus* ^30^. The new assemblies of the A, B and C genomes show the presence of these block associations in one or the other paleogenomes (Fig. 2, Supplementary Table 19 and 21). Loss of these associations might have occurred pre- or post-***b*** event. Intra-paleogenomes non-contiguous gene block associations are more interesting for understanding the structure of the paleogenomes pre-***b*** event. The previously reported association V–K–L– W–Q–X as such does not exist in any of the paleogenomes, only V–K–L association is common to all the three paleogenomes (Supplementary Table 19 and 21). We report some new associations in this category – three of these namely, Q–X, W–X–Q–R and V–K–L are present in all the three paleogenomes; W–E, M–N–T–U, D–P –W are present only in MF1_A_, MF1_B_ and MF1_C_. These associations could be present in the paleogenomes before the ***b*** event. Inter-paleogenome non-contiguous gene block associations J_MF1_–I_MF1_–S_MF2_–S_LF_ and R_MF2_–W_MF2_–J_MF2_–I_MF2_–W_MF2_–P_MF1_–O_MF1_ present in all the three genomes, in all probability, have resulted from the ***b*** event. Inter-paleogenome same gene block associations could have resulted from homoeologous recombinations after the ***b*** event.

## Discussion

We have reported a highly contiguous genome assembly of *B. juncea* variety Varuna, an oleiferous type belonging to the Indian gene pool of mustard, using long-read SMRT sequencing and optical mapping. The *B. juncea* assembly reported here has given contig N50 value of >5Mb, comparable to the other recent SMRT based genome assemblies^9–14^. We found recursive optical mapping with three different labeling reactions to be highly useful for correcting mis-assemblies. The final assembly showed a strong correlation with genetic marker-based maps, which were exclusively developed for this study. The assembly is a marked improvement over the previous assembly of *B. juncea* variety Tumida – particularly, for the B genome.

Different models have been proposed to explain the evolution of Brassica species and other taxa in the tribe Brassiceae. One model suggested genome triplication (the ***b*** event) with three near-similar PCK^31^ (Proto-Calepineae Karyotype; n=7) genomes as the major event that precipitated differential gene loss, reduction in chromosome number and new gene block associations followed by changes resulting from homoploid hybrid speciation^32^. An alternative model suggested that two of the paleogenomes – MF1 and MF2 formed an allotetraploid with MF1 genome establishing dominance over the MF2 genome in terms of gene retention, followed by crosses with the LF genome (the ***b*** event), leading to chromosomal restructuring, new gene block associations and further gene fractionation^24^. Contiguous assemblies of the A, B and C genomes have provided evidence for a more extensive gene block fragmentation than reported earlier and some new insights into gene block associations. The data on intra-paleogenome non-contiguous gene block associations shows that the three paleogenomes LF1, MF1 and MF2 differed from one-another in gene block associations. However, the inter-paleogenome non-contiguous and same gene block associations have resulted either from the ***b*** event and from the later homoploid hybrid speciation. Thus, karyotype changes, new gene block associations, and gene block fragmentations as they exist in the A, B and C genomes, in all probability have resulted from multi-step genome rearrangements.

Oleiferous *B. juncea* types are grown extensively in the dry-land areas of South Asia and are a potential crop for other rainfed cropping regions of the world. In India, *B. juncea* is grown in around 5-6 million hectares of land during the winter growing season^33,34^. Our earlier work identified two distinct gene pools in oilseed mustard – Indian and east European^35,36^. Hybrids between the Indian and East European gene pool lines are heterotic for yield^35^. The Indian gene pool lines like Varuna are well adapted to the dry land cultivation but have a narrow genetic base^37^. The east-European lines, as such, are ill-adapted to the growing conditions in South Asia due to photoperiod sensitivity but contain a large number of positive traits like high pod density^38–40^, oil and seed meal quality^41–42^, and disease resistance^43^. Several QTL have been mapped in the Indian and East European gene pool lines ^38–44^. The Chinese vegetable types like Tumida constitute another divergent gene pool^5^ that contains potentially useful traits like serrated leaves, basal branching^45^, and stem strength. The highly contiguous genome sequence assembly of *B. juncea* will allow fine mapping in the QTL regions, identification of the candidate genes and understanding molecular mechanisms underlying some of the critical quantitative traits.

## Materials and Methods

### Plant Material

*Brassica juncea* (AABB) variety Varuna, an oleiferous type belonging to the Indian gene pool of mustard, is one of the most extensively grown mega variety released by the public system in India^33^. Varuna material used for sequencing was maintained by strict selfing for over six generations. The other material used for sequencing – *B. nigra* (BB) variety Sangam (an Indian gene-pool line) is used extensively for condiment purposes in India. As *B. nigra* is self-incompatible and therefore highly heterozygous, sequencing was carried out on a derived doubled haploid (DH) line – named BnSDH-1 – which was maintained by selfing through bud pollination. A mapping population of *B. juncea* Varuna x Heera (VH population) was developed earlier^16,28^. A new F_1_DH mapping population was developed in *B. juncea* from a Tumida x Varuna cross (TuV population). A *B. nigra* F_1_DH mapping population was developed from the cross – BnSDH-1 x line 2782 (an exotic east European origin line). The DH mapping populations were maintained by strict self-pollination in the field during the growing season.

### PacBio library construction and sequencing

For DNA isolation *B. juncea* variety Varuna seedlings were grown in a growth chamber (light period 8 hrs, temp. 25 °C/ dark period 16 hrs, temp. 10 °C). For DNA isolation, 15 d old seedlings were kept in the dark for 24 hrs; young leaves were harvested and frozen immediately in liquid nitrogen. Leaf tissues were used to isolate the nuclear fraction; high-molecular-weight genomic DNA was extracted from the isolated nuclei and subjected to pulse-field gel electrophoresis for separating DNA in the range of 30-50 kb for library preparation. Libraries were developed using the SMRTbell^™^ template preparation kit following the manufacturer’s instructions (https://www.pacb.com/) and sequenced on the PacBio RSII platform (https://www.pacb.com).

### Illumina and Nanopore library construction and sequencing

*B. juncea* and *B. nigra* plants were grown in a growth chamber under conditions described above and used for DNA isolation by CTAB method^46^. DNA was purified using DNase Mini kit (Qiagen). Approximately 5 µg of genomic DNA was fragmented with a focused ultrasonicator system (Covaris). Fragmented DNA was used to prepare Illumina paired-end (PE) libraries of size 200-350 bp for *B. juncea* var Varuna following the manufacturer’s recommended protocol (https://illumina.com). For *B. nigra* three different PE libraries of sizes 200-350 bp, 300-450 bp and 400-550 bp and three mate-pair (MP) libraries with the size range of 2-3 kb, 4-6 kb, and 10 kb were constructed as per the recommended protocols. The PE libraries were sequenced on Illumina Hiseq1000 sequencer and MP libraries on Illumina MiSeq system (Illumina). For Oxford Nanopore technology (https://nanoporetech.com) based genome sequencing of *B. nigra*, around 3 µg of high molecular weight purified DNA was used directly for developing sequencing libraries following “1D gDNA selection for long reads” protocol using the ligation sequencing kit 1D (Cat no: SQK-LSK108) recommended by the manufacturer ((https://nanoporetech.com). Developed libraries were sequenced with R9.4 SpotON Minion-106 Flow cells (Cat no: FLO-MIN106). Base calling was performed upon completion of the sequencing runs with Albacore software (https://github.com/Albacore).

### Assembly of *B. juncea* and *B. nigra* genome

Sequencing of *B. juncea* libraries on PacBio Sequel platform generated > 9 million long reads; these were processed by Canu assembler^47^ (V1.4) – “minReadLength” and “minOverlapLength” set at 1000bp “rawErrorRate” set at 0.3 and “correctedErrorRate” at 0.045. Completeness of the assembly was checked by mapping sequences obtained from the Illumina PE libraries on the assembled SMRT based contigs using BWA mem^48^ (default parameters).

Sequences generated from the three PE libraries of *B. nigra* were assembled into contigs with MaSuRcA software^49^. Illumina PE reads were mapped on the raw Nanopore reads using BWA mem; 1,442,476 error corrected Nanopore reads were generated using Pilon program^50^. Scaffolding of the MaSuRcA based contigs with the error corrected Nanopore sequence data was carried out with SSPACE-LongRead.pl script^51^ which generated 29,808 scaffolds with an N50 value of 19,834. Further, three rounds of scaffolding were carried out using the 2 kb, 6 kb and 10 kb MP library sequence datasets using SSPACE-STANDARD-3.0.pl script^51^.

### BioNano assembly and validation

BioNano optical maps were developed with proprietary kits and protocols provided by BioNano Genomics (https://bionanogenomics.com). High molecular weight DNA was isolated from 1g tissue of freshly harvested etiolated young leaves of *B. juncea* using “IrysPrep Plant Tissue DNA isolation kit (Cat no: 80003)”. Approximately 750 ng of the genomic DNA was trapped in agarose gel plugs and labeled at nicks introduced by BssSI and BspQI enzymes in separate reactions using the “IrysPrep NLRS labeling kit (Cat no: RE-012-10)”. Around 900 ng of high molecular weight DNA was used for DLS labeling using “DNA labeling kit-DLS (Cat no: 80005)”. For each reaction, labeled DNA was imaged on the Saphyr system (https://bionanogenomics.com) using three lanes of the Saphyr Chip (Cat no: FC-030-01). For each of the three libraries, pairwise comparison was performed with RefAligner to identify all molecule overlaps, and three different consensus maps were constructed. All fragments were then mapped back to the consensus maps which were recursively refined and extended. The BioNano IrysSolve module ‘HybridScaffold’ was used to perform the hybrid assembly between PacBio-SMRT contigs and BioNano-assembled genome maps.

### Genetic mapping

Marker-based genetic maps were developed from two F_1_DH populations of *B. juncea* – Varuna x Heera (VH) and Tumida x Varuna (TuV). VH population was developed earlier and contained 709 – Intron polymorphic (IP), genic SSR, and SNP markers^28^. A subset of 92 F_1_DH lines of the VH population along with the two parents was used for GBS (Genotyping by Sequencing) based mapping using *Sph*I-*Mluc*I restriction enzyme combination^52^. PE reads that were generated from the sequencing of the F_1_DH individuals, and the two parental lines were mapped to the scaffolds and contigs generated with PacBio SMRT sequencing and Optical mapping using BWA. SNPs were identified, and potential markers were filtered – markers with read-depth <10, monomorphic markers and markers with >30 missing data in the population were rejected. Genetic mapping was carried out in the R statistical computing environment using the ASMap package^53^. The R/ASMap package uses an R/qtl^54^ format for the structure of its genetic objects and utilizes the MSTmap algorithm^55^. Marker data were assembled to be used as a DH population type and transformed into the R/qtl format. The genetic map was calculated by using the mstmap function with the default parameters: distance function Kosambi, cut off p-value 1e-10 and missing threshold 0.3. GBS marker tags of the VH population were used to anchor the assembled super-scaffolds and contigs on the genetic map. Anchored scaffolds and contigs were stitched into 18 pseudochromosomes using custom Perl scripts. Correlations were drawn between the GBS markers on the genetic map and their physical position on the pseudochromosomes using custom R scripts.

For the genetic map of TuV population, a total of 119 lines were used – first for genotyping with 524 anchor markers (179 IP, 184 SNP, and 161 SSR) that were common with the VH population. Subsequently, 8,517 GBS markers were mapped using the same parameters as used for the VH genetic map, except that the DNA was restricted with the *Hinf*I-*Hpy*CH*4*IV enzyme combination that is reported to provide more extensive genome coverage^56^. PE reads that were generated from the sequencing of the F_1_DH individuals and the two parental lines were mapped to the 18 pseudochromosomes and the scaffolds and contigs that remained unassigned after analysis with the VH genetic markers using BWA. GBS marker filtering, genetic mapping, and correlations in the form of dot plots were drawn between the physical positions of the GBS based SNPs and their genetic locations on the TuV genetic map as done for the VH map.

For genetic mapping in *B. nigra*, 92 F_1_DH population (Sangam × 2782) developed and a framework genetic map was developed with 208 anchor markers (180 IP and 28 SSR) that were common with the VH anchor map. Further, 2,723 GBS markers (enzyme combination *Sph*I-*Mluc*I) were added using the mapping procedures described above. *B. nigra* pseudochromosomes were developed by organizing the assembled scaffolds as per the order of the markers on the genetic map.

### Transcriptome sequencing

For *B. juncea*, PacBio based transcriptome sequencing (Iso-seq) was carried out on total RNA isolated from a pooled sample of seedlings, leaves, inflorescence with developing siliqua along with the seeds using Spectrum^™^ plant total RNA kit (Sigma). Quality of RNA was checked on Bioanalyzer 2100 (Agilent) using RNA 6000 Nano kit. 5 µg of high-quality RNA (RIN value ≥7) was used for Iso-Seq SMRTBell library preparation (https://pacbio.secure.force.com/SamplePrep). Two size ranges were collected, one of size range 0.5-1kb and the other of 1-2 kb. Raw sequencing reads for each of the libraries were generated using P4-C6 chemistry on PacBio RS II sequencing machine. Sequencing data were generated from seven SMRTcells for ≤1 kb library and eight SMRTcells for 1-2 kb library. Sequences were assembled separately using SMRT Link software (https://www.pacb.com/). For validating the genes predicted in the Varuna genome, besides the Iso-seq data – different transcriptome datasets deposited in the NCBI SRA database (https://www.ncbi.nlm.nih.gov/sra) were downloaded and used. QC was carried out using Trimmomatic software^57^ and mapped on the assembled genome using STAR aligner^58^.

For transcriptome assembly of *B. nigra*, RNA was isolated from the seedlings, leaves and developing inflorescence tissues. 6 µg of RNA samples (RIN value >=7) was used for cDNA library preparation following the method described earlier^59^. PE library with an insert size range of 250-370 bp was developed and sequenced (2 × 100bp) using Genome Analyzer IIx sequencer (Illumina). Quality filtering and the raw reads were carried out with Trimmometic^57^ and Fastx-Toolkit (http://hannonlab.cshl.edu/fastx_toolkit/). Sequences with phread value <20 in 70% of the bases of the sequence - were discarded. Filtered paired-end sequences were assembled with Trinity software^60^. Transcriptome sequenced on GS FLX Titanium platform (Roche) was assembled with Newbler software^61^.

### Gene and transposable element prediction

Transposable Elements (TEs) were defined into different subgroups based on criteria specified earlier^62^. For TE predictions, a *de novo* library was constructed using Repeatmodeler pipeline (http://www.repeatmasker.org/RepeatModeler/). Initial repeat database was generated using RECON^63^, RepeatScout^64^, Tandem Repeat Finder^65^, and NSEG (ftp://ftp.ncbi.nih.gov/pub/seg/nseg/) programs. LTRs were further identified using LTRfinder^66^ program. A reference library for the consensus repeats was developed and merged with the known repbase database of *A. thaliana*. This TE library was used to predict TE in the assembled genome using RepeatMasker (http://www.repeatmasker.org).

The repeat-masked DNA sequences were used to annotate the protein-coding genes using Augustus program^67^. A total of 543 randomly selected full-length CDS sequences were used as a training dataset for Augustus *ab-initio* prediction. The predicted genes were further validated by UniProt database (https://uniprot.org). Genes with <50 bp coding length and TE related genes were removed. Predicted genes were validated by mapping these with previous transcriptome analysis datasets of *B. juncea* using STAR aligner. Further, Iso-seq data was used to validate the predicted genes and to find the alternative transcript isoforms. Iso-seq data was mapped on the assembled pseudochromosomes with Minimap2^68^ program.

### Centromere region specific repeats prediction

Potential centromeric regions were identified from the correlation plots between GBS markers on the genetic maps and physical position of their respective tags on the pseudochromosomes. Python-based scripts were developed to identify kmers of 100bp length with >20 times occurrence in the predicted centromeric regions. The identified kmers were assembled using DNAstar (https://www.dnastar.com/) to get consensus contigs. The assembled contigs were further analysed for their presence in the genome assembly by Blast search analysis, and centromere-specific sequences were selected for both-A and B genomes separately.

### Syntenic block construction and determination of gene fractionation patterns

Syntenic gene blocks were constructed with MCScanX^69^ program. The all-against-all Blastp comparison was carried out against A and B genomes of *B. juncea* with At. MCScanX was run with the default parameters – Match_score: 50, match_size: 5, gap_penalty: −1, e-value: 1e-05, max_gaps: 25. The output was validated with the “synfind” program at CoGE website (https://genomevolution.org/coge). At least three syntenic regions were identified both in the A and B genomes for each gene block of *A. thaliana*. Gene retention pattern for each of the syntenic region was calculated as a proportion of genes retained for 500 flanking genes around a given gene locus considering *A. thaliana* syntenic region as 100% retained. Based on the gene retention pattern of the syntenic regions, the gene blocks were grouped into LF, MF1, and MF2 as defined in *B. rapa*^2^.

For ortholog identification, a two-step gene identification pipeline was implemented. In the first step, protein sequences from A and B genomes of *B. juncea* were compared to *B. oleracea* (C genome) and At using Orthofinder software^22^. All-against-all sequence comparison was carried out using Diamond software^70^. Orthologous groups were determined using the B genome specific genes which did not cluster with the genes of other analyzed species.

### Phylogenetic analysis of gene families

1,482 genes of At, that were present as single copy gene in At (representing all the ancestral gene blocks; A–X) and three orthologs each in the A, B and C genomes were selected for phylogenetic analysis. The amino acid sequences of orthologs were aligned using MUSCLE (v3.8.31)^71^. Poorly aligned regions were trimmed using GBLOCKS^72^ (v0.91) and subsequently with PAL2NAL script^73^. Concatenation of the sequences and conversion to Phylip format was performed using in-house developed Perl scripts. Norwik trees were development and Ka/Ks, and omega values were obtained using PAML package^26^. Divergence time was calculated assuming a mutation rate of 1.5×10^−8^ substitutions per synonymous site per year^74^.

Significance of variation between mean Ks values for orthologue divergence (AT–A, AT–B and AT–C genomes), paralog divergence (within A, B and C genomes) and homoeolog divergence (between A, B and C genomes) were analysed statistically by one-way ANOVA and Tukey’s post hoc test; p<0.05 was considered to be statistically significant.

## Acknowledgements

The work was supported by the Department of Biotechnology (DBT), Government of India through a Centre of Excellence (Grant no.-BT/01/COE/08/06-II) and DBT-UDSC Partnership Centre on Genetic Manipulation of Brassicas (Grant no.-BT/01/NDDB/UDSC/2016). Travel of KP to Arizona Genome Centre was supported by the National Dairy Development Board. We thank Alex Hastie, Bionano Genomics, for carrying out the optical mapping experiments at their centre.

## Authors contributions

KP, JZ, DK, DC and RW carried out the genome assembly; VBR generated optical maps; KP, NB carried out transcriptome analysis; SKY, KP, PS, LB, VG did the mapping work; AM developed DH lines; KP did gene annotation and the other bioinformatic analysis; KP and DP wrote the manuscript, RW, AKP and DP supervised the study. All the authors read and approved the final manuscript.

## Competing Interests

The authors declare no competing interests.

## Data availability

The genome sequence reads of *B. juncea* has been deposited under NCBI bioproject PRJNA550308. *B. nigra* genome and transcriptome sequences have been deposited under bioproject PRJNA324621. The *B. juncea* transcriptome data has been deposited under bioproject PRJNA245462.

