## Supplementary Files for "A chromosome-scale assembly of allotetraploid *Brassica juncea* (AABB) elucidates comparative architecture of the A and B genomes"

**Supplementary File 1 | *Brassica juncea* Varuna transcriptome analysis**

Transcriptome analysis was carried out to use full-length genes as a training dataset for Augustus gene prediction software and to validate the genes which were predicted by the Augustus software. For generating full-length transcriptome sequences, PacBio based sequencing (Iso-seq) was carried out. For validating the predicted genes illumina based short-reads from different previous studies were mapped on the assembled chromosomes.

RNA was isolated from the pooled sample of seedling, leaf, inflorescence with developing siliqua and seed. The pooled RNA sample was considered to represent most of the expressed genes in *B. juncea* plant. Two different transcriptome libraries, one of size range 0.5-1 kb and the other one of 1-2 kb were sequenced separately using P6-C4 chemistry on PacBio RSII system. A total of 551,943 reads on inserts (ROIs) were generated ­– 182,848 in 0.5-1 kb library and 369,095 in 1-2 kb size library. After filtering the sequences for the presence of 5´ adapter reads, poly A reads, chimeric reads, 114,068 and 176,916 full length non-chimeric reads were obtained for 0.5-1 kb and 1-2 kb libraries, respectively.

Finally, clustering and polishing yielded 42,618 and 70,604 consensuses isoforms were obtained of which 27436 and 43366 were high quality isofroms obtained for 0.5-1kb and 1-2kb size fractions, respectively (Supplementary File 1, Table 1).

**SF1-Table1 | Statistics of PacBio based transcriptome sequencing of *B. juncea* Varuna**

| **Classification of reads of Insert 0.5-1 kb 1-2 kb** | | |
| --- | --- | --- |
| Number of reads of insert | 182848 | 369095 |
| Number of five prime reads | 132278 | 243413 |
| Number of three prime reads | 138522 | 245774 |
| Number of poly-A reads | 135760 | 232429 |
| Number of filtered short reads | 19226 | 12394 |
| Number of non-full-length reads | 48948 | 176402 |
| Number of full-length reads | 114674 | 180299 |
| Number of full-length non-chimeric reads | 114068 | 176916 |
| Average full-length non-chimeric read length | 822 | 1347 |
| **Cluster stats** | | |
| Number of consensus isoforms | 42618 | 70604 |
| Number of polished high-quality isoforms | 27436 | 43366 |
| Number of polished low-quality isoforms | 15182 | 27238 |
| Average consensus isoforms read length | 873 | 1459 |

The obtained consensus isoforms were mapped on the assembled *B. juncea* genome sequences using GMAP software. Out of 113,222 consensus isoforms, 104,353 were mapped on the reference genome, representing 35,423 genes in the transcriptome dataset.

In addition to the PacBio based transcriptome sequencing, illumina based transcriptome datasets from three of the previous studies on *B. juncea* were mapped on the assembled genome sequence (SF1 Table2). Short read PE sequences from various studies- SRR1718914, SRR1718917, SRR1718918^1^, SRR1822192, SRR1822193^2^, SRR1269499^3^ were mapped on the assembled genome using STAR aligner^58^ . Unique reads mapped on each gene was calculated and genes with more than 10 mapped reads were designated as expressed.

**SF1 Table 2 | Details of previous transcriptome studies on *Brassica juncea***

|  | SRR number | Tissues | Conditions | Brassica variety | Genes represented | Reference |
| --- | --- | --- | --- | --- | --- | --- |
| 1 | SRR1269499 | Seedling, stem, leaf, pod, developing inflorescence | Field grown | *B. juncea* var Varuna | 74,125 | Paritosh et al, 2014^17^ |
| 2 | SRR1718914 | Seedling | Normal growth | *B. juncea* var Varuna | 58,867 | Bhardwaj et al 2015^20^ |
| 3 | SRR1718916 | Seedling | Temperature stress | *B. juncea* var Varuna | 55,231 | Bhardwaj et al 2015^20^ |
| 4 | SRR1718918 | Seedling | Drought stress | *B. juncea* var Varuna | 54,822 | Bhardwaj et al 2015^20^ |
| 5 | SRR1822192 | Seedling | Normal growth | *B. juncea* var CS52 | 55,886 | Sharma et al, 2015^19^ |
| 6 | SRR1822193 | Seedling | Salt stress | *B. juncea* var CS52 | 54,976 | Sharma et al, 2015^19^ |

A total of 82,008 genes were found to be represented in the transcriptome datasets, of which, 43,776 genes belonged to A genome whereas 38,232 genes belonged to B genome.

**Supplementary File 2 | Transcriptome assembly of *Brassica nigra* variety Sangam**

*Brassica nigra* DH line BnSDH-1 was used for RNA seq of the species. RNA was isolated from seedling and young inflorescence tissues using process as mentioned in the Method section.

Paired-end libraries (2x101bp) were sequenced using Genome Analyzer IIx sequencer (Illumina), a total of 117,766,948 raw sequence reads were produced. These sequences were filtered to remove low-quality reads, which had phread value <20 in 70% of the bases of the sequence. Further, 31 bases were trimmed from the 3′ end of each of the paired-end reads as the region invariably had a phread value of <25. This resulted in 73,864,802 high quality paired-end sequences of 70 bp length for *B. nigra* (SF2 Table 1). Trinity based assembly of the filtered paired end sequences generated 75,087 contigs with the longest read of 8,525bp and N50 value of 1333bp (Table File 2 Table1).

**SF2 Table1 | Statistics of Illumina based paired-end (PE) transcriptome reads**

| Library Type / Insert size | 250bp |
| --- | --- |
| Total number of reads | 117,766,948 |
| Adapter trimmed reads | 115,207,472 |
| Quality filtered (Q<30) reads | 100,115,158 |
| Base trimmed paired- end reads | 73,864,802 |
| Clean pre-processed reads | 73,864,802 |
| % GC | 45 |
| Data in Mb | **1074.1** |

Additionally, a total of 152,075 reads were generated with Roche 454 (GS FLX titanium) based sequencing of *B. nigra* with N50 value of 108bp. Newbler based assembly of the raw reads was carried out with 100bp overlap length with >90% identity. This yielded 25,798 isotigs with N50 value of 1244bp (SF2 Table 2).

While illumina based sequencing validated 35,556 of the 46,227 genes predicted in the genome assembly of *B. nigra* Sangam, 454 based sequencing could identify only 21,964 genes. Both the assemblies taken together validated 35,819 of the predicted genes.

**SF2 Table 2 | Statistics of the transcriptome assembly of *B. nigra* Sangam**

|  | **Roche sequencing**  **Newbler assembly** | **Illumina sequencing Trinity assembly** |
| --- | --- | --- |
| Number of contigs | 25,798 | 75,087 |
| Total size of contigs | 26,752,765 | 67,340,324 |
| Longest contig | 6,108 | 8,525 |
| Shortest contig | 77 | 224 |
| Number of contigs > 1K nt | 10,664 | 24,758 |
| Mean contig size | 1,037 | 897 |
| Median contig size | 871 | 632 |
| N50 contig length | 1,244 | 1,333 |
| L50 contig count | 7,372 | 16,540 |
| contig %A | 27.04 | 28.34 |
| contig %C | 23.18 | 21.52 |
| contig %G | 22.36 | 21.91 |
| contig %T | 27.42 | 28.23 |
| contig %N | 0 | 0 |
| contig %non-ACGTN | 0 | 0 |
