## Supplementary Tables for "A chromosome-scale assembly of allotetraploid *Brassica juncea* (AABB) elucidates comparative architecture of the A and B genomes"

**Supplementary Table 1** | ***Brassica juncea* variety Varuna raw sequencing data obtained with PacBio Sequel platform**

| **PacBio raw reads** | **stats** |
| --- | --- |
| Number of reads | 9,735,857 |
| Total size of reads (Mb) | 92,985.412 |
| Longest read | 89,848 |
| Number of reads > 1K nt | 9,158,458 |
| Number of reads > 10K nt | 36,685,68 |
| Number of reads > 100K nt | 0 |
| Mean read size | 9,551 bp |
| Median read size | 7,108 bp |
| N50 read length | 15,550 bp |
| L50 read count | 2,017,018 |

**Supplementary Table 2 | Statistics of paired-end (PE) reads obtained with Illumina based sequencing of *B. juncea* variety Varuna**

| **Illumina raw reads** | **stats** |
| --- | --- |
| Library Type / Insert size | 250bp |
| Total number of reads | 373,574,346 |
| Base trimmed paired- end reads |  |
| Clean pre-processed reads | 168,456,082 |
| % GC | 41.82 |
| % Q>30 | 98.65 |
| Data in Mb | 33,691.2 |

PE sequence library of insert size 250bp was developed using illumina TruSeq DNA HT Library Preparation kit. Sequencing was carried out on Illumina HiSeq1000 machine as 2x100bp sequences. Raw reads were adapter-trimmed, quality-filtered and sequences were used for mapping on the reference genome.

**Supplementary Table 3 | *Brassica nigra* variety Sangam sequencing data obtained with (a) Illumina paired-end, (b) mate-pair libraries, and (c) Nanopore sequencing raw data- filtering, trimming and quality statistics**

(a) **Illumina paired-end data**

| **Insert size** | **250bp** | **350bp** | **450bp** |
| --- | --- | --- | --- |
| Total number of reads | 129,869,920 | 127,538,609 | 119,911,370 |
| Adapter trimmed reads | 129,863,542 | 127,525,255 | 119,897,571 |
| Quality filtered (Q<30) reads | 116,729,076 | 114,929,773 | 104,455,209 |
| Duplicate reads removed reads | 109,758,727 | 107,021,962 | 98,730,822 |
| Base trimmed paired- end reads | 109,758,727 | 107,021,962 | 98,730,822 |
| Clean pre-processed reads | 108,574,492 | 105,969,982 | 98,100,726 |
| % GC | 38.94 | 41.09 | 40.99 |
| % Q>30 | 98.62 | 98.62 | 97.88 |
| Data in Mb | **19,694.36** | **19,479.45** | **17,639.32** |

(b) **Illumina mate-pair read data**

| **Insert size** | **2Kb** | **6Kb** | **10Kb** |
| --- | --- | --- | --- |
| Total Number of paired-end reads | 16,151,632 | 11,921,106 | 9,422,883 |
| NXtrim paired-end reads | 9,535,592 | 7,163,514 | 5,678,863 |
| Adapter Trimmed reads | 9,495,095 | 7,142,610 | 5,663,810 |
| Quality filtered (Q<30) reads | 5,071,276 | 3,520,150 | 2,570,429 |
| % GC | 40.60 | 39.32 | 38.21 |
| % Q>30 | 96.69 | 96.33 | 95.86 |
| Data in Mb | **1,255** | **908.68** | **643.39** |

**(c) Nanopore based sequencing raw data - filtering, trimming and quality statistics**

| **Nanopore raw reads** | **stats** |
| --- | --- |
| Number of reads | 1,442,476 |
| Total size of reads | 8,096.1 Mb |
| Longest read | 164,545 bp |
| Number of reads > 500 nt | 1,326,870 (92.0%) |
| Number of reads > 1K nt | 1,2390,79 (85.9%) |
| Number of reads > 10K nt | 249,442 (17.3%) |
| Number of reads > 100K nt | 34 |
| Mean read size | 5,613 bp |
| Median read size | 4,078 bp |
| N50 read length | 9,276 bp |
| L50 read count | 290,681 |
| NG50 read length difference | 9,276 bp |

**Supplementary Table 4 | Genome assembly statistics of *B. nigra* variety Sangam**

|  | *B. nigra* (BB, 2n=8) | | |
| --- | --- | --- | --- |
| Illumina | ✓ | ✓ | ✓ |
| MinION |  | ✓ | ✓ |
| Linkage map |  |  | ✓ |
| Total assembly size(bp) | 485,391,109 | - | - |
| Number of contigs | 57,941 | - | - |
| Longest contig (bp) | 253,277 | - | - |
| N50 contig length (bp) | 34,618 | - | - |
| Number of Scaffolds | - | 18,996 | 8 |
| Longest Scaffold(bp) | - | 1,154,440 | 51,978,192 |
| N50 scaffolds length(bp) | - | 85,890 | 43,233,984 |
| Scaffolds assigned to LGs | - | - | 14,147 |
| Unassigned contigs |  |  | - |
| Unassigned scaffolds |  |  | 4,849 |
| Length of assigned sequences(bp) | - | 470,829,086 | 346,918,646 |
| Length of Unassigned sequences(bp) | - | - | 122,832,840 |
| Missing bases (%) | 27.22 | 15.43 | 15.49 |

**Supplementary** **Table 5 | Characteristics of the linkage map of *Brassica juncea* developed using Varuna x Heera (VH) F_1_DH mapping population**

| **LG*** | **Length (cM)** | **No. of markers** | **Anchor * * markers** | **GBS *** markers** | **ave. spacing (cM)** | **max. spacing (cM)** |
| --- | --- | --- | --- | --- | --- | --- |
| Bju-A01 | 205.1 | 202 | 52 | 150 | 1 | 13.6 |
| Bju-A02 | 162.5 | 311 | 43 | 268 | 0.5 | 9.3 |
| Bju-A03 | 283.6 | 366 | 85 | 281 | 0.8 | 17.6 |
| Bju-A04 | 111.1 | 147 | 36 | 111 | 0.8 | 8.8 |
| Bju-A05 | 158.2 | 175 | 34 | 141 | 0.9 | 8.9 |
| Bju-A06 | 137.5 | 116 | 40 | 76 | 1.2 | 10.0 |
| Bju-A07 | 118.4 | 182 | 43 | 139 | 0.7 | 10.1 |
| Bju-A08 | 125.5 | 205 | 36 | 169 | 0.6 | 8.9 |
| Bju-A09 | 332.4 | 353 | 84 | 269 | 0.9 | 14.2 |
| Bju-A10 | 120.2 | 101 | 26 | 75 | 1.2 | 9.9 |
| Bju-B01 | 163.0 | 191 | 31 | 160 | 0.9 | 12.1 |
| Bju-B02 | 207.2 | 289 | 49 | 240 | 0.7 | 9.4 |
| Bju-B03 | 221.4 | 329 | 54 | 275 | 0.7 | 12.2 |
| Bju-B04 | 178.3 | 131 | 43 | 88 | 1.4 | 18.9 |
| Bju-B05 | 88.6 | 63 | 12 | 51 | 1.4 | 12.3 |
| Bju-B06 | 175.0 | 136 | 39 | 97 | 1.3 | 13.8 |
| Bju-B07 | 181.0 | 196 | 50 | 146 | 0.9 | 20.0 |
| Bju-B08 | 263.3 | 287 | 76 | 211 | 0.9 | 14.6 |
| **Overall** | **3,232.3** | **3,780** | **833** | **2,947** | **0.9** | **-** |

* LG nomenclature after Paritosh et al^17^

** Anchor markers consisted of 833 intron-polymorphism (IP), SSR and SNP markers

*** GBS markers were developed using DNA restricted with *Sph*I-*Mluc*I enzymes

**Supplementary** **Table 6 | Characteristics of the linkage map of *Brassica juncea* developed using Tumida x Varuna (TuV) F_1_DH mapping population**

| **LG*** | **Length (cM)** | **No. of markers** | **Anchor** markers** | **GBS*** markers** | **ave. spacing (cM)** | **max. spacing (cM)** |
| --- | --- | --- | --- | --- | --- | --- |
| Bju-A01 | 291.7 | 424 | 34 | 390 | 0.7 | 8.6 |
| Bju-A02 | 131.9 | 984 | 33 | 951 | 0.1 | 7.8 |
| Bju-A03 | 149.5 | 427 | 39 | 388 | 0.4 | 9.3 |
| Bju-A04 | 184.6 | 347 | 19 | 328 | 0.5 | 7.2 |
| Bju-A05 | 130.0 | 579 | 23 | 556 | 0.2 | 4.9 |
| Bju-A06 | 167.1 | 663 | 31 | 632 | 0.3 | 6.3 |
| Bju-A07 | 96.9 | 342 | 24 | 318 | 0.3 | 17.0 |
| Bju-A08 | 107.8 | 478 | 27 | 451 | 0.2 | 4.2 |
| Bju-A09 | 226.4 | 583 | 33 | 550 | 0.4 | 9.0 |
| Bju-A10 | 129.2 | 298 | 26 | 272 | 0.4 | 9.8 |
| Bju-B01 | 129.9 | 490 | 31 | 459 | 0.3 | 7.3 |
| Bju-B02 | 207.5 | 334 | 34 | 300 | 0.6 | 14.1 |
| Bju-B03 | 223.7 | 898 | 54 | 844 | 0.2 | 8.0 |
| Bju-B04 | 268.8 | 436 | 28 | 408 | 0.6 | 11.5 |
| Bju-B05 | 221.2 | 422 | 14 | 408 | 0.5 | 7.9 |
| Bju-B06 | 185.5 | 394 | 22 | 372 | 0.5 | 7.7 |
| Bju-B07 | 255.6 | 432 | 23 | 409 | 0.6 | 12.2 |
| Bju-B08 | 310.1 | 510 | 29 | 481 | 0.6 | 10.1 |
| **Overall** | **3,417.4** | **9,041** | **524** | **8,517** | **0.4** | **-** |

* LG nomenclature after Paritosh et al^17^

** Anchor markers consisted of 524 intron-polymorphism (IP), SSR and SNP markers

*** GBS markers were developed using DNA restricted with *Hin*fI-*Hpy*CH4IV enzymes

**Supplementary** **Table 7 | Features of the linkage map of *Brassica nigra* developed using Sangam x IC2782 F_1_DH mapping population**

| **LG*** | **Length (cM)** | **No. of Markers** | **Anchor** Markers** | **GBS*** Markers** | **ave. spacing (cM)** | **max. spacing (cM)** |
| --- | --- | --- | --- | --- | --- | --- |
| Bni-B01 | 125.8 | 379 | 23 | 356 | 0.3 | 7.7 |
| Bni-B02 | 159.3 | 456 | 36 | 420 | 0.4 | 6.6 |
| Bni-B03 | 206.7 | 365 | 25 | 340 | 0.6 | 9.9 |
| Bni-B04 | 111.6 | 287 | 24 | 263 | 0.4 | 15.7 |
| Bni-B05 | 109.8 | 252 | 25 | 227 | 0.4 | 7.0 |
| Bni-B06 | 130.3 | 364 | 12 | 352 | 0.4 | 6.6 |
| Bni-B07 | 114.9 | 341 | 23 | 318 | 0.3 | 11.0 |
| Bni-B08 | 250.5 | 487 | 40 | 447 | 0.5 | 16.9 |
| **Overall** | **1,208.9** | **2,931** | **208** | **2,723** | **0.4** | **-** |

* LG nomenclature after Paritosh et al^17^

** Anchor markers consisted of 208 intron-polymorphism (IP), SSR and SNP markers

*** GBS markers were developed using DNA restricted with *Sph*I-*Mluc*I enzymes

**Supplementary Table 8** | **Mapping data generated at each step of the hierarchical optical mapping analysis of *Brassica juncea* Varuna genome**

| **Enzyme** | **No. of maps** | **Average map length**  **(mb)** | **Coverage (mol > 150 kb) *** | **Assembly size (Gb)** | **N50 value (mb)** | **Label Density (/100kb)** |
| --- | --- | --- | --- | --- | --- | --- |
| *BssS*I | 1,973 | 0.62 | 501X (436 Gb) | 1.21 | 0.78 | 16.52/100kb |
| *BspQ*I | 1,342 | 0.94 | 400X (348 Gb) | 1.27 | 1.43 | 16.38/100kb |
| DLE-1 | 239 | 3.68 | 219X (190 Gb) | 0.879 | 12.40 | 10.53/100kb |

* Coverage was calculated based on the total size of the initial Canu assembly of 1253 contigs covering ~870Mb of the genome

**Supplementary Table 9** | **Position of the assembled scaffolds and contigs on each pseudochromosome of *Brassica juncea***

| S. No | Chromosome | Number of scaffolds/  contigs | Scaffolds name | Length (bp) | Features |
| --- | --- | --- | --- | --- | --- |
| 1 | BjuA01 | 10 | Scaffold_22 | 17,841,736 | Telomere |
|  |  |  | Scaffold_18 | 8,749,784 |  |
|  |  |  | Scaffold_62 | 9,267,426 | Centromere |
|  |  |  | tig00007318 | 125,523 |  |
|  |  |  | Scaffold_29 | 4,374,888 |  |
|  |  |  | Scaffold_100000423 | 162,856 |  |
|  |  |  | Scaffold_58 | 751,562 |  |
|  |  |  | Scaffold_26 | 2,305,339 |  |
|  |  |  | tig00002913 | 95,740 |  |
|  |  |  | tig00007563 | 129,123 |  |
| 2 | BjuA02 | 2 | Scaffold_48 | 19,836,190 | Centromere |
|  |  |  | Scaffold_63 | 12,820,079 | Telomere |
| 3 | BjuA03 | 1 | Scaffold_33 | 33,612,814 | Telomere, Centro |
| 4 | BjuA04 | 3 | Scaffold_69 | 7,211,206 | Telomere |
|  |  |  | Scaffold_21 | 15,664,184 | Centromere |
|  |  |  | Scaffold_59 | 234,554 |  |
| 5 | BjuA05 | 8 | Scaffold_13 | 15,836,498 | Centromere |
|  |  |  | Scaffold_7 | 15,624,972 |  |
|  |  |  | Scaffold_35 | 2,107,322 |  |
|  |  |  | Scaffold_100000619 | 158,038 |  |
|  |  |  | tig00006685_subseq_424042 - 506315^#^ | 82,273 |  |
|  |  |  | tig00002016 | 114,705 |  |
|  |  |  | Scaffold_1000002285 | 271,597 |  |
|  |  |  | tig00007291 | 130,749 |  |
| 6 | BjuA06 | 6 | tig00001484 | 123,751 |  |
|  |  |  | tig00001405 | 81,929 |  |
|  |  |  | Scaffold_100000345 | 128,387 |  |
|  |  |  | Scaffold_9 | 26,284,396 |  |
|  |  |  | Scaffold_24 | 10,688,200 | Centromere |
|  |  |  | Scaffold_39 | 2,983,050 | Telomere |
| 7 | BjuA07 | 3 | Scaffold_42 | 5,734,093 |  |
|  |  |  | Scaffold_60 | 823,447 | Telomere |
|  |  |  | Scaffold_23 | 22,346,303 | Centromere |
| 8 | BjuA08 | 3 | Scaffold_50 | 11,909,107 | Telomere |
|  |  |  | Scaffold_51 | 12,285,598 | Centromere |
|  |  |  | Scaffold_71 | 4,400,548 | Telomere |
| 9 | BjuA09 | 7 | Scaffold_49 | 356,474 | Telomere |
|  |  |  | tig00002419 | 173,991 |  |
|  |  |  | Scaffold_54 | 923,404 |  |
|  |  |  | Scaffold_19 | 4,927,525 |  |
|  |  |  | Scaffold_100146 | 13,965,245 | Centromere |
|  |  |  | Scaffold_16 | 4,311,000 |  |
|  |  |  | Scaffold_15 | 22,733,440 |  |
| 10 | BjuA10 | 3 | Scaffold_72 | 4,146,020 | Telomere |
|  |  |  | Scaffold_66 | 2,390,740 | Centromere |
|  |  |  | Scaffold_28 | 14,259,645 | Telomere |
| 11 | BjuB01 | 3 | Scaffold_100016 | 142,875 |  |
|  |  |  | Scaffold_55 | 1,800,428 |  |
|  |  |  | Scaffold_5 | 57,124,291 | Centromere |
| 12 | BjuB02 | 2 | Scaffold_3 | 49,309,900 | Telomere, Centromere |
|  |  |  | Scaffold_100186 | 17,950,225 | Telomere |
| 13 | BjuB03 | 5 | Scaffold_10 | 64,311,642 | Telomere, Centromere |
|  |  |  | Scaffold_27 | 1,531,659 |  |
|  |  |  | Scaffold_85 | 350,106 |  |
|  |  |  | Scaffold_43 | 656,597 |  |
|  |  |  | Scaffold_40 | 551,754 |  |
| 14 | BjuB04 | 1 | Scaffold_6 | 66,058,804 | Telomere, Centromere |
| 15 | BjuB05 | 3 | Scaffold_11 | 30,210,523 | Centromere |
|  |  |  | Scaffold_12 | 26,242,956 |  |
|  |  |  | Scaffold_100216 | 150,304 |  |
| 16 | BjuB06 | 2 | Scaffold_1 | 38,710,125 | Telomere, Centromere |
|  |  |  | Scaffold_14 | 20,330,672 |  |
| 17 | BjuB07 | 1 | Scaffold_4 | 58,609,396 | Telomere, Centromere |
| 18 | BjuB08 | 2 | Scaffold_8 | 72,092,522 | Telomere, Centromere |
|  |  |  | Scaffold_32 | 1759191 |  |
|  | Not Placed |  | Scaffold_100012 | 157,000 |  |
|  |  |  | Scaffold_100030 | 1,053,651 |  |
|  |  |  | Scaffold_100209 | 182,602 |  |
|  |  |  | Scaffold_70 (A01) * | 766,743 |  |
|  |  |  | Scaffold_2 (A02)** | 7,491,310 |  |

*****Scaffold 70 was found to belong to pseudochromosome A01 though the GBS markers in the TuV population linkage map

**Scaffold 2 was found to belong to pseudochromosome A02 though the GBS markers in the TuV population linkage map

^#^Subseq- partial sequence

**Supplementary Table 10 | A comparison of the features of the assembled *B. juncea* Varuna genome with previously reported genome of *B. juncea* Tumida (V1.1 and V1.5)**

|  | ***B. juncea* var Varuna** | | | | ***B. juncea* var Tumida (V1.1)** | | | ***B. juncea* var Tumida (V1.5)** | | |
| --- | --- | --- | --- | --- | --- | --- | --- | --- | --- | --- |
| Sub-genome | Chromosome id | Chromosome size (Mb) | Gaps (%) | Anchored percent (%) | Chromosome size (Mb) | Gaps (%) | Anchored percent (%) | Chromosome size (Mb) | Gaps (%) | Anchored percent (%) |
| **AA**  402.1 Mb | Bju-A01 | 44.57 | 12.73 | 10.89 | 45.32 | 26.62 | 11.27 | 38.84 | 21.84 | 10.54 |
|  | Bju-A02 | 40.14 | 0.59 | 8.12 | 25.47 | 11.8 | 6.33 | 33.16 | 10.06 | 9.00 |
|  | Bju-A03 | 33.61 | 0.02 | 8.36 | 43.96 | 8.48 | 10.93 | 40.70 | 8.49 | 11.04 |
|  | Bju-A04 | 23.11 | 2.78 | 5.75 | 33.18 | 11.31 | 8.25 | 25.57 | 14.84 | 6.94 |
|  | Bju-A05 | 34.33 | 4.70 | 8.54 | 38.84 | 23.74 | 9.66 | 28.36 | 11.28 | 7.69 |
|  | Bju-A06 | 40.29 | 17.10 | 10.02 | 38.05 | 12.63 | 9.46 | 29.21 | 5.91 | 7.92 |
|  | Bju-A07 | 28.90 | 1.08 | 7.19 | 28.55 | 18.69 | 7.1 | 30.95 | 15.50 | 8.39 |
|  | Bju-A08 | 28.60 | 8.78 | 7.11 | 29.35 | 9.15 | 7.3 | 25.00 | 10.41 | 6.78 |
|  | Bju-A09 | 47.39 | 2.14 | 11.79 | 62.78 | 17.48 | 15.61 | 52.97 | 19.47 | 14.37 |
|  | Bju-A10 | 20.80 | 6.88 | 5.17 | 22.35 | 6.53 | 5.56 | 18.38 | 4.69 | 4.99 |
|  | **Total** | 341.75 | 5.72 | 82.93 | **367.85** | 15.50 | 91.48 | **323.15** | 13.17 | 87.66 |
| **BB**  547.5 Mb | Bju-B01 | 59.07 | 1.72 | 10.79 | 32.47 | 26.12 | 5.93 | 39.10 | 17.53 | 7.04 |
|  | Bju-B02 | 67.26 | 0.66 | 12.28 | 60.37 | 22.17 | 11.03 | 52.65 | 25.23 | 9.49 |
|  | Bju-B03 | 67.40 | 1.92 | 12.31 | 83.57 | 36.79 | 15.26 | 64.58 | 28.26 | 11.63 |
|  | Bju-B04 | 66.06 | 3.24 | 12.06 | 28.34 | 4.8 | 5.18 | 29.44 | 4.12 | 5.30 |
|  | Bju-B05 | 56.45 | 2.16 | 10.31 | 50.57 | 20.32 | 9.24 | 45.54 | 17.80 | 8.20 |
|  | Bju-B06 | 59.04 | 1.12 | 10.78 | 18.74 | 1.2 | 3.42 | 40.40 | 19.21 | 7.28 |
|  | Bju-B07 | 58.61 | 0.82 | 10.70 | 44.22 | 28.69 | 8.08 | 56.73 | 25.48 | 10.22 |
|  | Bju-B08 | 73.85 | 1.73 | 13.49 | 77.67 | 36.23 | 14.19 | 49.13 | 24.06 | 8.85 |
|  | **Total** | **507.74** | 1.67 | 92.73 | **395.95** | 26.59 | 72.32 | **377.59** | 21.65 | 68.03 |

**Supplementary Table 11 | *Brassica juncea* Varuna genome – distribution of TEs and other repeats on the A and B genomes of *B. juncea***

|  |  | ***B. juncea* A genome** | | | | ***B. juncea* B genome** | | | |
| --- | --- | --- | --- | --- | --- | --- | --- | --- | --- |
|  |  | Repeat Fragment | Intact copies | Length | % genome coverage | Repeat Fragment | Intact copies | Length | % genome coverage |
| **Class I: Retrotransposon** | LTR/Copia | 13138 | 11521 | 14832709 | 4.45 | 37311 | 31188 | 46131465 | 9.09 |
|  | LTR/Gypsy | 25282 | 22209 | 24961078 | 7.48 | 67644 | 59196 | 109400497 | **21.55** |
|  | LTR/Unclassified | 1247 | 1217 | 234093 | 0.07 | 1220 | 1163 | 245257 | 0.05 |
|  | LTR/Cassandra | 937 | 917 | 313982 | 0.09 | 1397 | 1363 | 616357 | 0.12 |
|  | LTR/Caulimovirus | 441 | 393 | 233821 | 0.07 | 879 | 844 | 499124 | 0.10 |
|  | LTR/DIRS | 70 | 69 | 21652 | 0.006 | 234 | 231 | 82745 | 0.02 |
|  | LTR/ERV1 | 43 | 30 | 20513 | 0.006 | 47 | 29 | 20720 | 0.01 |
| **Subtotal** | LTR/Pao | 35 | 31 | 8321 | 0.002 | 177 | 150 | 34968 | 0.01 |
|  | SINE | 124 | 124 | 7283 | 0.002 | 124 | 124 | 7359 | 0.001 |
|  | SINE/tRNA | 4630 | 4590 | 730588 | 0.22 | 3751 | 3724 | 571818 | 0.11 |
|  | LINE/Unclassified | 378 | 366 | 107767 | 0.03 | 207 | 195 | 64845 | 0.01 |
|  | LINE/L1 | 13647 | 12696 | 8317266 | 2.49 | 15391 | 14059 | 9181807 | 1.81 |
|  | LINE/Penelope | 239 | 229 | 25218 | 0.01 | 923 | 871 | 289111 | 0.06 |
|  |  | **60211** | **54392** | **49814291** | **14.94** | **129305** | **113137** | **167146073** | **32.92** |
| **Class II: DNA transposon** | DNA/Unclassified | 5027 | 4677 | 1496636 | 0.45 | 7065 | 6329 | 2030898 | 0.40 |
|  | DNA/CMC-EnSpm | 5751 | 5256 | 4025741 | 1.21 | 12749 | 11624 | 8729040 | 1.72 |
|  | DNA/Crypton-S | 114 | 108 | 30302 | 0.01 | 2058 | 1822 | 752723 | 0.15 |
|  | DNA/hAT | 1010 | 938 | 297987 | 0.09 | 2567 | 2331 | 1255843 | 0.25 |
|  | DNA/hAT-Ac | 14750 | 13456 | 4993697 | 1.50 | 16514 | 15077 | 6001122 | 1.18 |
|  | DNA/hAT-Charlie | 879 | 864 | 571709 | 0.17 | 864 | 857 | 480524 | 0.10 |
|  | DNA/hAT-Tag1 | 4611 | 4263 | 997320 | 0.30 | 5388 | 4960 | 1457492 | 0.29 |
|  | DNA/hAT-Tip100 | 678 | 572 | 246679 | 0.07 | 1014 | 885 | 429855 | 0.09 |
|  | DNA/IS3EU | 784 | 673 | 288397 | 0.09 | 360 | 339 | 127896 | 0.03 |
|  | DNA/Maverick | 839 | 728 | 306089 | 0.09 | 213 | 209 | 60514 | 0.01 |
|  | DNA/Merlin | 35 | 29 | 11675 | 0.01 | 39 | 29 | 11468 | 0.01 |
|  | DNA/MuLE-MuDR | 7674 | 6310 | 4938721 | 1.48 | 14992 | 12173 | 7975759 | 1.57 |
|  | DNA/PIF-Harbinger | 7145 | 6439 | 2187997 | 0.66 | 7171 | 6516 | 2360355 | 0.47 |
|  | DNA/RC | 2990 | 2541 | 1101598 | 0.33 | 4268 | 3377 | 1983887 | 0.39 |
|  | DNA/TcMar-Pogo | 2461 | 2320 | 590679 | 0.18 | 3251 | 2999 | 763910 | 0.15 |
|  | DNA/TcMar-Stowaway | 7688 | 7504 | 1638827 | 0.49 | 7124 | 6979 | 1466747 | 0.29 |
|  | DNA/Zisupton | 1109 | 957 | 298172 | 0.09 | 1502 | 1336 | 476769 | 0.09 |
| **Subtotal** |  | **63545** | **196405** | **67928039** | **20.37** | **87139** | **244000** | **100421863** | **19.78** |
|  | **Simple repeat** | NA | 1713 | 798160 | 0.24 | NA | 1648 | 603530 | 0.12 |
| **Other repeats** | **Unknown** | NA | 126786 | 42405922 | 12.71 | NA | 163894 | 63274593 | 12.46 |
|  | **Satellite** | 0 | 10271 | 701731 | 0.21 | 0 | 616 | 178938 | 0.04 |
| **Grand Total** |  | **123756** | **250797** | **117742330** | **35.30** | **216444** | **357137** | **267567936** | **52.70** |

**Supplementary Table 12 | *Brassica juncea* Varuna genome – centromere-specific repeat sequences identified in the A genome**

| **Chromosome** | **CentBr1 (176 bp)** | **CentBr2 (176 bp)** | **Cent_A_2 centBr3** | **Cent_A_3 centBr4** | **Cent_A_4**  **(TR238)** | **CentBr5** | **Cent_A_6**  **(CRB3)** |
| --- | --- | --- | --- | --- | --- | --- | --- |
| Bju_A01 | 15 | 3288 | 5 | 3 | 93 | 6 | 9 |
| Bju_A02 | 10 | 0 | 0 | 0 | 2 | 0 | 0 |
| Bju_A03 | 0 | 64 | 0 | 1 | 1 | 0 | 0 |
| Bju_A04 | 6 | 0 | 1 | 1 | 1 | 0 | 0 |
| Bju_A05 | 281 | 0 | 9 | 9 | 204 | 8 | 49 |
| Bju_A06 | 76 | 0 | 5 | 5 | 144 | 44 | 26 |
| Bju_A07 | 4 | 0 | 0 | 0 | 1 | 0 | 0 |
| Bju_A08 | 2 | 0 | 10 | 10 | 119 | 14 | 13 |
| Bju_A09 | 13 | 1 | 0 | 0 | 4 | 0 | 0 |
| Bju_A10 | 4 | 0 | 0 | 0 | 8 | 0 | 2 |
| Bju_B01 | 0 | 0 | 10 | 11 | 68 | 14 | 0 |
| Bju_B02 | 0 | 0 | 9 | 14 | 43 | 3 | 0 |
| Bju_B03 | 0 | 0 | 13 | 8 | 61 | 2 | 0 |
| Bju_B04 | 0 | 0 | 7 | 6 | 55 | 4 | 0 |
| Bju_B05 | 0 | 0 | 1 | 6 | 55 | 2 | 0 |
| Bju_B06 | 0 | 0 | 10 | 8 | 47 | 0 | 0 |
| Bju_B07 | 0 | 0 | 4 | 9 | 49 | 3 | 0 |
| Bju_B08 | 0 | 0 | 16 | 7 | 62 | 4 | 0 |

>**centBr1**

AAGCTTGATTTTGATACATAAAGTAATGGAGAATGACYAGGAAGTTGAATAAATCTCATAGGAGTTAGGATGAAGAAGTTATCCAACTTTCAAATCAGGTGATTCCAGATTCCAGTTTGGGAATAGCACAGCTTCTTCGTCGTTCCAATCAAACCAGGATGAATCTCTTTGTAAG

>**centBr2**

AAGCTTGATTTTGATACATAAAGTAATGGAGAATCACTAGGAAGTTGAATAAATCTCATAGGAGTTAGGATGAAGAAGCTATCCCACTTTCAAATCAGGTGATCCCAGTTTTCCTGTTTGGGAATATGACAGCTTCTTTGTCATTCTAATCAAACCAGGATGAATCGCGATGTAA

> **centBr3**

TGGTGTGATCCTGATCAGGATTGGAATCAATGGATTGAAGCCAATGTATGGGTCAGCTATACGACTGATCATGGTGCAGTTTAATAGACAGTATCATACTCTTATTGGCATGATCTATGAGGTTCTGGATAGAGAGAAGAGAAGAACTGTTGGTACTGCCTTGAAACAGCTTGAACAAAGAGGATTCAACCAACTGAGCTTAACAGGAGCTATGGAGACCATTCACCTCATCCAAATTTGTATGGAGAACAAGGAAAACATGTCAGGAAGTTTCTGGATTGGGTTATATTTGGATCTAGGCGAAATAGGGCTTTTTCTGTCCAAAAAGAGAGTTGCAGCCAGAATCTTGATCCACTTGCTTTCTAGCTGGCCTAACAGGACGTGGAGGATCTAAAGTTCATTAGGAGACACTCCAAATCAAGTTCAAGTCGTGGCCCATCATAAACCAAAGAAGAACCAGCTTGTTGGTTTGATTTAATTGCAATGTAATGGGTTTAGTTTATTGGGCTTTTCATTTAATTTAAACCAAAGACAATTAGGAATTTGAGTAAACCAGATTTGGGTAGATTAGTATTTAAAATAAACCAATTTCGAATTGGATTAGGAGATTAAACCATCTTGGTTTAAATTTAGGGTTTTCAGTTTTATTCTTTTAACATATGTAGCCACGTTTTGGCATGAAAATCAAT

> **centBr4**

TTTTTTTCAACTTCTTTGTTTTGAGTTTTCTCTCAATTCTGCTGGTGTGTAATATCCAGCTTGGGTTCTCTGATTTGGTGTGTCATATCCAGATCAGAAGCCTTGGAGTATCAGGAGACGAGCCATCTCTCTTGTGTCCCACATCAATCCACAAACCTTGATACAGATCGAGTCTTTGATTCTCTGTTTGCAAGGATTATCTCTGTGATCCATCCTAGACACTAAGCAGGGTGATCCTGTGTCTGTGGAGCAGAAAAAAA

> **TR238_partial**

ACATCATTAGGACTAAAGATAACATTGCATTGTAAATCATTAGTACATTTCTTCACAGATAATAGGTTTGATGTGAATTCAGGCATGTAAAAAGCTCTAGATTCTTTGTTAAACAACTTAAGCTTTCCTATTCCTCTAATAGGTATCTTATCCCCATTTGCTATCATCACATGCCCATG

> **centBr5**

GAGAGATGGGCAAAGTGGTCCTGGATAGTGACAGGTCCTAACGGGTGGGAAGACTTTGAATCCTTAACCAGACCCGTGACCTGCACTCTCAACGGTTCCCCACTTCCCTTAGGCGATTCCAAACATTCTTCTTGCCTTCTTCTCTTCTACATCAGTCACTCTTA

> **CRB3_partial**

ACAAGAGGCGAAATGGGCACTATTGAAGCATGTAGAGTCAAGGTTGCTTTCCGGATGATTGGATGATAAGGCTCACACGCTTTGCCTCTTTACAGCCTGTTCCAGCCTGGCACGCTCTTCCCTTATCCACTCAACGGTCTGATCCCTTCTCCTTAAG

**Supplementary Table 13 | *Brassica juncea* Varuna genome – centromere-specific repeat sequences identified in the B genome**

| **Chromosome** | **CentBni 1** | **CentBni 2** | **CentBni 3** | **CentBni 4** | **CentBni 5** | **CentBni 6** | **CentBni 7** |
| --- | --- | --- | --- | --- | --- | --- | --- |
| **Bju_A01** | 8 | 0 | 24 | 2 | 41 | 8 | 0 |
| **Bju_A02** | 2 | 0 | 0 | 23 | 13 | 18 | 18 |
| **Bju_A03** | 0 | 0 | 0 | 0 | 1 | 1 | 0 |
| **Bju_A04** | 0 | 0 | 0 | 0 | 0 | 0 | 0 |
| **Bju_A05** | 1 | 0 | 10 | 0 | 0 | 0 | 0 |
| **Bju_A06** | 13 | 0 | 0 | 14 | 7 | 12 | 3 |
| **Bju_A07** | 0 | 0 | 0 | 0 | 1 | 1 | 0 |
| **Bju_A08** | 7 | 1 | 18 | 0 | 0 | 0 | 0 |
| **Bju_A09** | 0 | 0 | 2 | 17 | 13 | 14 | 10 |
| **Bju_A10** | 0 | 0 | 0 | 0 | 0 | 0 | 0 |
| **Bju_B01** | 16 | 23 | 66 | 0 | 0 | 0 | 0 |
| **Bju_B02** | 0 | 5 | 59 | 25 | 38 | 40 | 48 |
| **Bju_B03** | 0 | 17 | 51 | 36 | 15 | 13 | 45 |
| **Bju_B04** | 8 | 77 | 112 | 3 | 20 | 20 | 49 |
| **Bju_B05** | 2 | 18 | 78 | 63 | 28 | 27 | 66 |
| **Bju_B06** | 41 | 16 | 58 | 11 | 22 | 26 | 46 |
| **Bju_B07** | 0 | 43 | 57 | 19 | 45 | 46 | 53 |
| **Bju_B08** | 0 | 41 | 79 | 12 | 16 | 16 | 75 |

**Sequences:**

>Contig_8g **Cent_Bni1**_8

TGTTGACATGAAGAATAAAGCAAGAAGATAGGAACTTGGAGAACAAGGCTGTCCGTACAACCTAAGAGTGACTTGCAAGGAAAGGAAGCAAGTCATGAGTTAACCAACTCGGTCAAGTGATTAAAAGATCTCCTAGTGAGTGGGAATGTCGACATGTATCCGTTGAAGAGAAGTGAGTGGAGGAGTCACGGGTGTGACTAAGAGAAGAGAGAGGGAGCTGACCGTTGGGTGCTGTCACTATCCCTCTTGCTGAACGAGCCATCCGCGGATCTGCCTCCTCTTTGTTTATTCTTTCCCTTGTACTATCTTCTCATT

>Contig_9 **Cent_Bni2**

AGTGTTGAGATGAAGCAATGAAAGACCAAGGAAGATCAAAGGAAGAGCATTGGCGCCAGCAAAGATAATTTGAATCTACCTTGGGAATCAAGGCAATCCGTACAGACTTGGGAACCAAGGAACTAATGTGCAACTAATGTGCCTTATGAAATCAAAGTTGAAGAAGTCCAAGGTGATTAAATAATAGCTGGATGTCGGCAAGGCTGAAGATGAACGTTGGAGTGGTGTGGACCACTTGGTCACATGCTGGTGCAGGCTGAAGAGAGACGTTGGCGTGCTGTCCCTTTCAAGCTCCACTCGCAGCTCTTCCTTCTCTTTGTTTATTCTTTTCCTTGTCTCCTTGTGTCCTGATTTGGTCTTTAAATATTGTTGTATGAGCTACTCTTTTGGCAAGCAATCAACGAGCTCTTTAGTTGCTTCACTCTCTCTCATGAATTCTCTTATCTCTCTCTCTTGTTCTTGCTCGGTTTCCAGCAAATAACTCTCTTAAATCTCTTAATCT

>Contig_12 **Cent_Bni3**

TGTTGGAATGAAGAAGTAAGAAGCTGGAGTTGATGAaGAGACCAAGCATCAAGAGCAAGCAAATCCGGAACAAGCAGATTTGAATCTGTccGTACAGTtttggCTTRSAaGTRCAAgcTAACCTAGAGTGTCTTGGAAGTCAAGGCAAGTGGGRAACTCGTCCAGCTGAGtCTTCTGAAGCAGATTTGAAGTTTAAAAAGGGTATCAGACCGTTGGAGAGAGTTTAGGCAAGTTTGGAATTAAATAGAGTAAAAGTCACATGTGCATAAGAGGATGAGCCTGCTGTCACTTTCACTCAACCAGACTGTGCCCATCTCTCCCTCCTGTCACTCTCACTCCCTCTGGTCGAGATTCACCTTGGCTGCATCCTCCTGGCTCCATTAGAAACCCCACAATAATATTAGTTTAGTAGATTTGTACTTTATGATTTGTTTAATACCTTGCAAGTTGTAATCTGGCTATAAAGCCACCATTGTACACAATTTGTAACCTCAAGCAAAGAGGAGTTCTCTTTCTTCATTTTCTCTGGTTCCAAACTCTCTCAGACTCTCAATCTCTCTATTCTCTCTTGTTACTCACGAAATTACTCTCTTGGATACTCACTGAAACTCTCATACTCTCTTACTCTCAATTGCTTACTCTTTGATTCTCTCTTAGTCTTTTACTCTCTCCATCTCTTACACTCGCCCATACTCTGTTTCTATTACTCTCTCACATCATATTGACTCTTTTCTCA

>Contig_29 **Cent_Bni4**

TTTTCCTTAGAATCAAAGTGTGGCAAGTTACAAGTATTTAAAGACAAAGCTTGGCAAGTTGCCTCACTTCAGAAACAATAAACTAATCTAGGCTAAGTACAGAAACTATTAAACTAATCAAATAGAATAAATAAATGTGTTCTGATCAAGGCATGAAAGTGACATGTCCCAAACGGCTGGATTCAAATTTGGAACATGTGACCTTTCTTATTTAAATCATTCTTCTTGTCTCAGCAGCCTCCAACGGTCTTCTCCTCTCCTAATGTCGACCAAGTCAAGTCTGAGCTTGCCTTGACTCCTAAGGCATGTTAGGTTCAGCCTTGGATGCTTGCTTCACAAGTATTGTACGGAATAGCTTGTGATCCAAGCAAGTGCAGTTCACATGTTCTTCACTCATCCGGATACTTCTTCTGCCTTATCTCTTCTCTTCTTGTCTTCATGT

>Contig_30 **Cent_Bni5**

ACCAACACTGATGAGTCTGAGGGTCAGGAAGTTCCTCACCTTGACCATGAGGGGGGCAACAACAATACTGAGCCAATCCATGATCAAGATGAGCAACAAGATCAAGAGGATCAAGAAGCACCAGTGGTGAATGAGAACCAGACTTTGGCAAATGAAGAGGAGGTTCAACATGAAGAACCACAACCAGTACTAAGGAGGAGTACTAGGATCAAACGACCGGCTTCTAATTGGATCAACACAAGGGTTTACTTCAACAGCCAAGCAGTTGCTCATCCCACTCAAGCAACATGCTCCCTTGCTCAATATCCTATGGATCATCAAGCTTTCACCACTAATCTTGATGAGGCGTACATACCAAGGTCATATGAAGAGGCTATGGCAATCAAGGAATGGAGAGATTCAGTAGGAGATGAGATGGGTGCTATGGAGAAGAATGGAACCTGGTTTGAAACTGAGCTTCCCAAAGGCAAAAAGGCTGTTACTAGCATGCT

>Contig_31 **Cent_Bni6**

CAGGCTACTTGCCTGAAGCATGAAGACAAGTGGGTTCAAGAGGAACATCAAGCACTTGGAATTATACACAGATCTCTGTCTGAACAGATCTTGGAGGATCATTATCATCTCAAAACCGCCAAGAATCTATGGGAGAAGCTTCAAAATGTTTATGGAAATCTATGGGAGAGTCTTCAAGCTGTTTATGAGCAAGACAAAATTTTCAGTCTTCTCCTAAACCTGGATTCACCCTACAATAGTCTGATAAGTCATATCATCCAAGAAAGGAAGCTTCCAAGTTCCCACGGAAGCCAGCCGAGTACACTTAAGCTCAAGGAGTTTGAGCATCAAGTAATTGAAGGAAGAACCATGTATTCTGCATTATGTGGTGATTCTAATGAAGGAGGAACATGGAGGAGTATGGCAGGGGCGACCAATTCTGATGGCCCAGTGACCAGAAGAGACTTTGATTCCTTTATAAGAGCTGCCAAGGCCCTTATCTCACCCAGGGAGGTTGGTAAAAATTCTCCTGACTCTAAACCCATTATCATAGATTCAGGAGCTAG

>Contig_34 **Cent_Bni7**

CCATTGCAAGAAGAATTCAACAAGGACAGCAACAACTTGGTGTTAGAACAGATGGCCACCAGAGACTACTTGACAAGATCAGGTGGAGAGTCAAGCTGAGTGGAAAGCTATGAATCAGCCCAAGGCATGGAGTTAGAACAGTTACCTTCTGGTTGGGAAGCTATGGATCAGCTTCTAAAAAGGAGTTAGAACAGTTACCTTCAGTGGGGAAGCTATGAATCAGCCCATAGCATGTGTGCTTGTACCTGGTTGATGAAGCTTCAAGTTAAGAGGGCTGTTGTCTCAAGGCATGGTGATCATCATTGGCTCTCATCTTCCAAGCTTGACCACATACTCT

**Supplementary Table 14 | *Brassica juncea* Varuna genome – centromere specific transposable elements in the B genome**

| **Chromosome** | **rnd-1_family-218**  **(LTR/Copia)** | **rnd-1_family-306**  **(DNA/Maverick)** | **rnd-1_family-312**  **(Unknown)** | **rnd-1_family-320**  **(DNA/CMC-EnSpm)** | **rnd-1_family-398**  **(Unknown)** | **rnd-1_family-469**  **(Unknown)** | **rnd-1_family-507**  **(Unknown)** | **rnd-6_family-4293**  **(Unknown)** | **rnd-1_family-304**  **(Unknown)** | **rnd-1_family-960**  **(Unknown)** | **rnd-5_family-1821**  **(Unknown)** |
| --- | --- | --- | --- | --- | --- | --- | --- | --- | --- | --- | --- |
| **Bju_A01** | 0 | 0 | 26 | 0 | 0 | 0 | 0 | 20 | 24 | 0 | 0 |
| **Bju_A02** | 0 | 0 | 0 | 0 | 0 | 0 | 0 | 0 | 0 | 0 | 0 |
| **Bju_A03** | 0 | 0 | 0 | 0 | 0 | 0 | 0 | 0 | 0 | 0 | 0 |
| **Bju_A04** | 0 | 0 | 2 | 0 | 0 | 0 | 0 | 0 | 0 | 0 | 0 |
| **Bju_A05** | 9 | 4 | 53 | 4 | 0 | 0 | 2 | 33 | 14 | 1 | 1 |
| **Bju_A06** | 0 | 0 | 36 | 0 | 0 | 0 | 0 | 21 | 0 | 0 | 0 |
| **Bju_A07** | 2 | 0 | 0 | 0 | 0 | 0 | 0 | 0 | 0 | 0 | 0 |
| **Bju_A08** | 9 | 2 | 15 | 2 | 0 | 0 | 1 | 14 | 17 | 0 | 0 |
| **Bju_A09** | 0 | 0 | 0 | 0 | 0 | 0 | 0 | 0 | 0 | 0 | 0 |
| **Bju_A10** | 0 | 0 | 0 | 0 | 0 | 0 | 0 | 1 | 0 | 0 | 0 |
| **Bju_B01** | 10 | 33 | 51 | 34 | 60 | 0 | 9 | 1 | 26 | 14 | 12 |
| **Bju_B02** | 6 | 27 | 61 | 27 | 28 | 0 | 12 | 2 | 37 | 11 | 8 |
| **Bju_B03** | 16 | 27 | 38 | 27 | 26 | 2 | 24 | 1 | 3 | 14 | 9 |
| **Bju_B04** | 5 | 64 | 61 | 64 | 29 | 2 | 22 | 3 | 64 | 17 | 17 |
| **Bju_B05** | 12 | 30 | 28 | 30 | 34 | 7 | 11 | 2 | 11 | 20 | 20 |
| **Bju_B06** | 26 | 34 | 43 | 34 | 49 | 3 | 18 | 3 | 19 | 28 | 21 |
| **Bju_B07** | 36 | 34 | 41 | 35 | 33 | 7 | 0 | 3 | 12 | 24 | 21 |
| **Bju_B08** | 0 | 51 | 72 | 51 | 19 | 0 | 18 | 4 | 29 | 31 | 26 |

**Supplementary Table 15 | A catalogue of the genes predicted on different pseudochromosomes of *Brassica juncea* Varuna and their orthologs in *Arabidopsis thaliana* (along with their respective gene blocks) and *B. rapa* Chiifu, and homoeologs in the A and B genomes.** Column A – gene blocks as identified in *A. thaliana*; Column B – *A. thaliana* orthologs with gene ids; Column C – predicted *B. juncea* genes with the assigned gene id; Column D – physical position of the genes on the pseudochromosomes; Column E – paleogenome to which the gene belongs; Column F – expression status of the predicted *B. juncea* genes. (“Expressed” means that the gene was found in transcriptome analysis in this study or other studies – Supplementary File 1, blank box represents – expression not reported); Column G – homoeolog of the gene described in Column C in the A or the B genome of *B. juncea*.

**Supplementary Table** **16. | Number of unassigned contigs from previous genome assemblies of *B. rapa* variety Chiifu and *B. juncea* variety Tumida that could be placed on the 18 pseudochromosomes of *B. juncea* Varuna assembly**

| **Species** | **Unassigned contigs** | **Assigned to *B. juncea* Varuna** | | **Remaining** |
| --- | --- | --- | --- | --- |
|  |  | **A genome** | **B genome** |  |
| ***B. rapa* V1.5*** | 40,357 | 35,644 | - | 4,663 |
| ***B. juncea* V1.1**** | 1,948 | 525 | 1,277 | 146 |

* *B. rapa* variety Chiifu genome version 1.5 as in BRAD database

** *B. juncea* variety Tumida version 1.1 as in BRAD database

**Supplementary Table** **17** **|** **Block wise number of genes retained in the three constituent paleogenomes of the A and B genomes of *B. juncea*. Blocks are as described in *A. thalina***

| block | LF_A | MF1A | MF2A | LF_B | MF1B | MF2B | LFA vs MFs | MF1A vs MF2A | LFB vs MFBs | MF1B vs MF2B |
| --- | --- | --- | --- | --- | --- | --- | --- | --- | --- | --- |
| A | 1068 | 672 | 605 | 1116 | 738 | 627 | 8.0E-130^*^ | 1.2E-04^*^ | 4.6E-127^*^ | 3.6E-10^*^ |
| B | 708 | 451 | 452 | 720 | 479 | 481 | 2.4E-65^*^ | 9.5E-01^*^ | 3.9E-54^*^ | 9.0E-01 |
| C | 401 | 310 | 259 | 432 | 349 | 275 | 1.5E-24^*^ | 7.4E-06^*^ | 1.3E-25^*^ | 2.8E-10^*^ |
| D | 214 | 134 | 62 | 210 | 162 | 123 | 2.0E-93^*^ | 3.0E-38^*^ | 3.2E-18^*^ | 6.6E-07^*^ |
| E | 835 | 618 | 437 | 874 | 646 | 462 | 2.1E-97^*^ | 1.8E-34^*^ | 2.1E-99^*^ | 9.8E-34^*^ |
| F | 1313 | 1006 | 820 | 1387 | 1026 | 862 | 7.9E-87^*^ | 4.1E-20^*^ | 3.6E-99^*^ | 2.8E-15^*^ |
| G | 20 | 15 | 0 | 24 | 26 | 0 | - | - | - | - |
| H | 260 | 144 | 54 | 326 | 162 | 147 | 3.1E-193^*^ | 3.3E-67^*^ | 1.7E-85^*^ | 8.0E-02 |
| I | 308 | 322 | 83 | 340 | 325 | 107 | 8.5E-135^*^ | 2.8E-301^*^ | 1.7E-112^*^ | 3.4E-195^*^ |
| J | 937 | 696 | 599 | 949 | 584 | 654 | 1.4E-61^*^ | 2.1E-08^*^ | 1.5E-80^*^ | 1.1E-04^*^ |
| K | 104 | 97 | 59 | 109 | 132 | 40 | 3.6E-09^*^ | 2.6E-12^*^ | 1.4E-28^*^ | 4.9E-94^*^ |
| L | 189 | 145 | 97 | 190 | 148 | 82 | 1.1E-23^*^ | 5.5E-12^*^ | 2.1E-35^*^ | 6.5E-25^*^ |
| M | 216 | 131 | 109 | 256 | 122 | 125 | 1.0E-36^*^ | 2.9E-03^*^ | 8.0E-64^*^ | ***7.0E-01*** |
| N | 690 | 437 | 382 | 710 | 454 | 414 | 7.4E-88^*^ | 6.9E-05^*^ | 2.1E-79^*^ | 5.4E-03^*^ |
| O | 234 | 151 | 75 | 244 | 163 | 98 | 3.2E-85^*^ | 2.3E-35^*^ | 5.3E-58^*^ | 1.6E-20^*^ |
| P | 113 | 86 | 50 | 123 | 92 | 67 | 7.0E-21^*^ | 6.0E-13^*^ | 3.8E-14^*^ | 1.6E-05^*^ |
| Q | 291 | 175 | 167 | 287 | 191 | 171 | 1.2E-38^*^ | ***3.8E-01*** | 1.9E-29^*^ | 3.1E-02^*^ |
| R | 1114 | 738 | 760 | 1165 | 768 | 798 | 1.7E-79^*^ | ***2.6E-01*** | 2.5E-83^*^ | ***1.3E-01*** |
| S | 252 | 96 | 66 | 274 | 130 | 44 | 3.8E-171^*^ | 1.8E-07^*^ | 4.2E-298^*^ | 4.3E-75^*^ |
| T | 83 | 136 | 96 | 79 | 135 | 102 | 2.2E-06^*^ | 7.8E-09^*^ | 9.8E-08^*^ | 3.8E-06^*^ |
| U | 1409 | 932 | 696 | 1419 | 934 | 732 | 6.1E-214^*^ | 1.1E-36^*^ | 5.3E-197^*^ | 4.6E-26^*^ |
| V | 283 | 194 | 180 | 305 | 211 | 193 | 1.7E-23^*^ | ***1.4E-01*** | 4.7E-25^*^ | ***6.7E-02*** |
| W | 565 | 443 | 407 | 604 | 444 | 424 | 2.0E-22^*^ | 1.2E-02^*^ | 5.3E-31^*^ | 1.7E-01 |
| X | 458 | 266 | 236 | 476 | 264 | 237 | 1.5E-77^*^ | 5.7E-03^*^ | 1.9E-91^*^ | 1.3E-02^*^ |

Chi square test was performed between the observed and expected number of genes retained. Equal number of genes was expected to be retained in the three sub genomes in each of the gene block. P values <0.05 was considered as significant difference between the observed and expected numbers.

^* significant difference (p <0.05)^

**Supplementary Table 18** **| Divergence analysis between *A. thaliana* and the constituent paleogenomes of the A and B genomes of *B. juncea*, based on synonymous nucleotide substitutions** – (a) Mean Ks values of ortholog, paralog and homeolog comparators (b) One-way ANOVA and Tukey’s post hoc test to find the significant mean differences between the analysed datasets.

(a)

| Ortholog divergence | | Paralog divergence | | Homeolog divergence | |
| --- | --- | --- | --- | --- | --- |
|  | Mean Ks value |  | Mean Ks value |  | Mean Ks value |
| AT vs LF_A_ | 0.50±0.19 | LF_A_-MF1_A_ | 0.45±0.21 | LF_A_-LF_B_ | 0.27±0.12 |
| AT vs MF1_A_ | 0.54 ±0.21 | LF_A_-MF2_A_ | 0.44±0.20 | MF1_A_ -MF1_B_ | 0.27±0.11 |
| AT vs MF2_A_ | 0.53±0.21 | MF1A-MF2_A_ | 0.45±0.21 | MF2_A_-MF2_B_ | 0.25±0.09 |
| AT vs LF_B_ | 0.54±0.23 | LF_B_-MF1_B_ | 0.48±0.25 |  |  |
| AT vs MF1_B_ | 0.54±0.23 | LF_B_-MF2_B_ | 0.46±0.22 |  |  |
| AT vs MF2_B_ | 0.52±0.20 | MF1_B_-MF2_B_ | 0.48±0.24 |  |  |

(b)

|  | diff | lwr | upr | p adj | Sig. |
| --- | --- | --- | --- | --- | --- |
| **A_ genome paralogs** | | | | | |
| LF_A_/ MF2_A_ *vs* LF_A_/ MF1_A_ | -0.008746928 | -0.032157580 | 0.014663724 | 0.8952734 | - |
| LF_A_/ MF2_A_ *vs* LF_A_/ MF2_A_ | -0.005185955 | -0.028596608 | 0.018224697 | 0.9887014 | - |
| MF1_A_/ MF2_A_ *vs* LF_A_/ MF2_A_ | 0.003560972 | -0.019849680 | 0.026971624 | 0.9980656 | - |
| **B_ genome paralogs** | | | | | |
| LF_B_/ MF2_B_ *vs* LF_B_/ MF1_B_ | 0.008281364 | -0.015129288 | 0.031692016 | 0.9153664 | - |
| LF_B_/ MF2_B_ *vs* LF_B_/ MF2_B_ | -0.004859352 | -0.028270004 | 0.018551300 | 0.9916207 | - |
| MF1_B_/ MF2_B_ *vs* LF_B_/ MF2_B_ | 0.003422012 | -0.019988640 | 0.026832664 | 0.9984021 | - |
| **Paralogs vs Homeologs** | | | | | |
| LF_A_/ MF1_A_ *vs* LF_A_/ LF_B_ | 0.18855632 | 0.16387427 | 0.21323837 | 0 | *** |
| LF_A_/ MF2_A_ *vs* LF_A_/ LF_B_ | 0.17794528 | 0.15326323 | 0.20262733 | 0 | *** |
| MF1_A_ / MF2_A_ *vs* LF_A_/ LF_B_ | 0.18388637 | 0.15920432 | 0.20856842 | 0 | *** |
| LF_B_/ MF1_B_ *vs* LF_A_/ LF_B_ | 0.1768369 | 0.15215485 | 0.20151895 | 0 | *** |
| LF_B_/ MF2_B_ *vs* LF_A_/ LF_B_ | 0.17055199 | 0.14586994 | 0.19523404 | 0 | *** |
| MF1_B_/ MF2_B_ *vs* LF_A_/ LF_B_ | 0.17784067 | 0.15315862 | 0.20252272 | 0 | *** |
| LF_A_/ MF1_A_ *vs* MF2_A_ / MF2_B_ | 0.20463186 | 0.18133531 | 0.22792842 | 0 | *** |
| LF_A_/ MF2_A_ *vs* MF2_A_ / MF2_B_ | 0.19168172 | 0.16838517 | 0.21497828 | 0 | *** |
| MF1_A_/ MF2_A_ *vs* MF2_A_ / MF2_B_ | 0.19559706 | 0.1723005 | 0.21889362 | 0 | *** |
| LF_B_/ MF1_B_ *vs* MF2_A_ / MF2_B_ | 0.18738845 | 0.16409189 | 0.210685 | 0 | *** |
| LF_B_ / MF2_B_ *vs* MF2_A_ / MF2_B_ | 0.18038452 | 0.15708797 | 0.20368108 | 0 | *** |
| MF1_B_ / MF2_B_ *vs* MF2_A_ / MF2_B_ | 0.1910159 | 0.16771934 | 0.21431245 | 0 | *** |
| LF_A_/ MF1_A_ *vs* MF1_A_ / MF1_B_ | 0.18228362 | 0.15898298 | 0.20558426 | 0 | *** |
| LF_A_/ MF2_A_ *vs* - MF1_A_ / MF1_B_ | 0.16933348 | 0.14603284 | 0.19263412 | 0 | *** |
| MF1_A_/ MF2_A_ *vs* MF1_A_ / MF1_B_ | 0.17324882 | 0.14994818 | 0.19654946 | 0 | *** |
| LF_B_/ MF2_B_ *vs* MF1_A_ / MF1_B_ | 0.15803628 | 0.13473565 | 0.18133692 | 0 | *** |
| LF_B_/ MF1_B_ *vs* MF1_A_ / MF1_B_ | 0.16504021 | 0.14173957 | 0.18834084 | 0 | *** |
| MF1_B_/ MF2_B_ *vs* MF1_A_ / MF1_B_ | 0.16866766 | 0.14536702 | 0.1919683 | 0 | *** |

**Supplementary Table 18** **| Ancient gene block associations described in the published studies as summarized by Lysak et al. (2017)^30^. The gene block associations in the A and B genome assembly of *B. juncea* and C genome using the assembly of Belsar et al (2018)^15^**

| **Species** | **Family** | | **A-B** | **F-G** | **K-L** | **M-N** | **O-P** | **Q-R** | **T-U** | **W-X** | **Wb-R** | **V-K-L-Wa-Q-X** |
| --- | --- | --- | --- | --- | --- | --- | --- | --- | --- | --- | --- | --- |
| **ACK ancestor** | | | | | | | | | | | | |
| *Arabidopsis lyrata* | Camelineae A (I) | | + | + | + | + | + | + | + | + | – | – |
| *Arabidopsis thaliana* | Camelineae A (I) | | + | – | – | + | + | + | + | + | – | – |
| *Ballantinia antipoda* | Microlepidieae A (I) | | + | + | + | + | – | + | – | + | – | – |
| *Boechera divaricarpa* | Boechereae A (I) | | + | + | + | + | + | + | + | + | – | – |
| *Boechera stricta* | Boechereae A (I) | | + | + | + | + | + | + | + | + | – | – |
| *Camelina sativa* | Camelineae A (I) | | + | + | + | + | + | + | + | + | – | – |
| *Cardamine amara* | Cardamineae A (I) | | + | + | + | + | + | + | + | + | – | – |
| *Cardamine flexuosa* | Cardamineae A (I) | | + | + | + | + | + | + | + | + | – | – |
| *Cardamine hirsuta* | Cardamineae A (I) | | + | + | + | + | + | + | + | + | – | – |
| *Capsella rubella* | Camelineae A (I) | | + | + | + | + | + | + | + | + | – | – |
| *Crucihimalaya wallichii* | Crucihimalayeae A (I) | | + | + | + | + | + | + | + | + | – | – |
| *Hornungia alpina* | Descurainieae A (I) | | + | + | + | + | + | + | + | + | – | – |
| *Neslia paniculata* | Camelineae A (I) | | + | + | + | + | + | + | + | + | – | – |
| *Pachycladon exilis* | Microlepidieae A (I) | | + | + | + | + | + | + | + | + | – | – |
| *Stenopetalum lineare* | Microlepidieae A (I) | | – | + | + | + | – | – | + | + | – | – |
| *Stenopetalum nutans* | Microlepidieae A (I) | | – | + | + | + | – | – | – | + | – | – |
| *Transberingia bursifolia* | Crucihimalayeae A (I) | | + | + | + | + | + | + | + | + | – | – |
| *Turritis glabra* | Turritideae A (I) | | + | + | + | + | + | + | + | + | – | – |
| *Transberingia bursifolia* | Crucihimalayeae A (I) | | + | + | + | + | + | + | + | + | – | – |
| *Turritis glabra* | Turritideae A (I) | | + | + | + | + | + | + | + | + | – | – |
| **PCK ancestor** | | | | | | | | | | | | |
| *Brassica oleracea* | Brassiceae B (II) | | – | – | + | – | + | – | – | – | + | + |
| *Brassica rapa* | Brassiceae B (II) | | – | – | + | – | + | – | – | – | + | + |
| *Brassica napus* | Brassiceae B (II) | | – | – | + | – | + | – | – | – | + | + |
| *Calepina irregularis* | Calepineae B (EII) | | + | + | + | + | + | – | + | – | + | + |
| *Conringia orientalis* | Conringieae B (EII) | | + | + | + | + | + | – | + | – | + | + |
| *Caulanthus amplexicaulis* | Thelypodieae ? (EII) | | + | – | ? | + | – | – | + | – | + | + |
| *Glastaria glastifolia* | Isatideae B (II) | | + | + | + | + | + | – | + | – | + | + |
| *Goldbachia laevigata* | Calepineae B (EII) | | + | + | + | + | – | – | + | – | + | + |
| *Myagrum perfoliatum* | Isatideae B (II) | | + | + | + | + | + | – | + | – | + | + |
| *Noccaea caerulescens* | Coluteocarpeae B (EII) | | + | + | + | + | – | – | – | – | + | + |
| *Noccaea jankae* | Coluteocarpeae B (EII) | | + | + | + | + | + | – | – | – | + | + |
| *Ochthodium aegyptiacum* | Sisymbrieae B (II) | | + | + | + | + | + | – | + | – | + | + |
| *Raparia bulbosa* | Coluteocarpeae B (EII) | | + | + | + | + | + | – | – | – | + | + |
| *Schrenkiella parvula* | Unassigned B (EII) | | + | + | + | + | + | – | + | – | + | + |
| *Thellungiella salsuginea* | Eutremeae B (EII) | | + | + | + | + | + | – | + | – | + | + |
| **Unresolved ancestral genome** | | | | | | | | | | | | |
| *Arabis alpina* | Arabideae ? (EII) | | + | + | + | + | – | + | – | + | – | – |
| *Biscutella laevigatad* | Biscutelleae C (?) | | + | + | + | + | + | – | ? | – | + | – |
| **This study** | | | | | | | | | | | | |
| *Brassica juncea* A genome | Brassiceae B | LF | + | + | + | + | + | + | + | + | – | – |
|  |  | MF1 | – | – | + | + | + | + | + | + | – | – |
|  |  | MF2 | + | + | + | + | + | + | + | + | – | – |
| *Brassica juncea* B genome | Brassiceae B | LF | + | + | + | + | + | + | + | + | – | – |
|  |  | MF1 | + | – | + | + | + | + | + | + | – | – |
|  |  | MF2 | + | + | + | – | + | + | + | + | – | – |
| *Brassica oleracea^#^* | Brassiceae B | LF | + | + | + | + | + | + | + | + | – | – |
|  |  | MF1 | – | – | + | + | + | + | + | + | – | – |
|  |  | MF2 | – | + | + | + | + | + | + | + | – | – |

**Supplementary Table 20 |** **Genomic block fragmentation patterns in the A and B sub-genomes of *B. juncea* Varuna**

|  | A genome | | | | B genome | | | |
| --- | --- | --- | --- | --- | --- | --- | --- | --- |
| Block | LF | MF1 | MF2 | Total blocks | LF | MF1 | MF2 | Total blocks |
| A | A06, A10 | A09 | A08, A08^*^ | 4 | B04, B06,B07^*^ | B08, B07 | B07 | 6 |
| B | A09, A06 | A07, A05^*^ | A08, A08^*^ | 6 | B06 | B07 | B01, B07, B07^*^ | 5 |
| C | A10, A09^*^ | A08 | A05, A01* | 5 | B06, B06* | B01, B01* | B04 | 5 |
| D | A09, A03 | A03 | A02 | 4 | B06, | B01 | B08 | 3 |
| E | A07 | A02 | A07 | 3 | B07, B04 | B02 | B04, B07 | 5 |
| F | A05, A07 | A01 | A03 | 4 | B05 | B01 | B01, B07, B08* | 5 |
| G | A07 | - | A03 | 2 | B05 | - | B08 | 2 |
| H | A07, A09* | A04 | A03* | 4 | B08, B04 | B05 | B03 | 4 |
| I | A07, A09* | A04 | A03 | 4 | B08, B04 | B05 | B03 | 4 |
| J | A05 | A04 | A08 | 3 | B04, B04 | B05 | B03 | 4 |
| K | A06 | A02 | A09 | 3 | B08 | B06 | B06 | 3 |
| L | A06 | A02 | A09, A09 | 4 | B08 | B06 | B01, B06 | 4 |
| M | A06, A09 | A06, A03* | A01 | 5 | B03 | B04, B04*, B08* | B02 | 5 |
| N | A09 | A04, A03 | A07, A01 | 5 | B03 | B04, B02*, B08* | B02 | 5 |
| O | A09. A09 | A03 | A02 | 4 | B03 | B03 | B02 | 3 |
| P | A09 | A03 | A02 | 3 | B03 | B03 | B02 | 3 |
| Q | A06, A10* | A09, A02* | A02, A03* | 6 | B08, B08* | B01, B02* | B06, B03* | 6 |
| R | A10 | A02 | A03 | 3 | B08 | B02, B02 | B03 | 4 |
| S | A04, A08* | A07 | A04, A05 | 5 | B01, B05 | B04 | B05 | 4 |
| T | A01 | A04, A03*, A05* | A08, A08* | 6 | B02 | B07, B04*, B08* | B01, B01, B07* | 7 |
| U | A01 | A03, A06* | A08 | 4 | B02, B02 | B08, B08 | B07, B07, B01 | 7 |
| V | A06, A06* | A02, A02* | A09 | 5 | B08, B08* | B06 | B06, B01* | 5 |
| W | A10, A06* | A02, A02* | A03, A03*, A09* | 7 | B08 | B02, B02, B02, B06* | B03, B03, B01* | 8 |
| X | A06, A03*, A07*, A10* | A09, A02* | A02 | 7 | B04, B08*, B08* | B01, B02* | B06 | 6 |

*Small blocks

**Supplementary Table 21 |** Gene block associations between the ancestral paleogenomes that constituted the A, B and C genomes. Red, green and blue colours represent the LF, MF1 and MF2 paleogenomes, respectively

(a) Intra-paleogenome contiguous gene block associations

|  | A genome | B genome | C genome |
| --- | --- | --- | --- |
| A– B | C–A– B– M–M(A06) | H–A– B*–C (B06) | A– B–C–*–B (C05) |
|  | A– B–T–U–T–B (A08) | F–A– B–A (B07) | T–U–A– B–M (C08) |
|  |  | A– B–U–T–B–U–F (B07) |  |
| F– G | B–F–G*–H (A07) | F– G–*–S (B05) | B–F– G–H (C07) |
|  | O–F– G–H (A03) | A–F– G–H (B08) | O–F– G–H (C03) |
| I-J | J – I–S–S (A04)  W–J – I–W (A03) | M–J – I–X (B04)  J – I–S–S (B05)  W–J – I–W (B03) | J – I – V(C04)  J – I–S–S (C04)  W–J – I–W (C03) |
| K– L | V–K– L–V–W–Q– H– X(A06) | D–V–K– L–V–U–T (B08) | X–V–K– L–V–W–Q–X (C07) |
|  | P–V–K– L–V–W–Q–X (A02) | L–V–K– L–V–W–Q (B06) | P*–V–K– L–V–W–Q–X (C02) |
|  | C–V–K– L–O–P (A09) | C–V–K– L–V–K (B06) | C–V–K– L–O–P (C09) |
| M– N | B–M– N–I–H (A09) | M–M– N (B03) | B–M– N–I–H (C08) |
|  | X–M– N–T–U (A03) | X–M– N–T–U (B08) | X–M– N–T–U(C07) |
|  | D–*–M– N–T–U(A01) |  | C–*–M– N–T–U (C01) |
| O– P | L–O– P–*–B (A09) | O–O–P–*–M (B03) | L–O– P–*–W (C09) |
|  | F–O– P–W (A03) | O–O–P–W (B03) | F–O– P– W (C03) |
|  | E–O– P*–V (A02) | M–O– P–W (B02) | E–O–P*–V (C02) |
| Q– R | W–X–Q–R (A10) | W–X–Q–R (B08) | W–X–Q–R (C09) |
|  | W–X–Q–R (A02) | W–X–Q–R (B02) | W–X–Q–R (C02) |
|  | W–X–Q–R (A03) | W–X–Q–R (B03) | W–X–Q– R (C03) |
| T– U | M–N–T–U (A01) | R–N–T–U–W (B02) | M–N–T–U (C01) |
|  | M–N–T–U–D (A03) | M–N–T–U–V (B08) | M–N–T–U (C07) |
|  | B–T–U–T–B (A08) | T–B–T–U–F–L (B01) | S–T–U–A (C08) |

(b) Intra-paleogenome non-contiguous gene block associations

|  | A genome | B genome | C genome |
| --- | --- | --- | --- |
| Q–X | V–W–Q–H–X–M (A06) | U–Q–X–M–N–T (B08) | L–V–W–Q–X (C07) |
|  | L–W–Q–X–H (A09) | V–W–Q–X (A02) | L–W–Q–X–H (C09) |
|  | V–W–Q–X (A02) | V–W–Q–X (B06) | V–W–Q–X (C02) |
|  | A–C–*–W–X–Q–R (A10) | I–*–W–X–Q–R (B08) | P–*–W–X–Q–R (C09) |
| W– X–Q–R | E–W–X–Q–R (A02)  W–I–J–W–X–Q–R (A03) | U–W–X–Q–R–M (B02)  I–J–W–X–Q–R (B03) | E–W–X–Q–R (C02)  W–I–J–W–X–Q– R (C03) |
| V–K–L-(V) | U–V–K– L–V–W–Q–H (A06)  P*–V–K– L–V–W–Q–X (A02) | D–V–K– L–V–U–T (B08)  L–V–K– L–V–W–Q (B06) | X–V–K– L–V–W–Q–X (C07)  P*–V–K– L–V–W–Q–X (C02) |
|  | C–V–K– L–O–P (A09) | C–V–K– L–V–K (B06) | C–V–K– L–O–P (C09) |
| W-E | X–W–E–O (A02) | P–W-E (B02) | X–W–E–O (C02) |

(c) Inter-paleogenome gene block associations

|  | A genome | B genome | C genome |
| --- | --- | --- | --- |
| J-I-S-S | J–I–S–S–T* (A04) | J–I–S–S–* (B05) | J–I–S–S–T (C04) |
| R-W-J-I-W-P-O | R–W–J–I–W–P–O (A03) | R–Q–W–J–I–W–P–O (B03) | R–W–J–I–W–P–O (C03) |

(d) Inter-paleogenome same gene block associations

|  | A genome | B genome | C genome |
| --- | --- | --- | --- |
| M-M | M–M (A06) | – | M–M (C03) |
| O-O | – | O–O (B03) | – |
| S-S | S–S (A04) | S–S (B05) | S–S (C04) |
| E-E | E–E (A07) | E–E (B07) | E–E (C06) |
