## Supplementary Figures for "A chromosome-scale assembly of allotetraploid *Brassica juncea* (AABB) elucidates comparative architecture of the A and B genomes"

**Bju-A01**

**Bju-A02**

**Bju-A03**

**Bju-A04**

**Bju-A05**

**Bju-A06**

**Bju-A07**

**Bju-A08**

**Bju-A09**

**Bju-A10**

**Bju-B01**

**Bju-B02**

**Bju-B03**

**Bju-B04**

**Bju-B05**

**Bju-B06**

**Bju-B07**

**Bju-B08**

0

20

40

60

80

100

120

140

160

180

200

220

240

260

280

300

320

**Supplementary Figure 1 | Genetic map of *Brassica juncea* Varuna x Heera (VH) F_1_DH population**. The LGs have been named as Bju-A01 to Bju-A10 and Bju-B01 to Bju-B08^17^. The black bands in the LG bar represent the mapped markers. A total of 3,780 markers were mapped on the 18 LGs; these include 833 anchor markers and 2,947 GBS based SNPs derived from *Sph*I-*Mluc*I restriction enzyme combination. The map covered a total genetic length of 3,232.3 cM. Sidebar, cM.

**Bju-A01**

**Bju-A02**

**Bju-A03**

**Bju-A04**

**Bju-A05**

**Bju-A06**

**Bju-A07**

**Bju-A08**

**Bju-A09**

**Bju-A10**

**Bju-B01**

**Bju-B02**

**Bju-B03**

**Bju-B04**

**Bju-B05**

**Bju-B06**

**Bju-B07**

**Bju-B08**

0

20

40

60

80

100

120

140

160

180

200

220

240

260

280

300

**Supplementary Figure 2 | Genetic map of *Brassica juncea* Tumida x Varuna (TuV) F_1_DH population**. The LGs belonging to the A genome were named as Bju-A01 to Bju-A10 and those to the B genome as Bju-B01 to Bju-B08^17^. The black bands in the LGs represent the mapped markers. A total of 9,041 markers – 524 anchor markers common with the VH population and 8,517 GBS based SNPs generated using the *Hinf*I-*Hpy*CH4IV enzyme combination were mapped on 18 LGs. The map covered a total genetic length of 3,417.4 cM. Sidebar, cM.

**BniB01**

**BniB02**

**BniB03**

**BniB04**

**BniB05**

**BniB06**

**BniB07**

**BniB08**

0

20

40

60

80

100

120

140

160

**Supplementary Figure 3 | Genetic map of *Brassica nigra* Sangam x 2782 F_1_DH population.** The LGs have been named as Bni-B01 to Bni-B08. The black bands in the LGs represent the mapped markers. A total of 1,765 markers were mapped on to the 8 LGs and covered a total genetic length of 822.6 cM. The map incorporates 197 anchor markers common with the VH population and 1,568 GBS-based SNPs generated using the *Sph*I-*Mluc*I enzyme combination. Sidebar, cM.

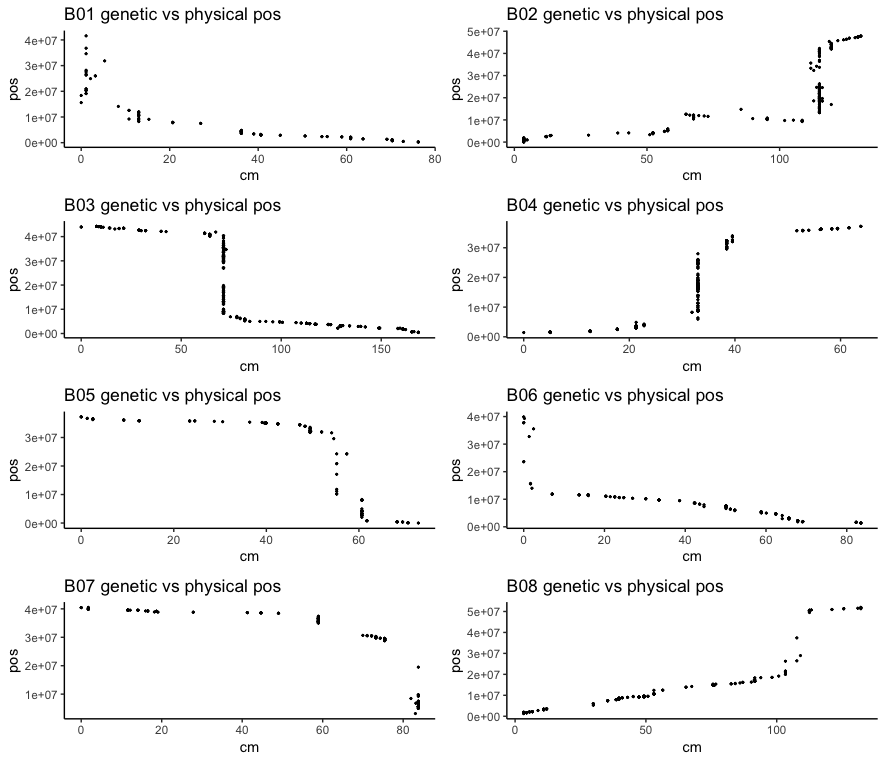

**Supplementary Figure** **4 | Relationship between the GBS markers on the genetic map of *B. nigra* Sangam x 2782 F_1_DH population and physical position of the respective marker tags on the *B. nigra* genomic sequences**. Genetic positions of the markers have been shown at the x-axis and physical positions at the y-axis. A linear relationship was found between the physical and genetic distances of the markers present on the LGs except at the centromeric regions where the rate of recombination would be much lower than in the other chromosomal regions.

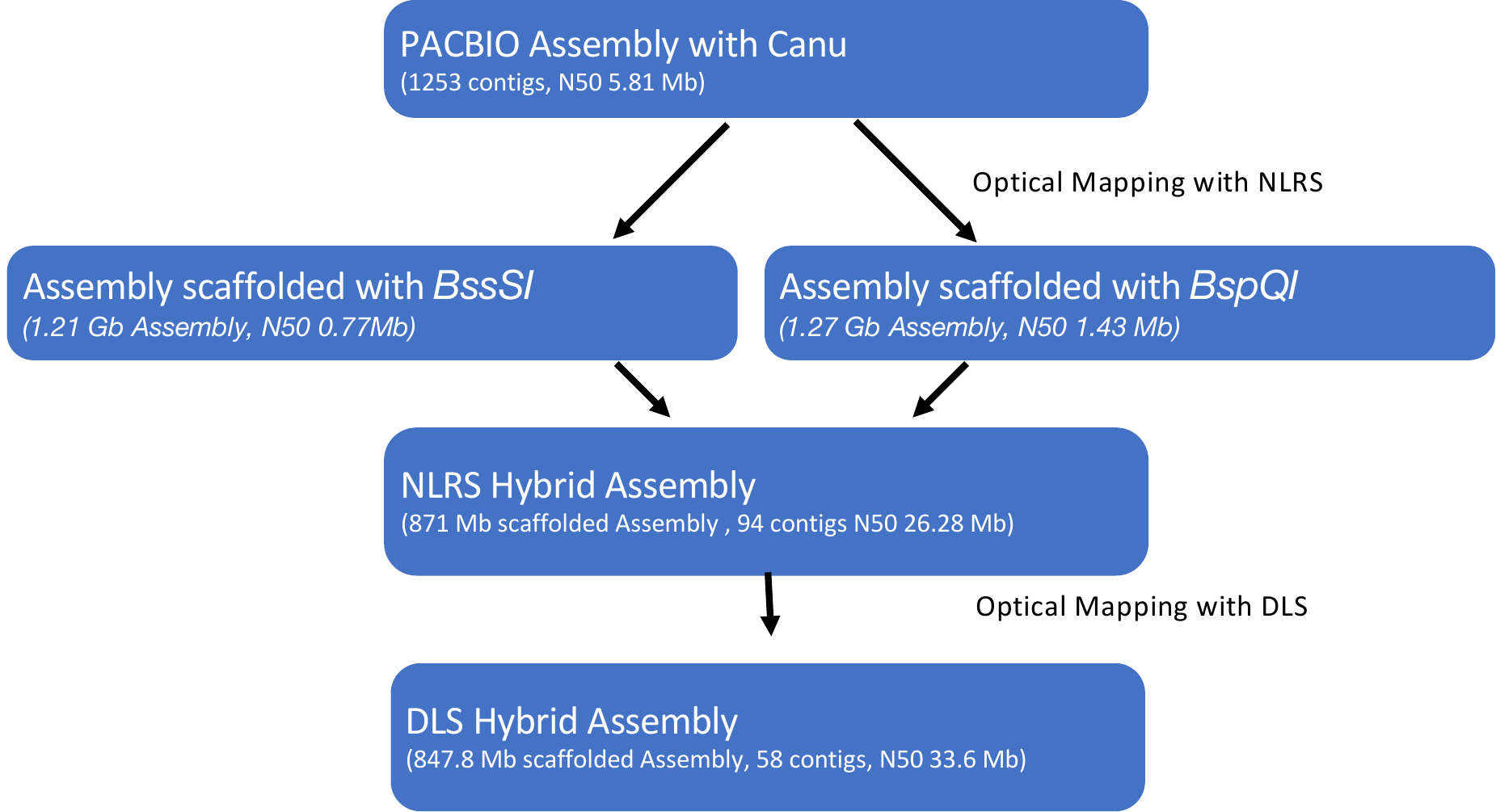

**Supplementary figure 5 |** **Workflow in BioNano optical mapping based hierarchical scaffolding analysis of *Brassica juncea* genome.** Two NLRS based labeling reactions were carried out, c-maps were generated, and the assembled PacBio contigs were aligned to these. A hybrid assembly of the two c-maps and corrected PacBio contigs yielded 94 scaffolds, covering ~871 Mb of the genome. The assembled scaffolds were aligned again with the DLS maps to generate 847.8 Mb assembly consisting of 58 scaffolds.

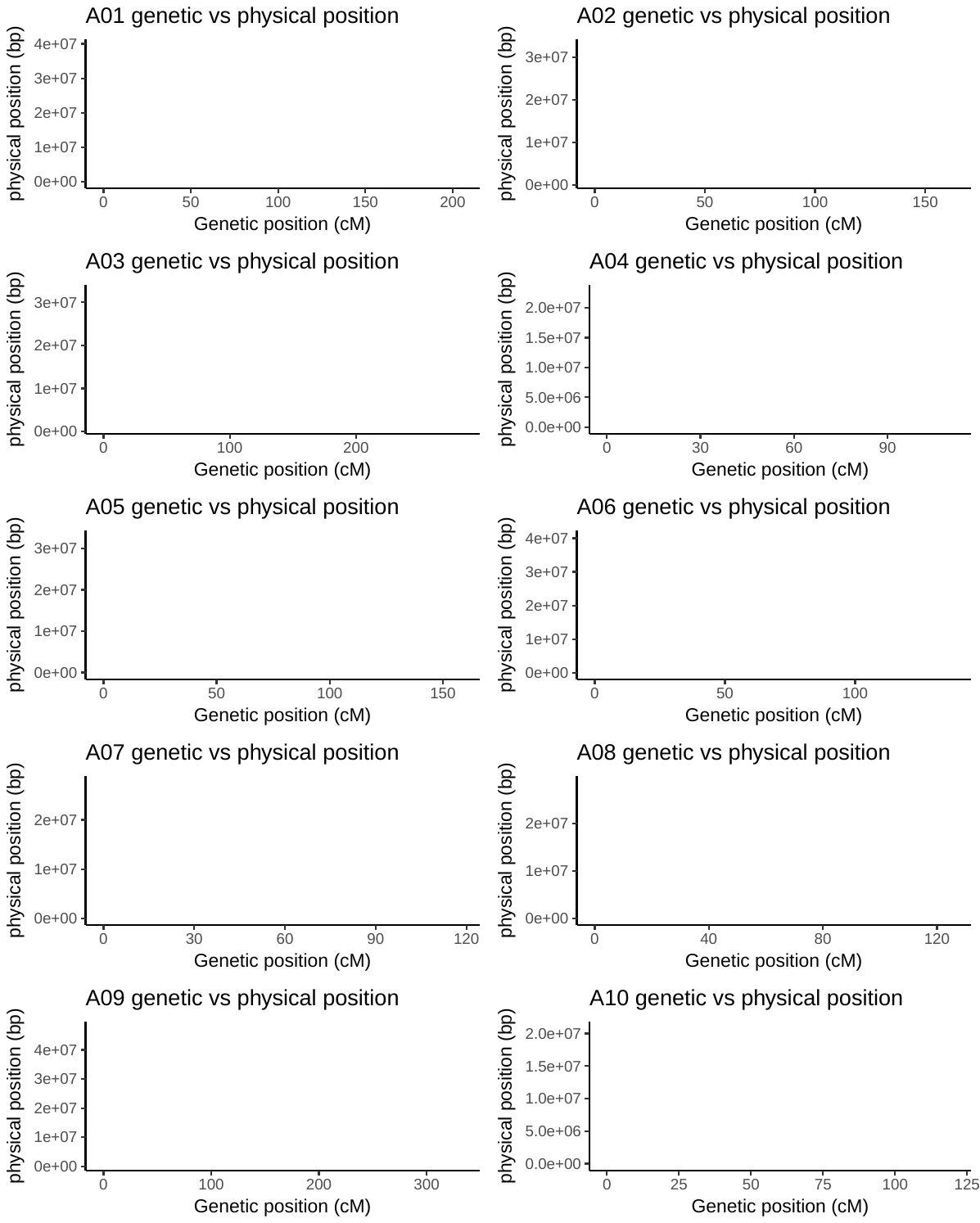

**Supplementary Figure** **6 | Relationship between the GBS markers on the *Brassica juncea* VH (Varuna x Heera) F1DH population linkage map and physical position of the respective marker tags on the Varuna assembled genome.** Genetic positions of the markers have been shown at the x-axis and physical positions are shown at the y-axis. Linear relationship was found between the physical and genetic distances of the markers present on an LG except at the centromeric region where the rate of recombination would be much lower than the other chromosomal regions.

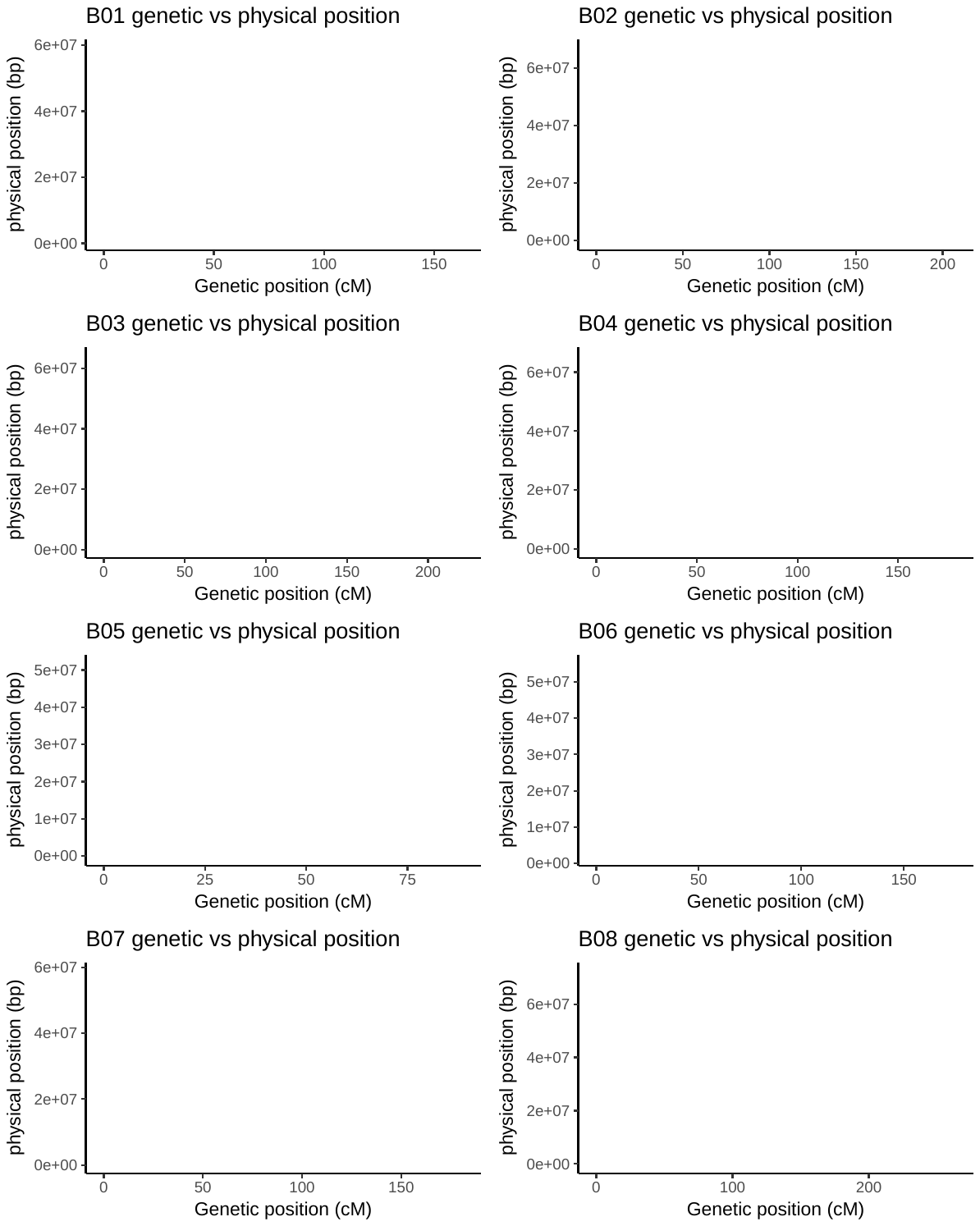

**Supplementary Figure 6** | contd..

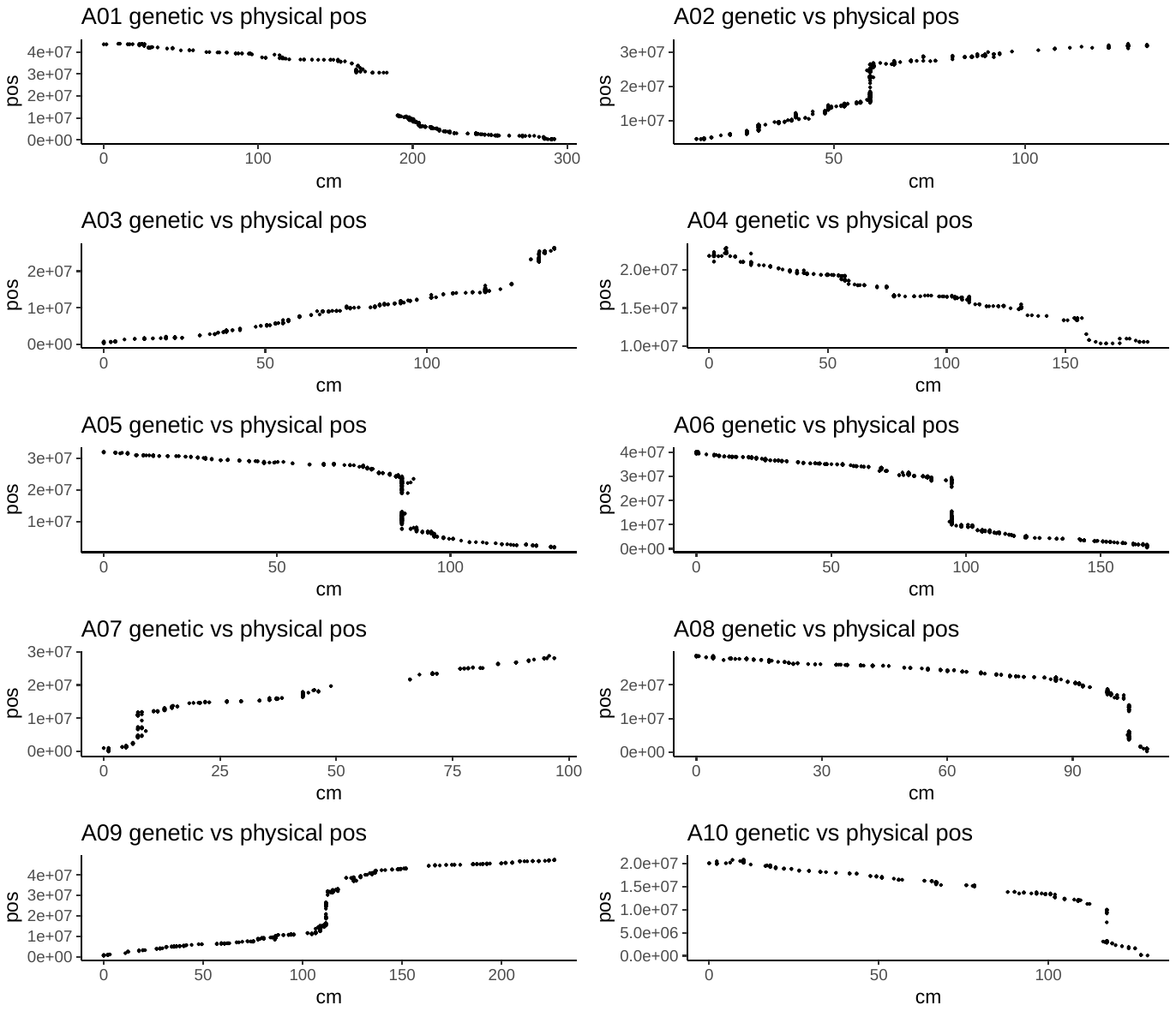

**Supplementary Figure** **7 | Relationship between the GBS markers on the *Brassica juncea* Tumida x Varuna F_1_DH population genetic map and the physical position of the respective GBS markers on the assembled Varuna genome.** Genetic positions of the markers in centimorgan (cM) have been shown at the x-axis and physical positions are shown at the y-axis. Linear relationship was found between the physical and genetic positions of the makers except at the centromeric regions. A region of the B07 pseudochromosome could not be covered with the linkage map analysis as no polymorphic marker could be detected between Tumida and Varuna parental lines in the first 30Mb region of the LG B07.

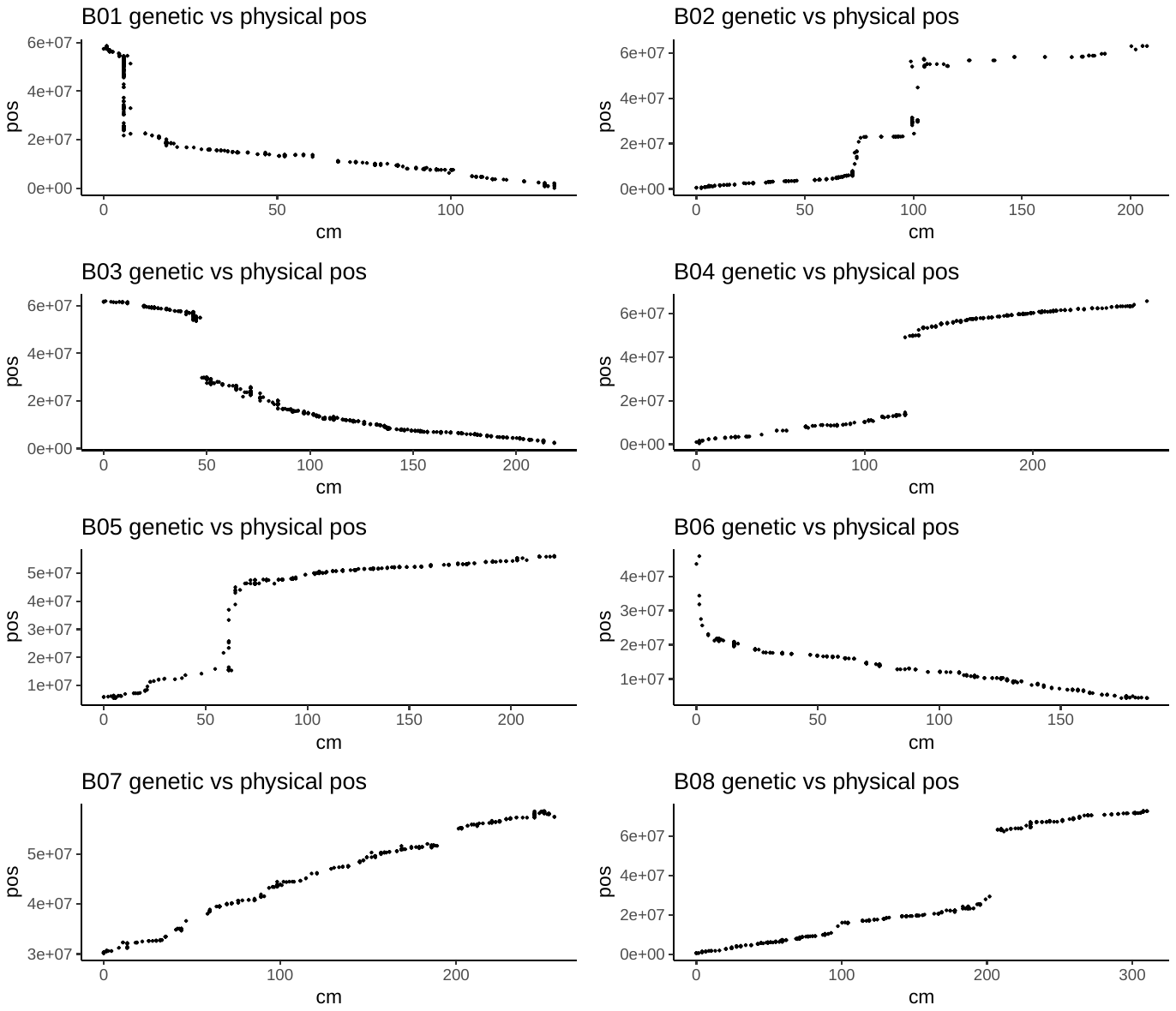

**Supplementary Figure** **7|** contd..

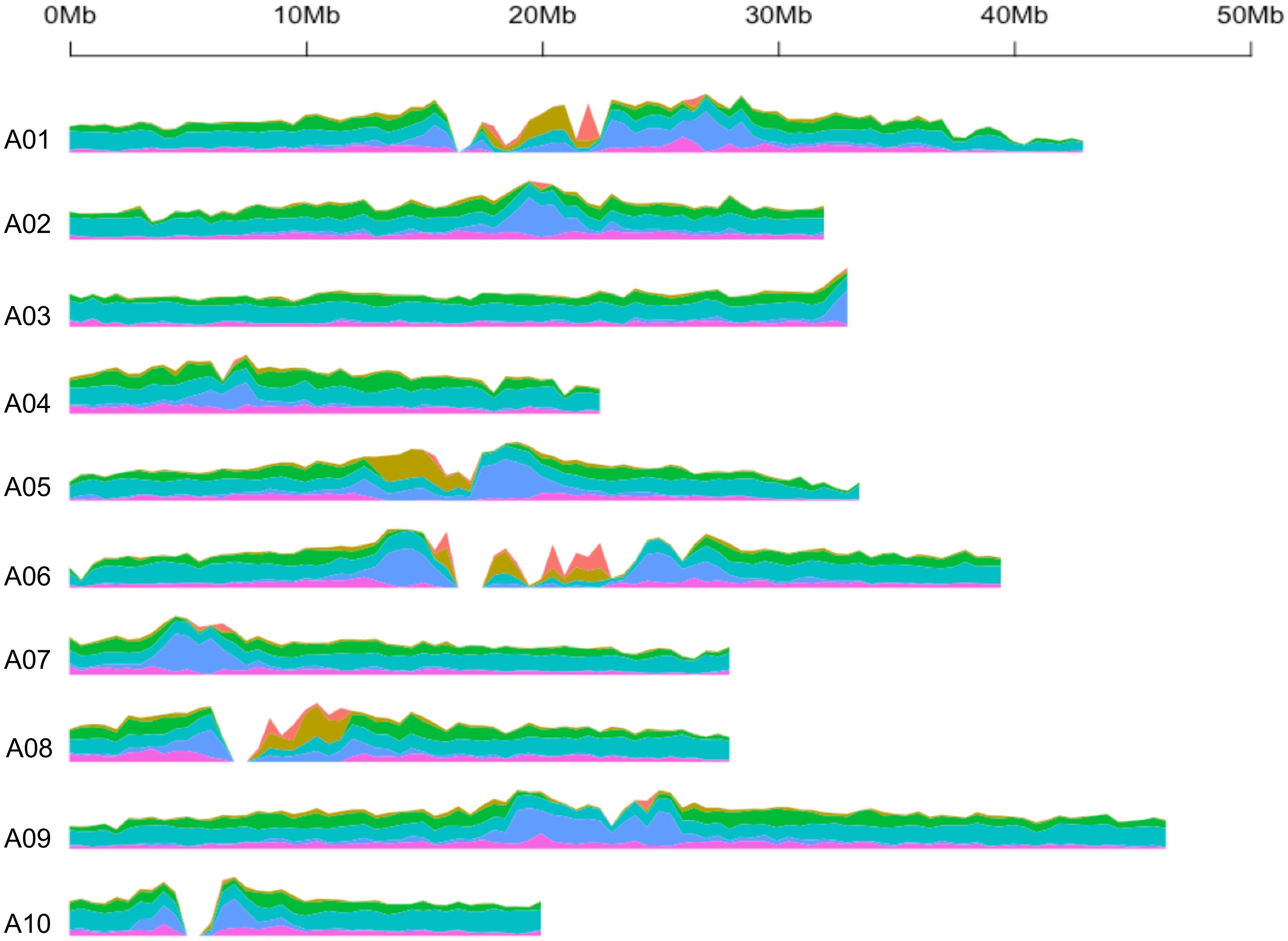

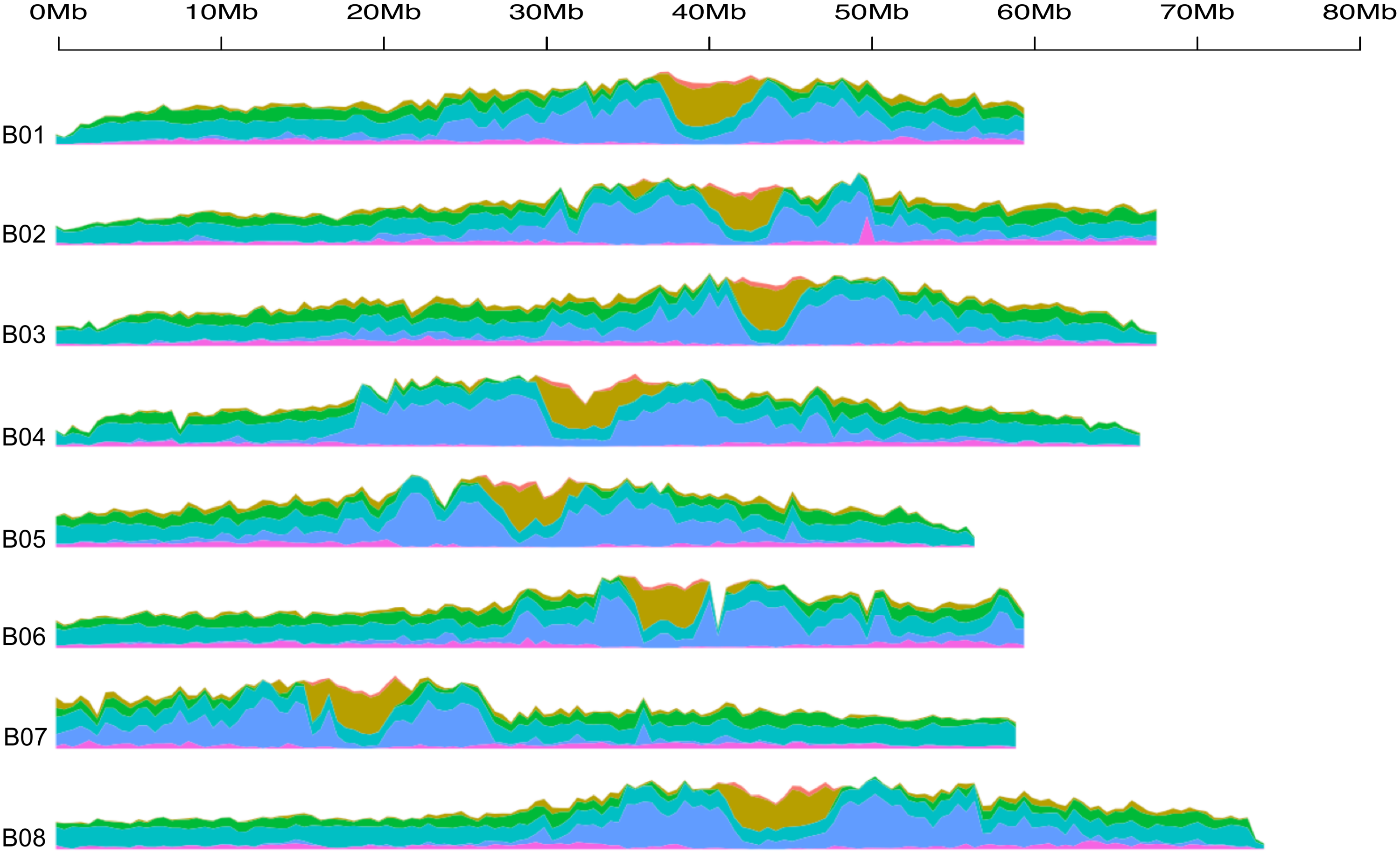

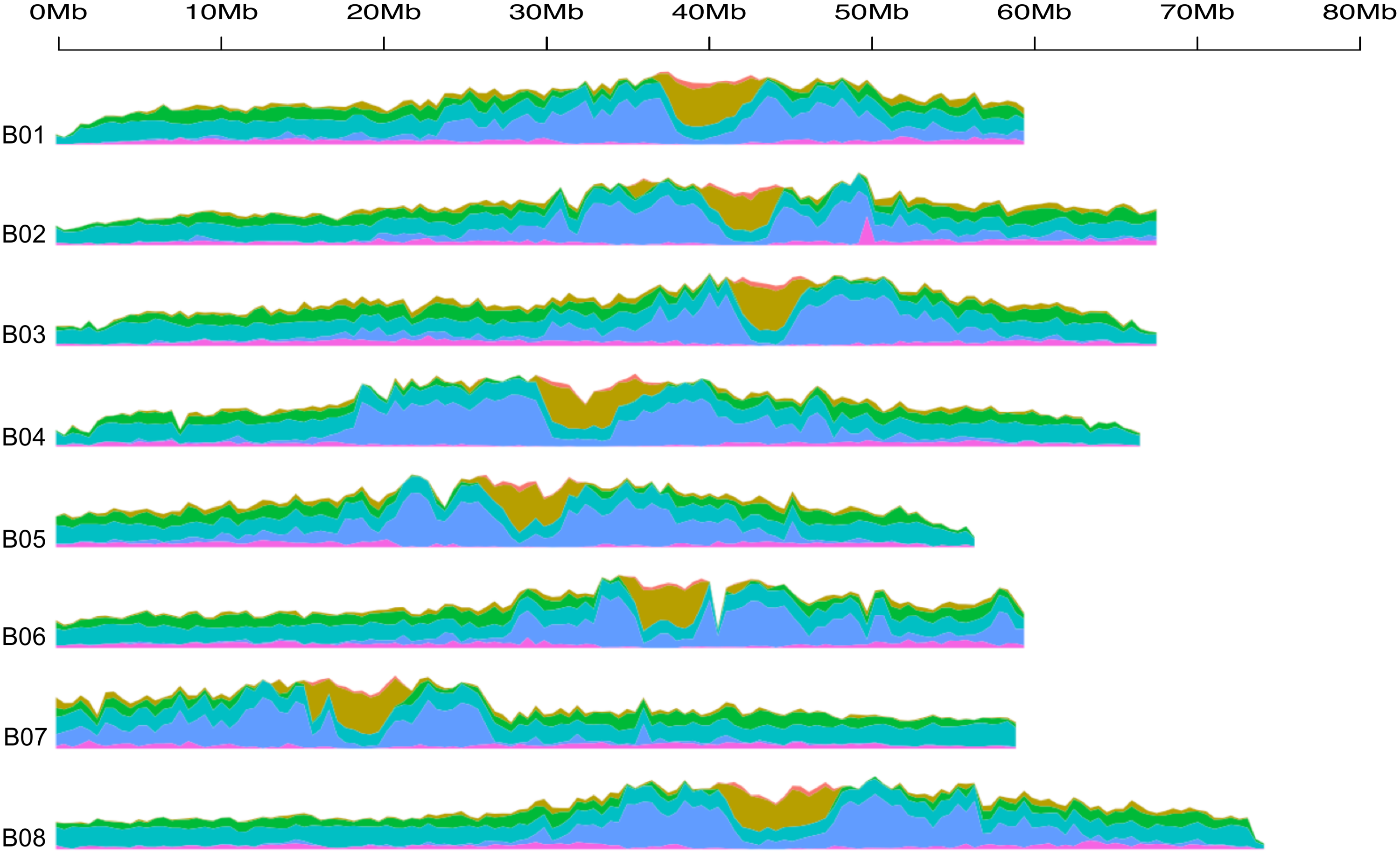

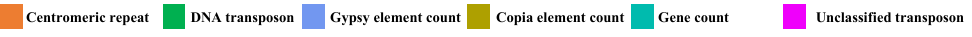

**Supplementary Figure** **8 | Distribution of different types of transposable elements on *Brassica juncea* pseudochromosomes.** Centromeric regions are boxed and show much higher content of TEs in the B genome.

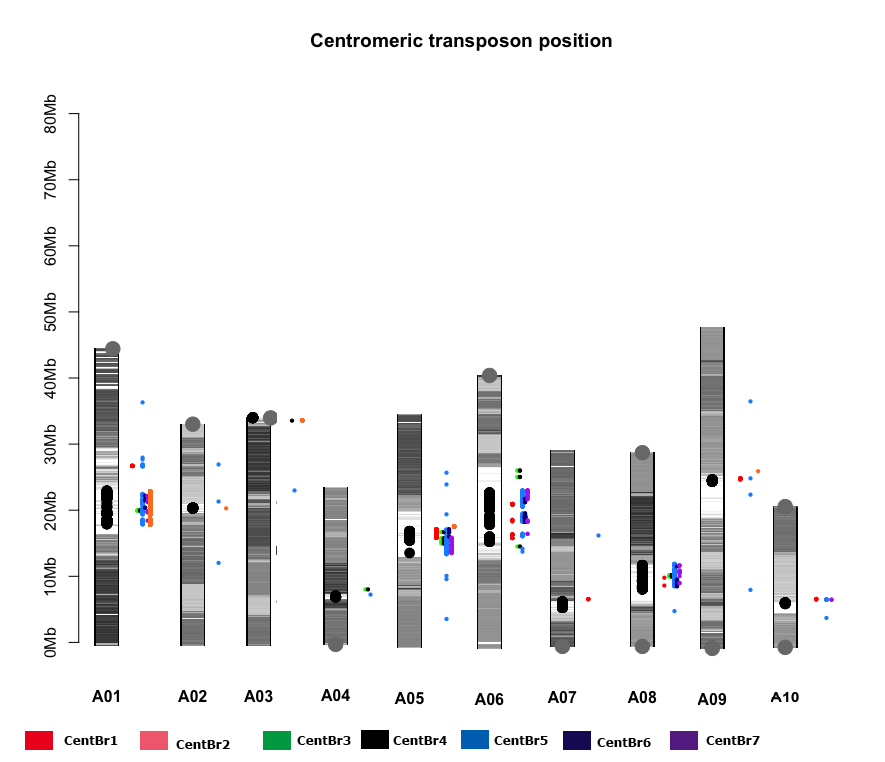

**Supplementary Figure** **9** | **Position of the predicted A genome centromeric sequences on different pseudochromosomes of the A genome of *Brassica juncea*.** The number of dots shows the extent of repetition on the pseudochromosomes. Different centromere-specific sequences have been assigned different colors.

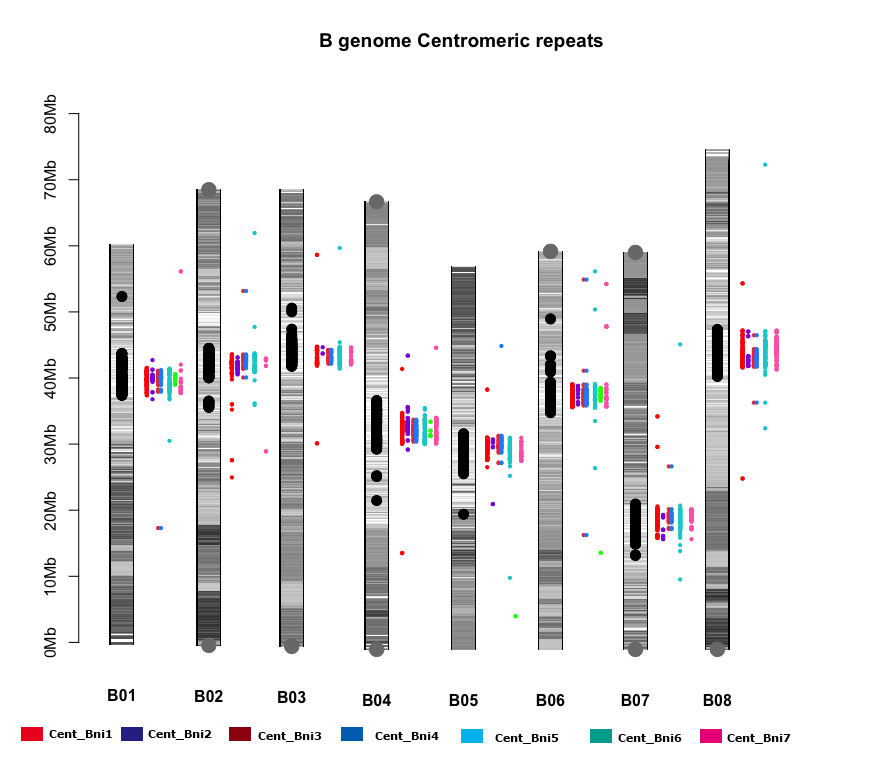

**Supplementary Figure 10** | **Position of the predicted B genome centromeric sequences on different pseudochromosomes of the B genome of *Brassica juncea*.** Different centromere-specific sequences have been assigned different colours. Number of dots show the extent of repetition on the pseudochromosomes.

**Comparison of A01- Chiifu and Tumida assemblies vs Varuna assembly**

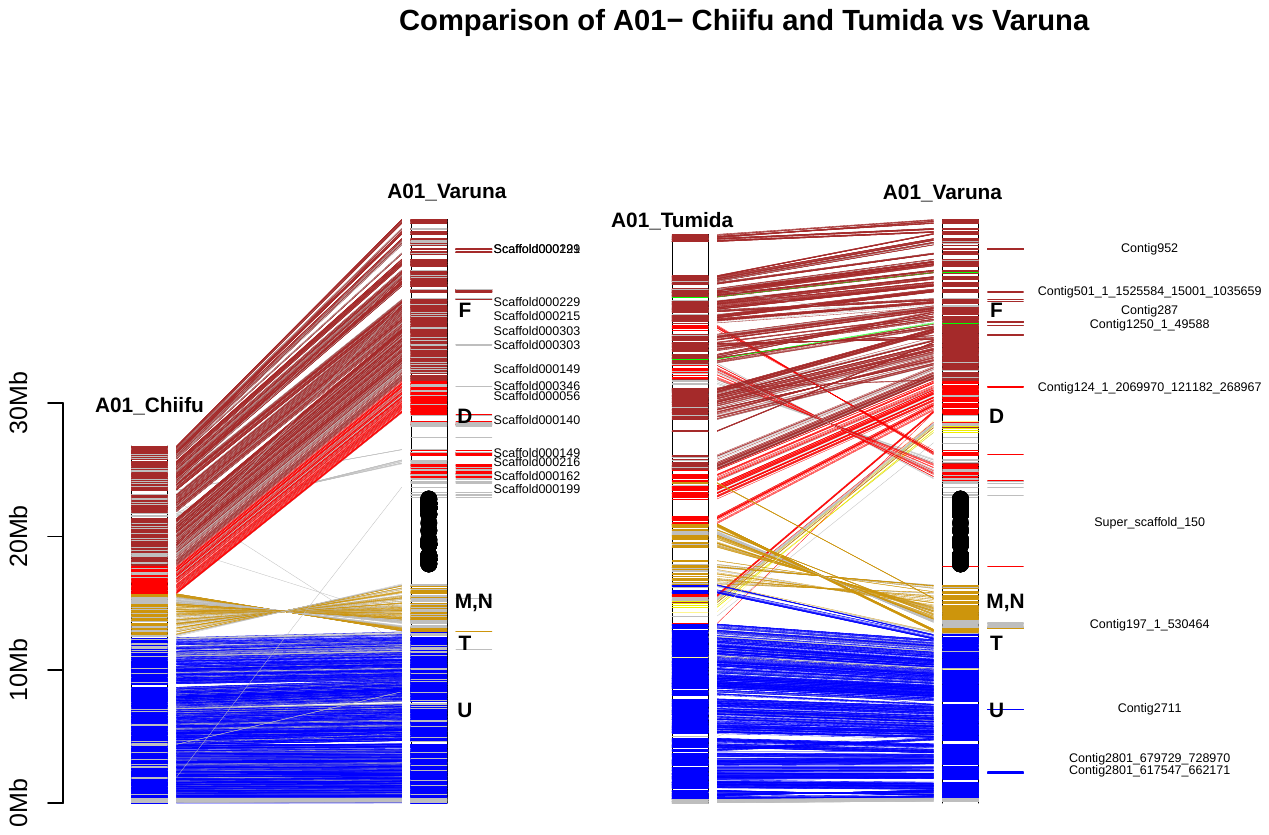

**Comparison of A02- Chiifu and Tumida assemblies vs Varuna assembly**

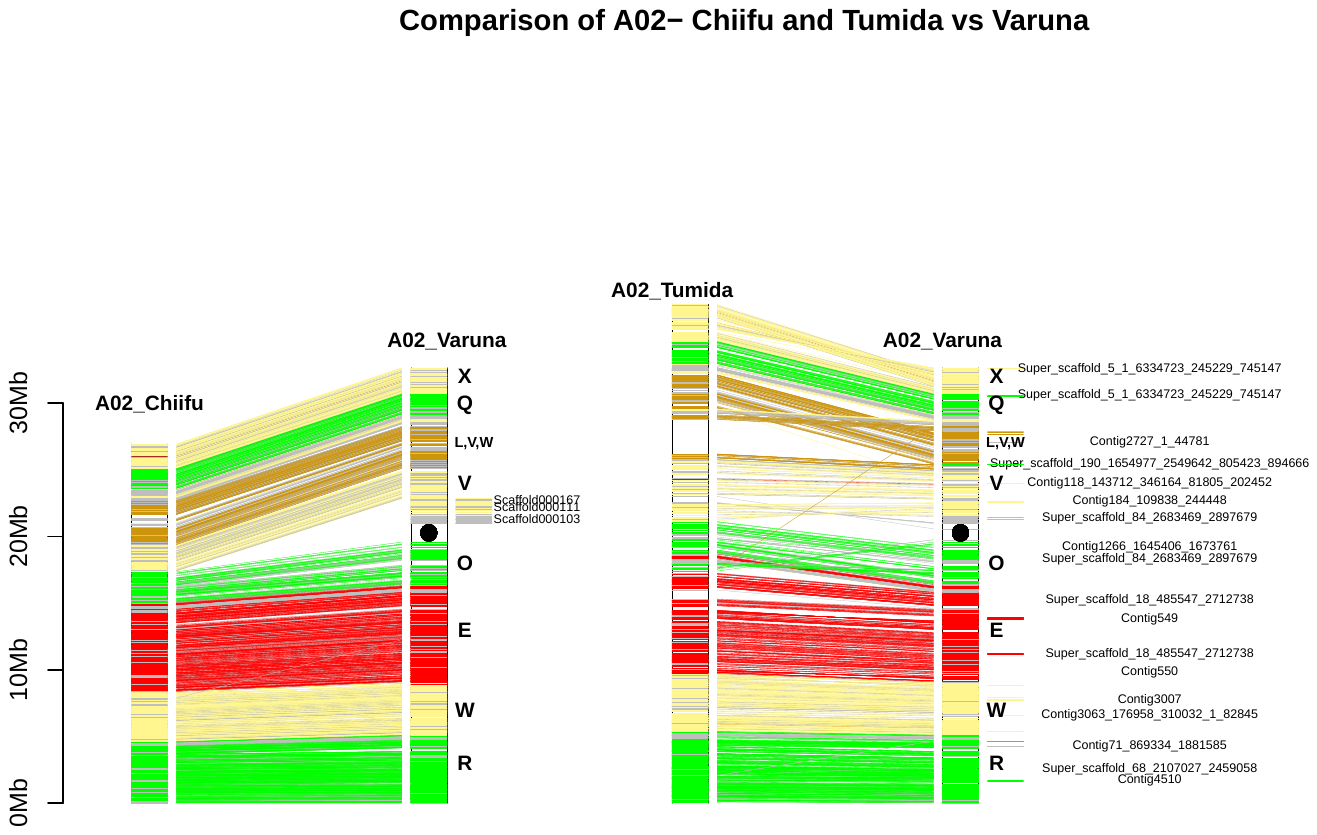

**Supplementary Figure 11** | **Comparison of the genome assembly of *B. juncea* Varuna with previous assemblies of *B. rapa* Chiifu for the A genome and *B. juncea* variety Tumida for both the A and B genomes.** Comparisons were made based on gene order on each pseudochromosome in the three assemblies. Scaffolds and contigs of the earlier assemblies that were unassigned to any pseudochromosomes could be assigned in the Varuna assembly (shown on the right of the Varuna pseudochromosomes; only some have been demonstrated due to inadequate space). The A genome pseudochromosomes of Varuna matched the Chiifu assembly more than the Tumida assembly. The B genome assembly of Tumida requires revision.

**Comparison of A03- Chiifu and Tumida assemblies vs Varuna assembly**

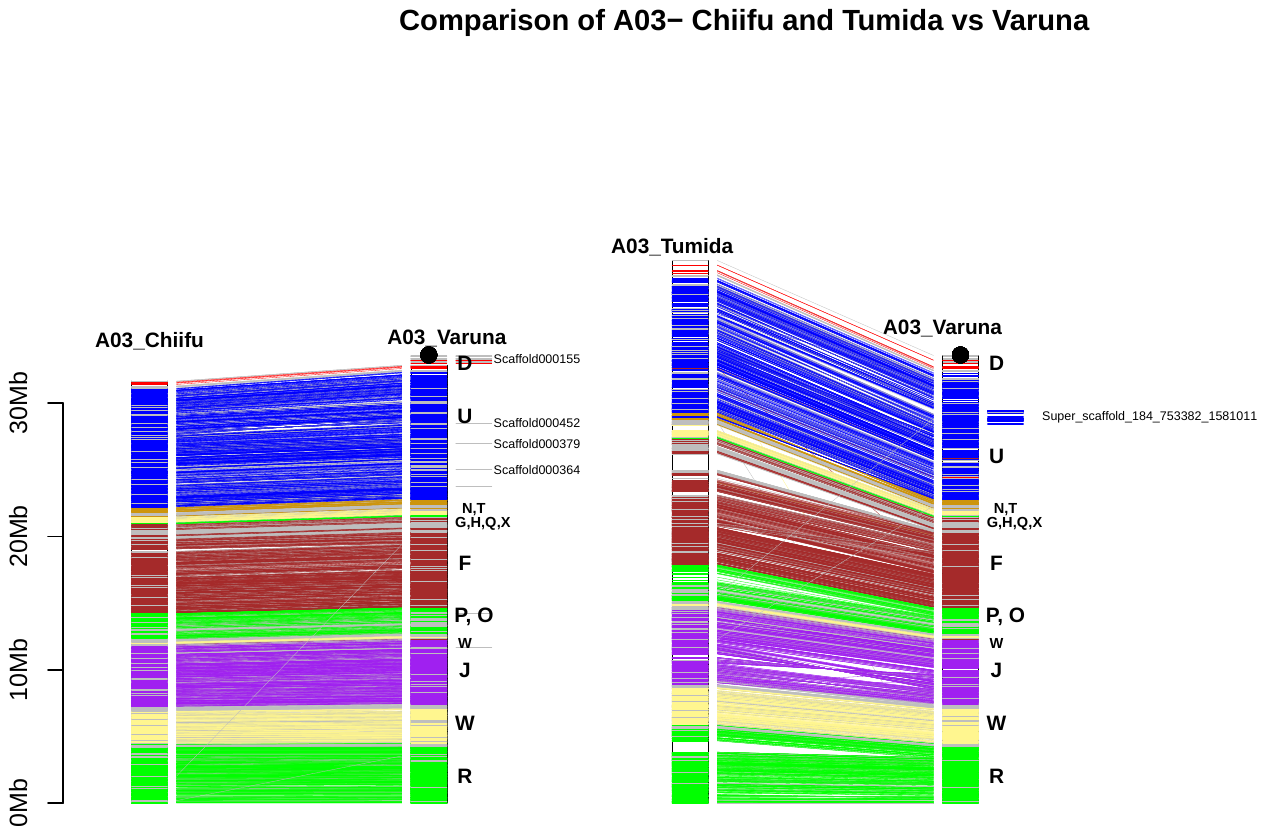

**Comparison of A04- Chiifu and Tumida assemblies vs Varuna assembly**

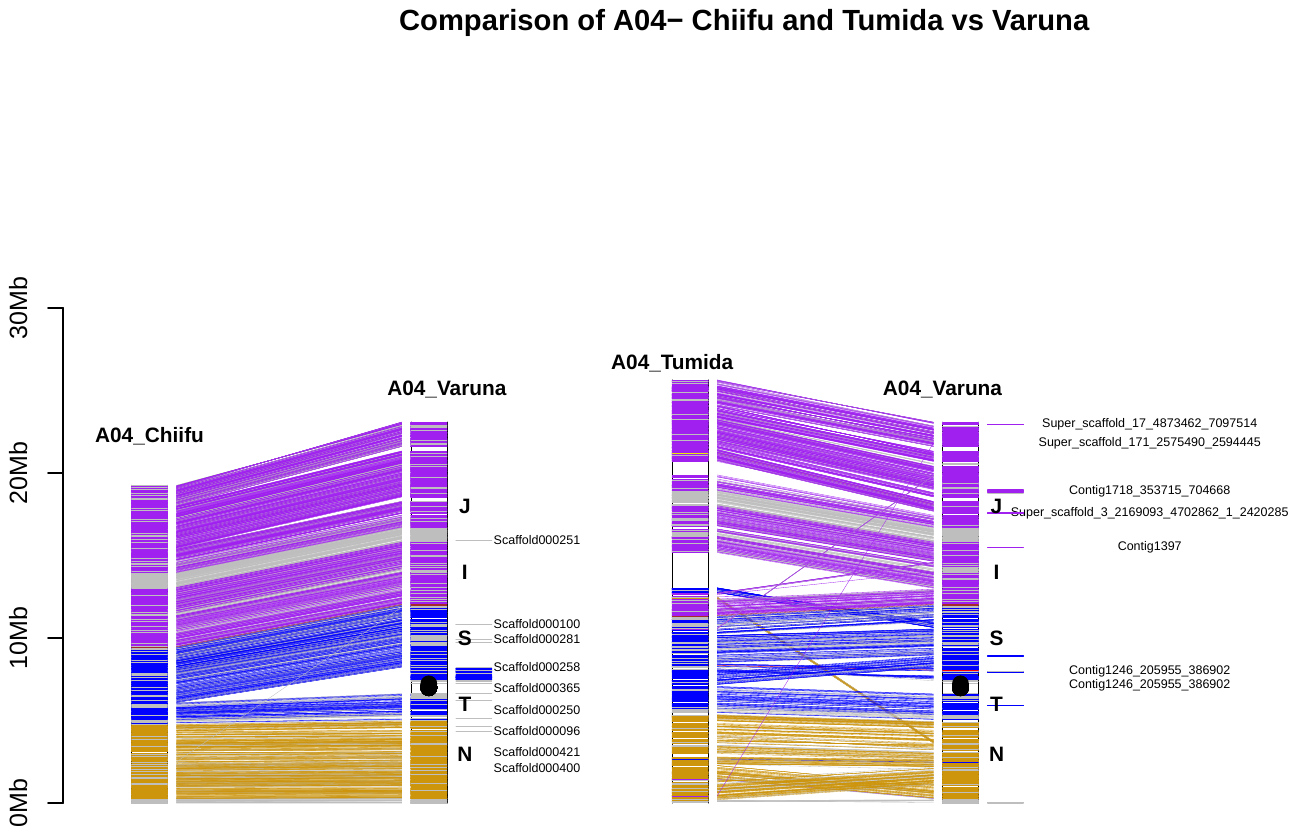

**Supplementary Figure 11 |** contd..

**Comparison of A05- Chiifu and Tumida assemblies vs Varuna assembly**

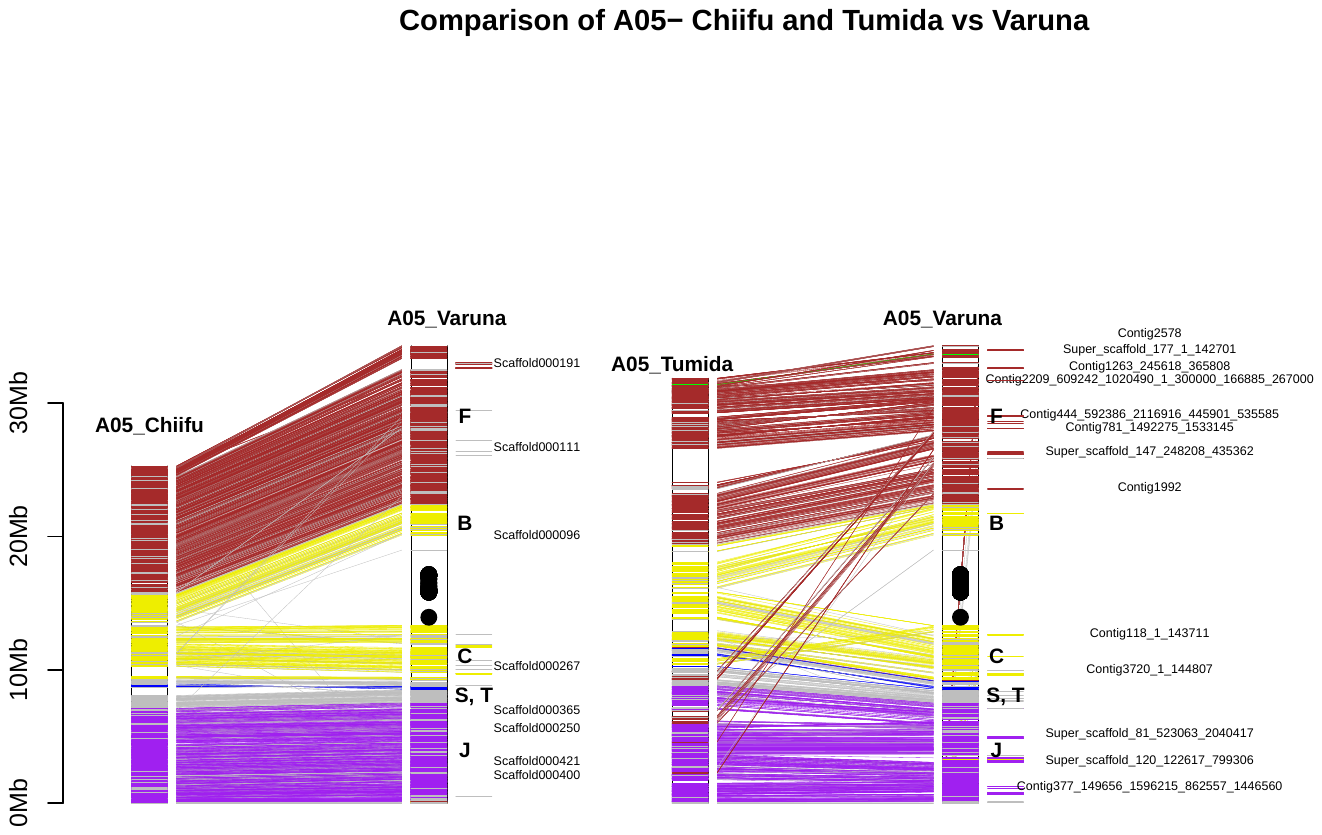

**Comparison of A06- Chiifu and Tumida assemblies vs Varuna assembly**

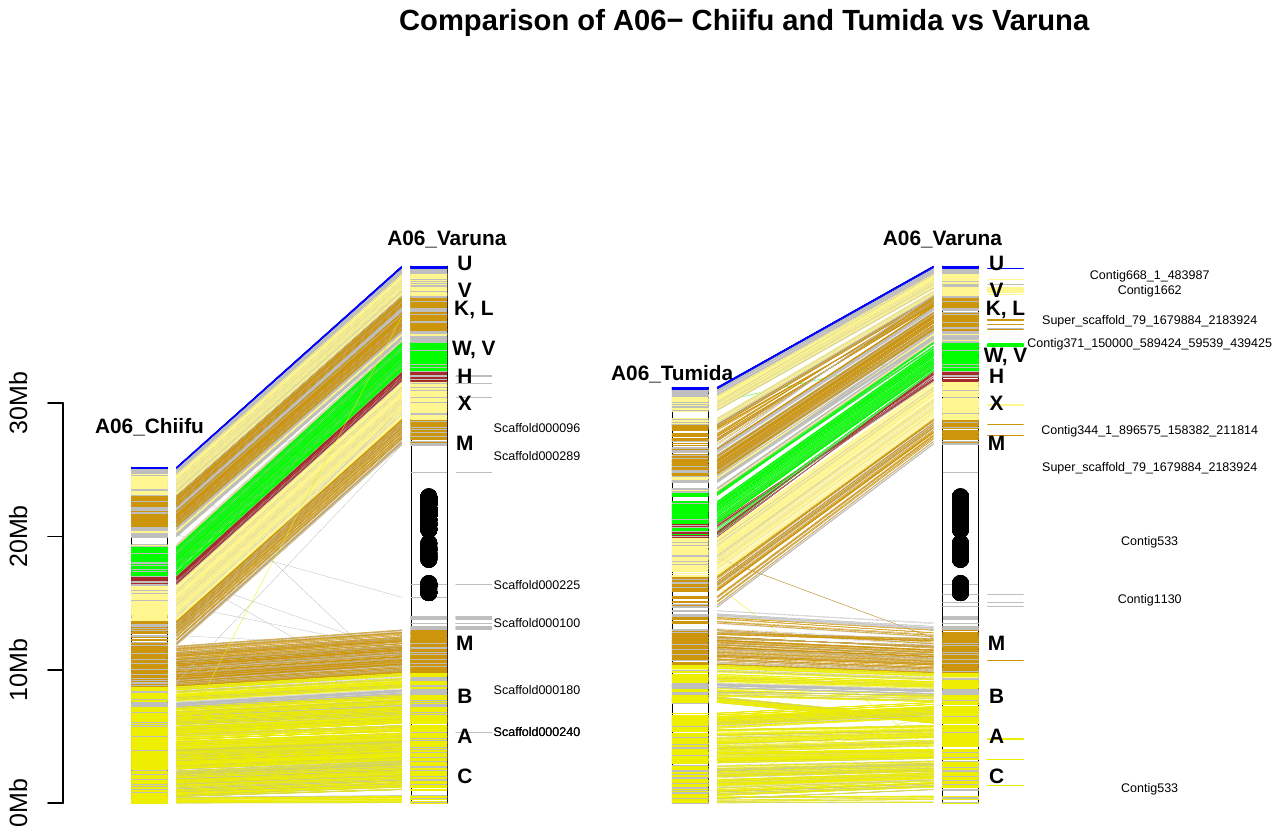

**Supplementary Figure 11 |** contd..

**Comparison of A07- Chiifu and Tumida assemblies vs Varuna assembly**

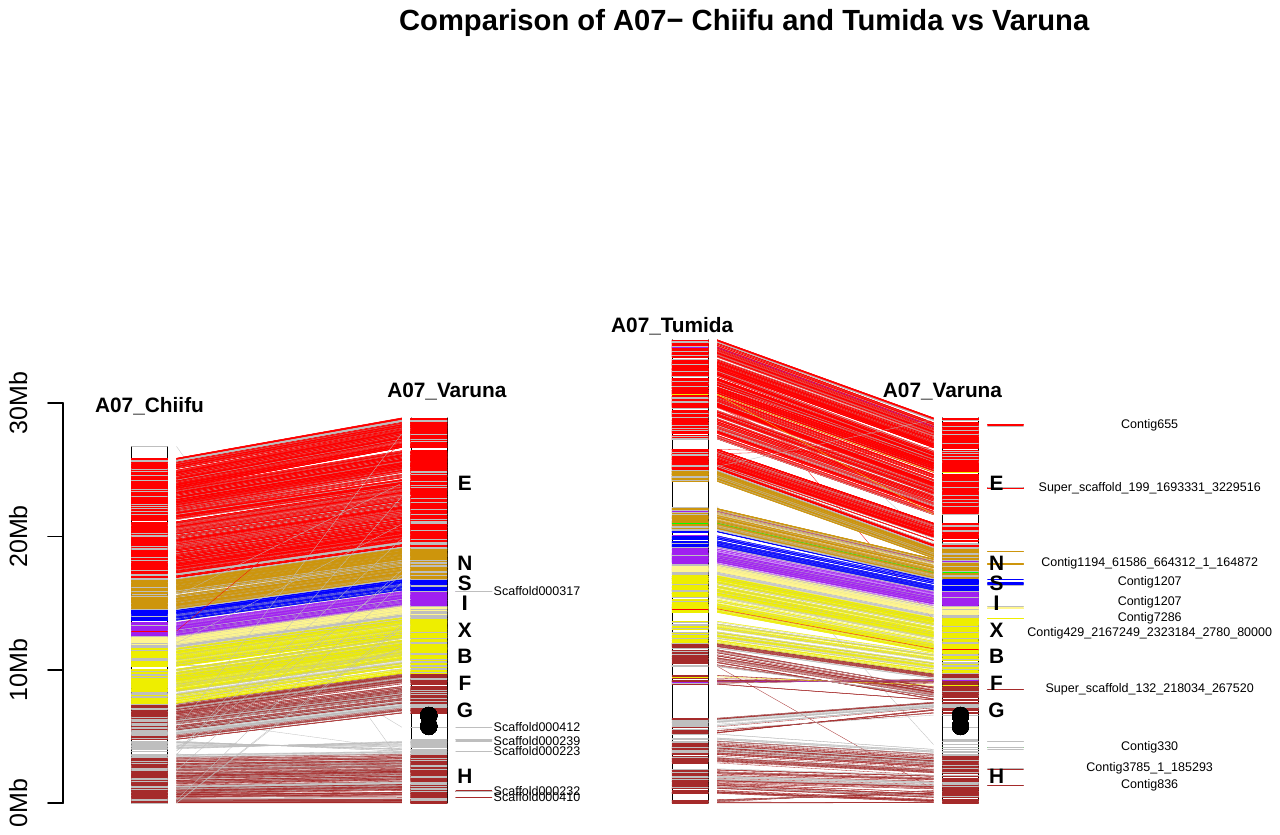

**Comparison of A08- Chiifu and Tumida assemblies vs Varuna assembly**

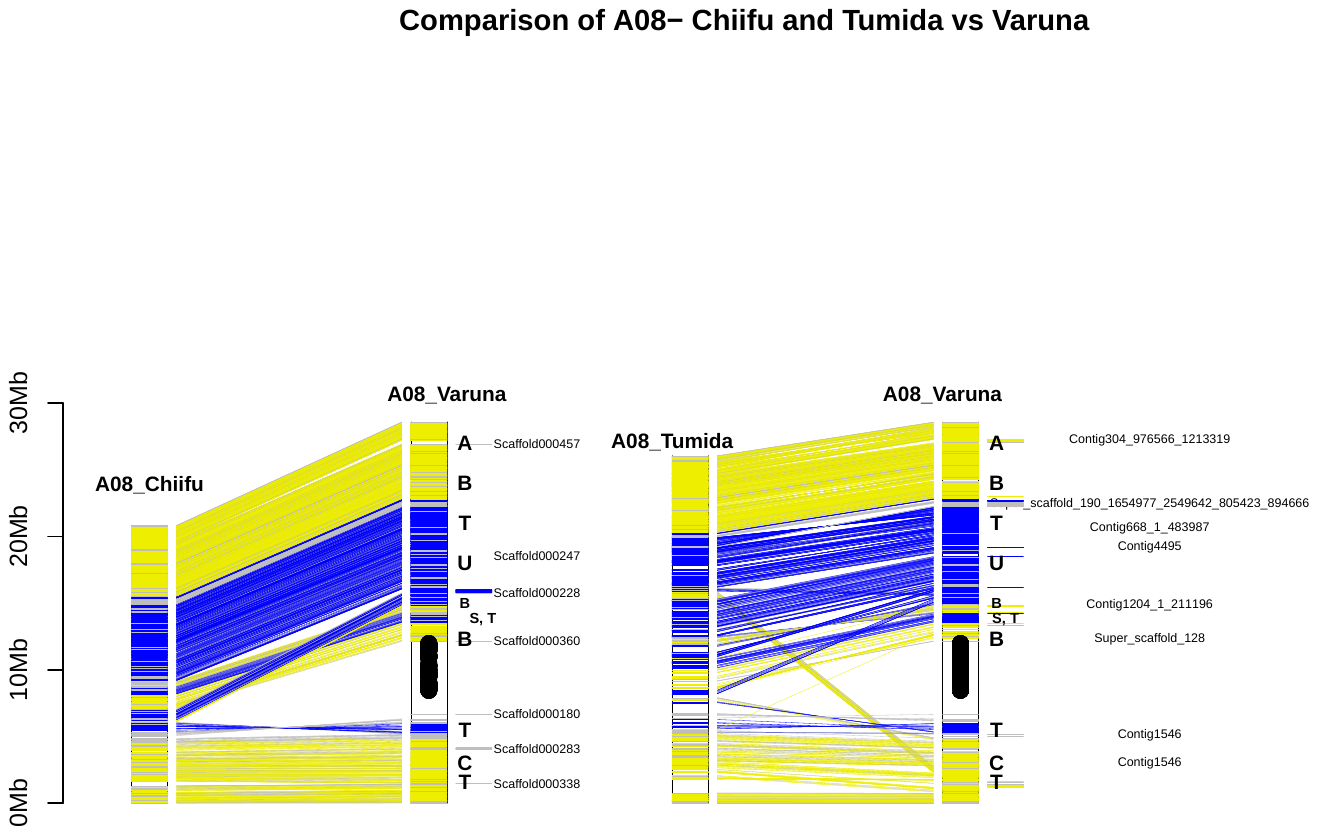

**Supplementary Figure 11 |** contd..

**Comparison of A09- Chiifu and Tumida assemblies vs Varuna assembly**

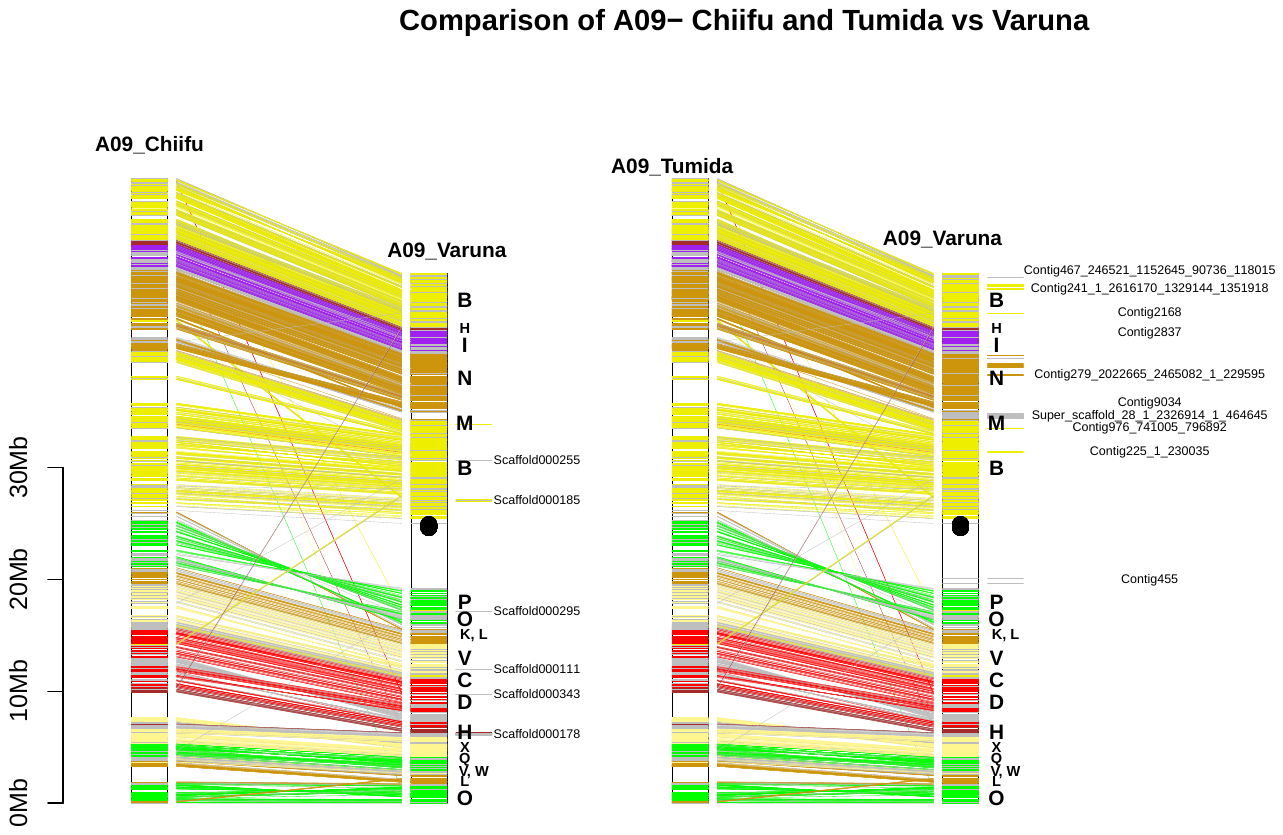

**Comparison of A10- Chiifu and Tumida assemblies vs Varuna assembly**

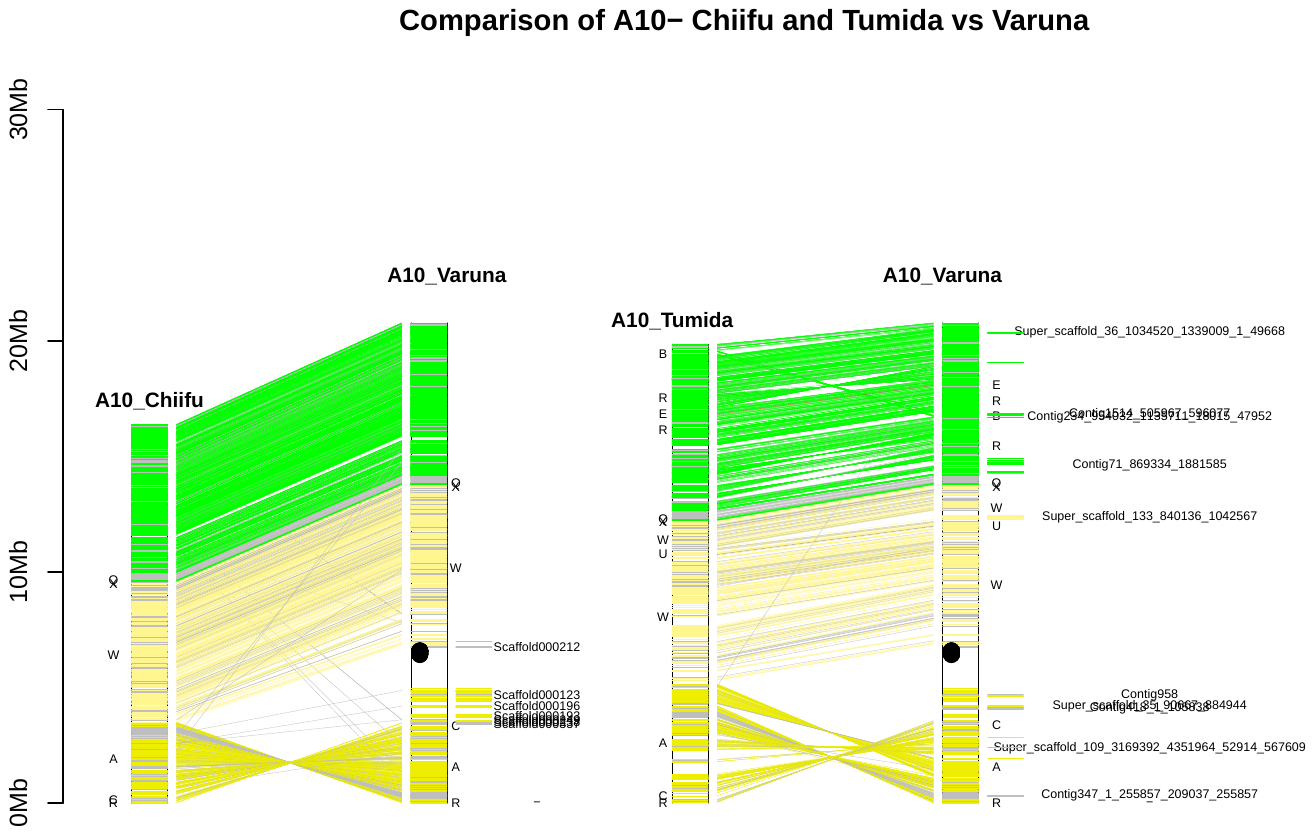

**Supplementary Figure 11 |** contd..

**Comparison of B01 - B04 pseudochromosomes- Tumida vs Varuna assembly**

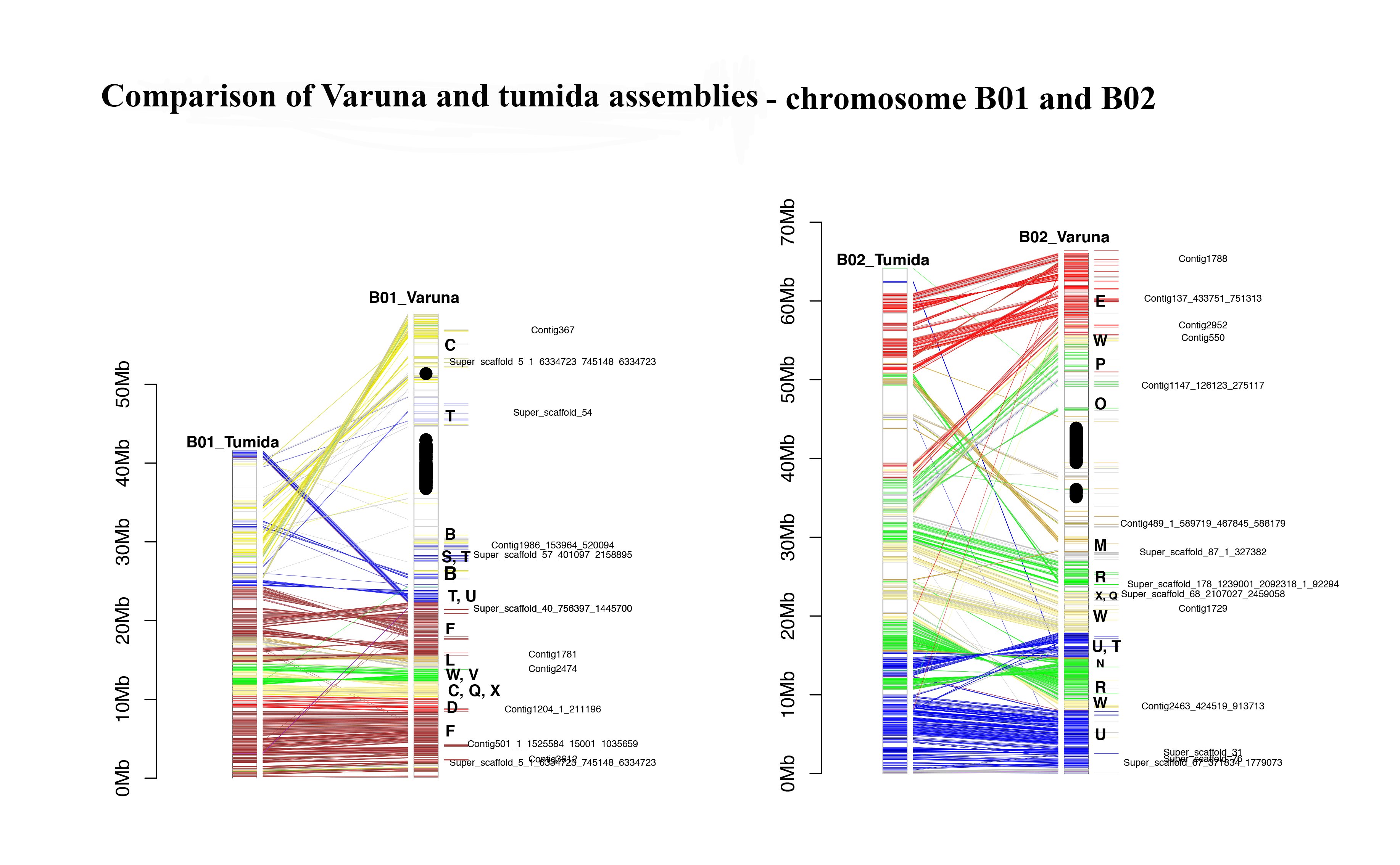

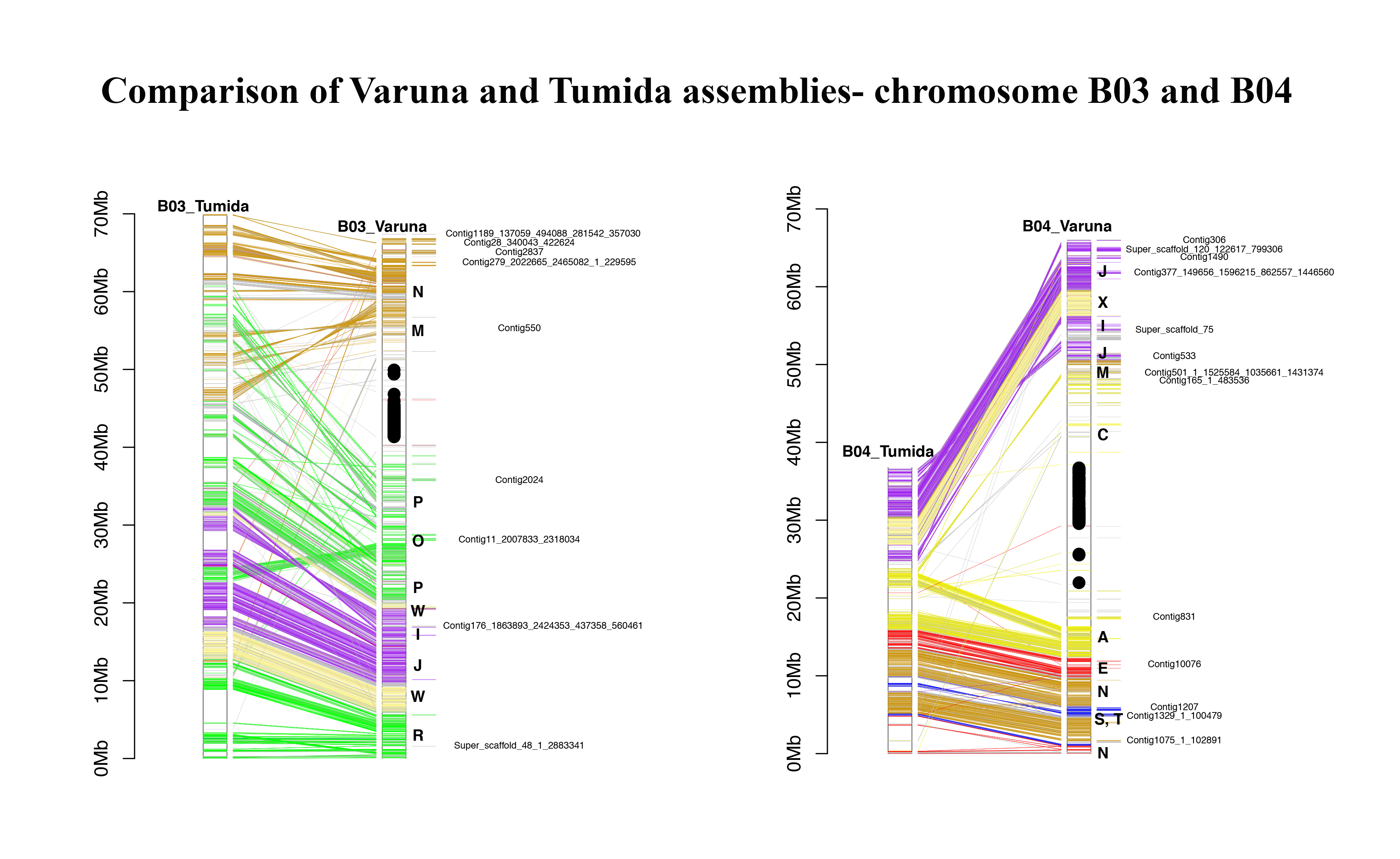

**Supplementary Figure 11 |** contd..

**Comparison of B05 - B08 pseudochromosomes- Tumida vs Varuna assembly**

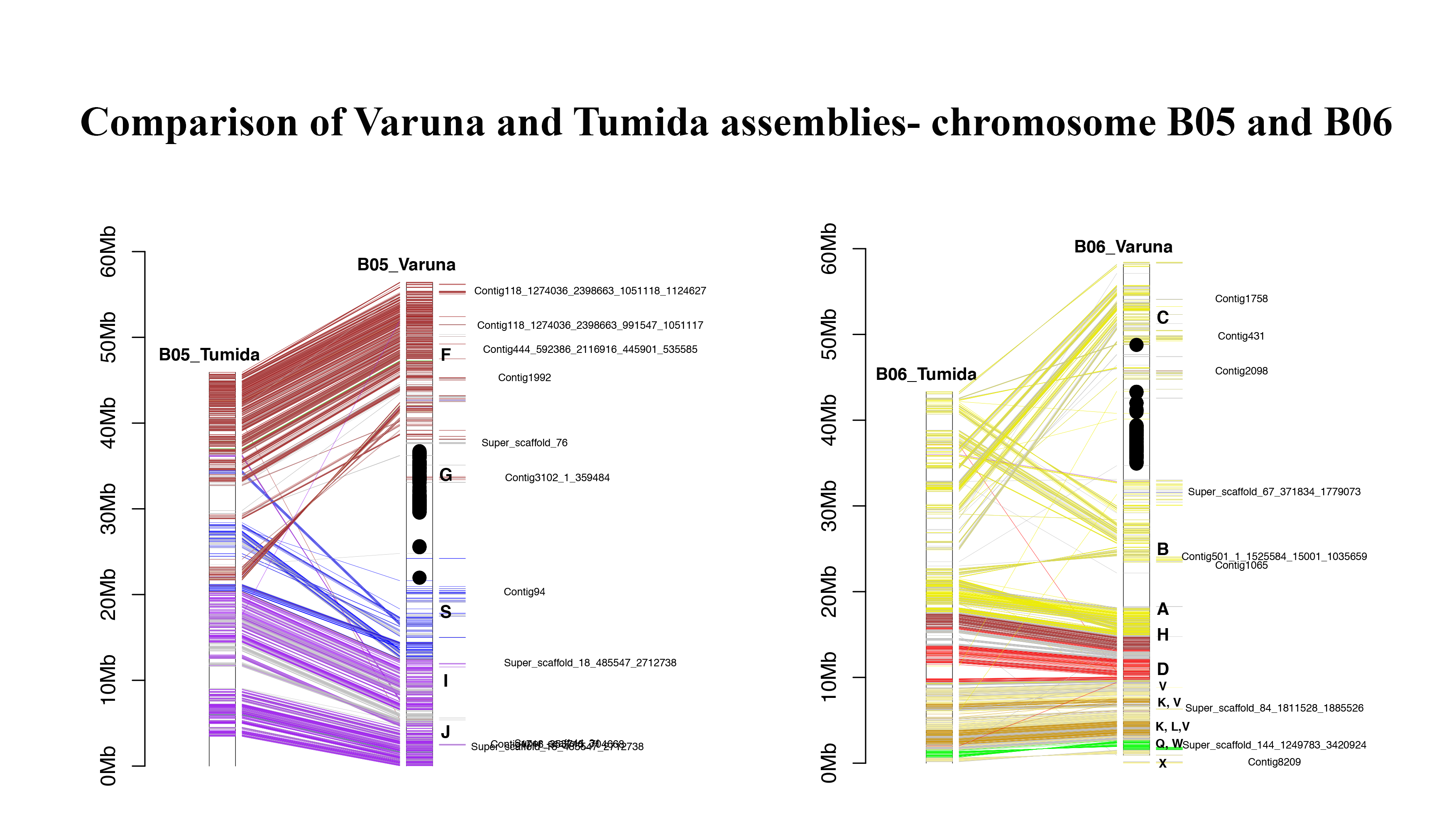

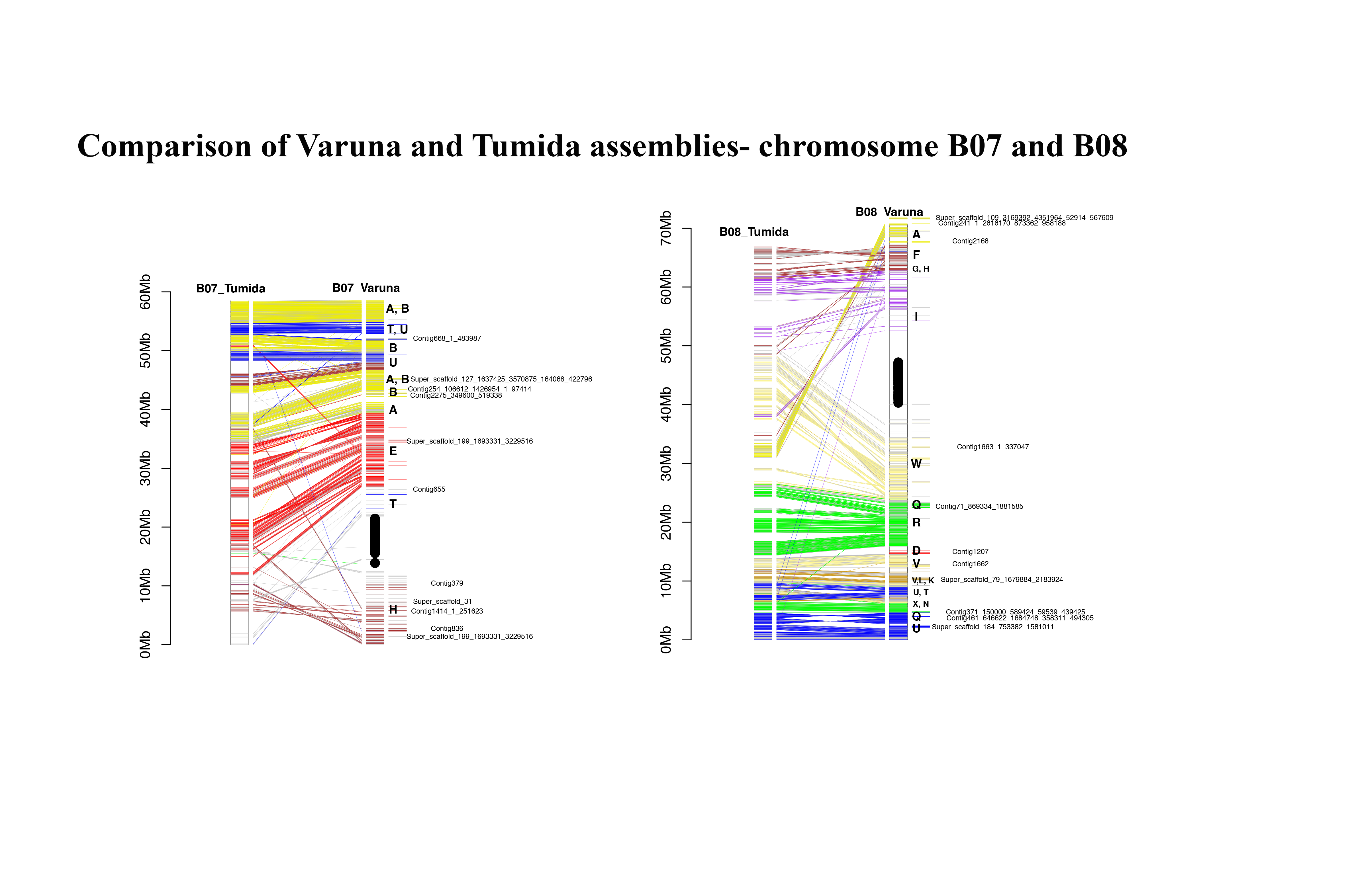

**Supplementary Figure 11 |** contd..

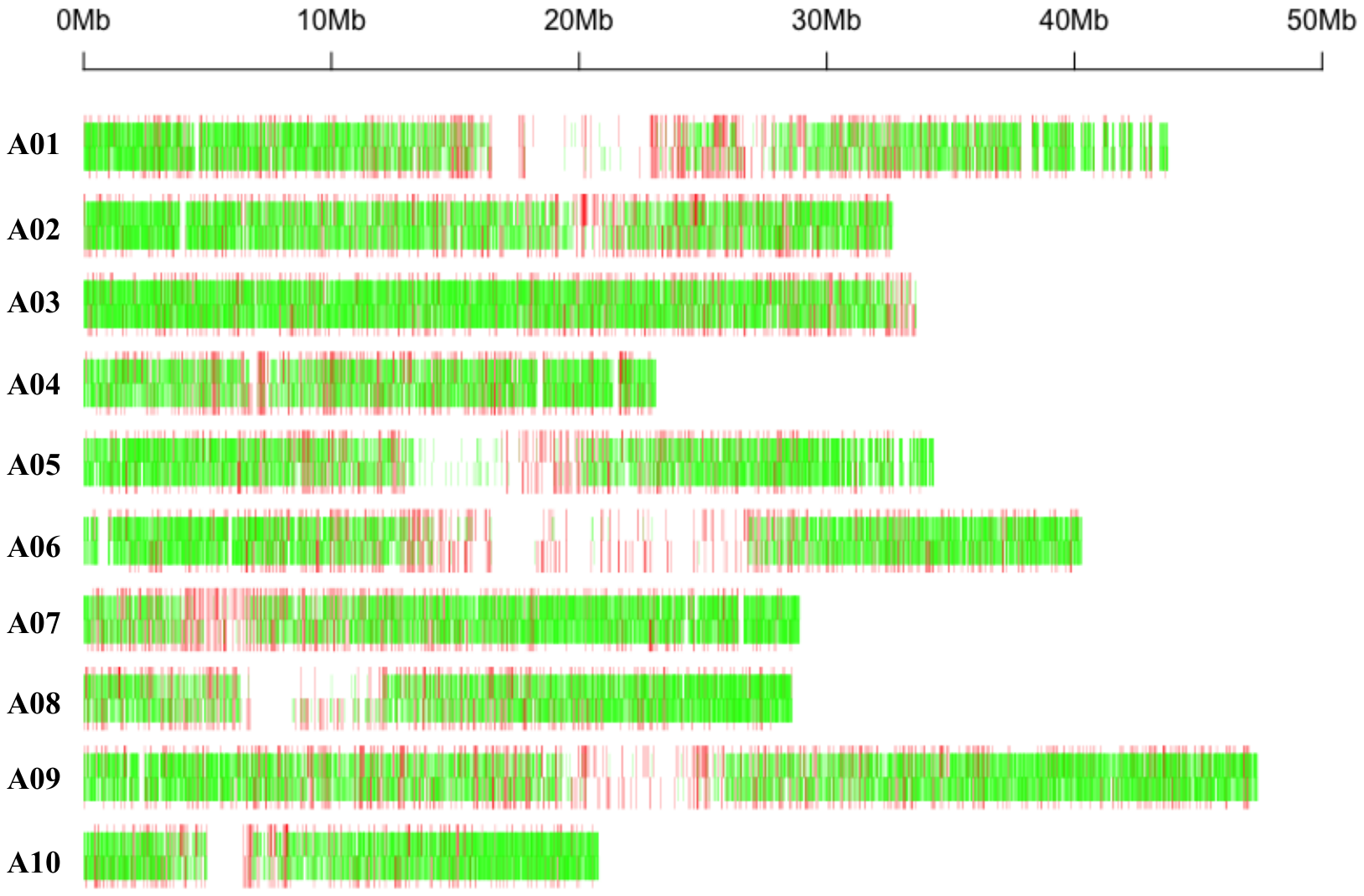
(a)

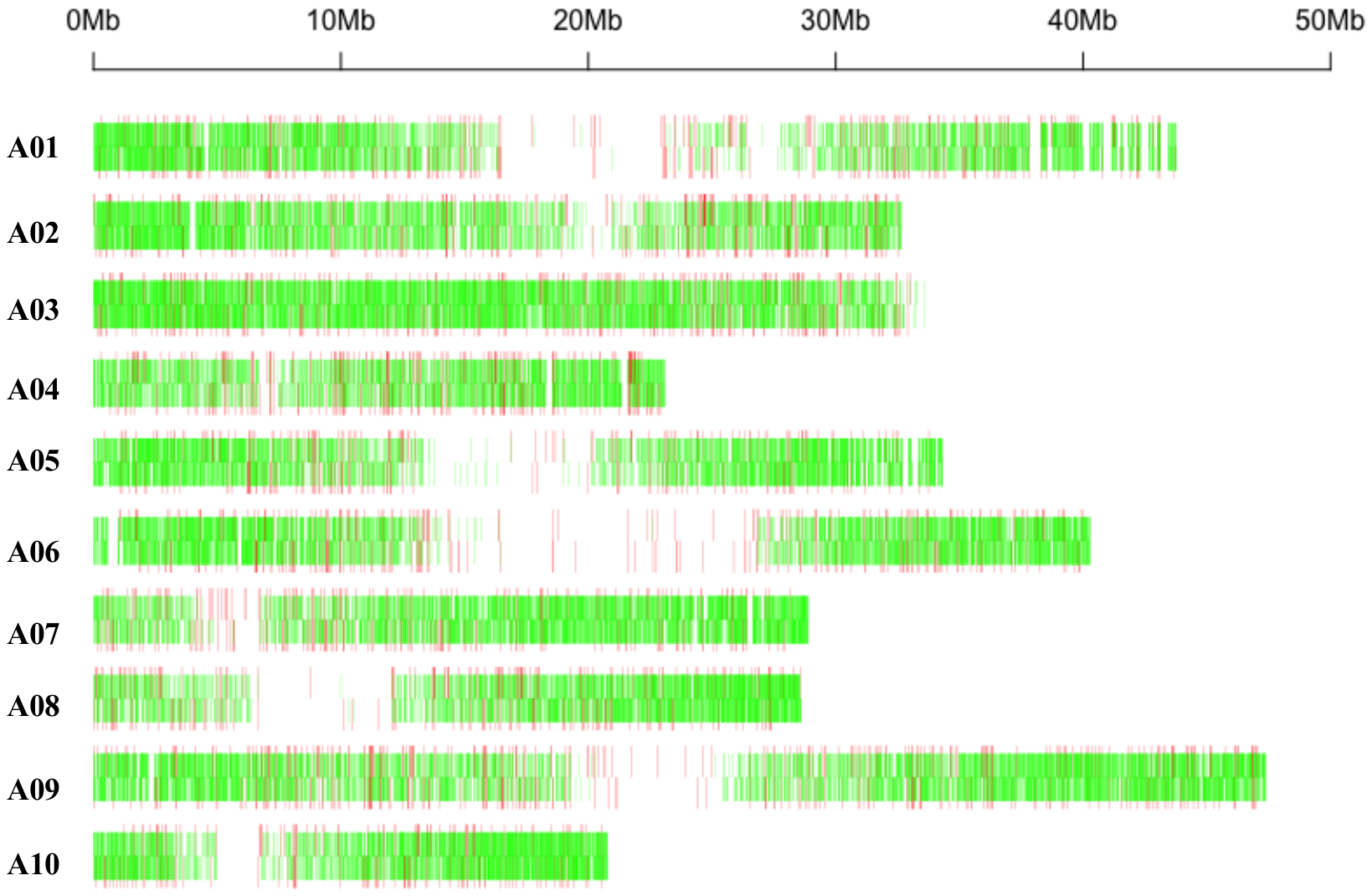
(b)

**Supplementary Figure 12 | Gene density on *B. juncea* A genome pseudochromosomes.** Green bars show genes that have orthologs in the B genome and *A. thaliana*. Red lines represent the A genome-specific genes – (a) total predicted genes (total 5,942) on A01-A10 pseudochromosomes (b) genes that were found to express in the transcriptome data (total 2,937).

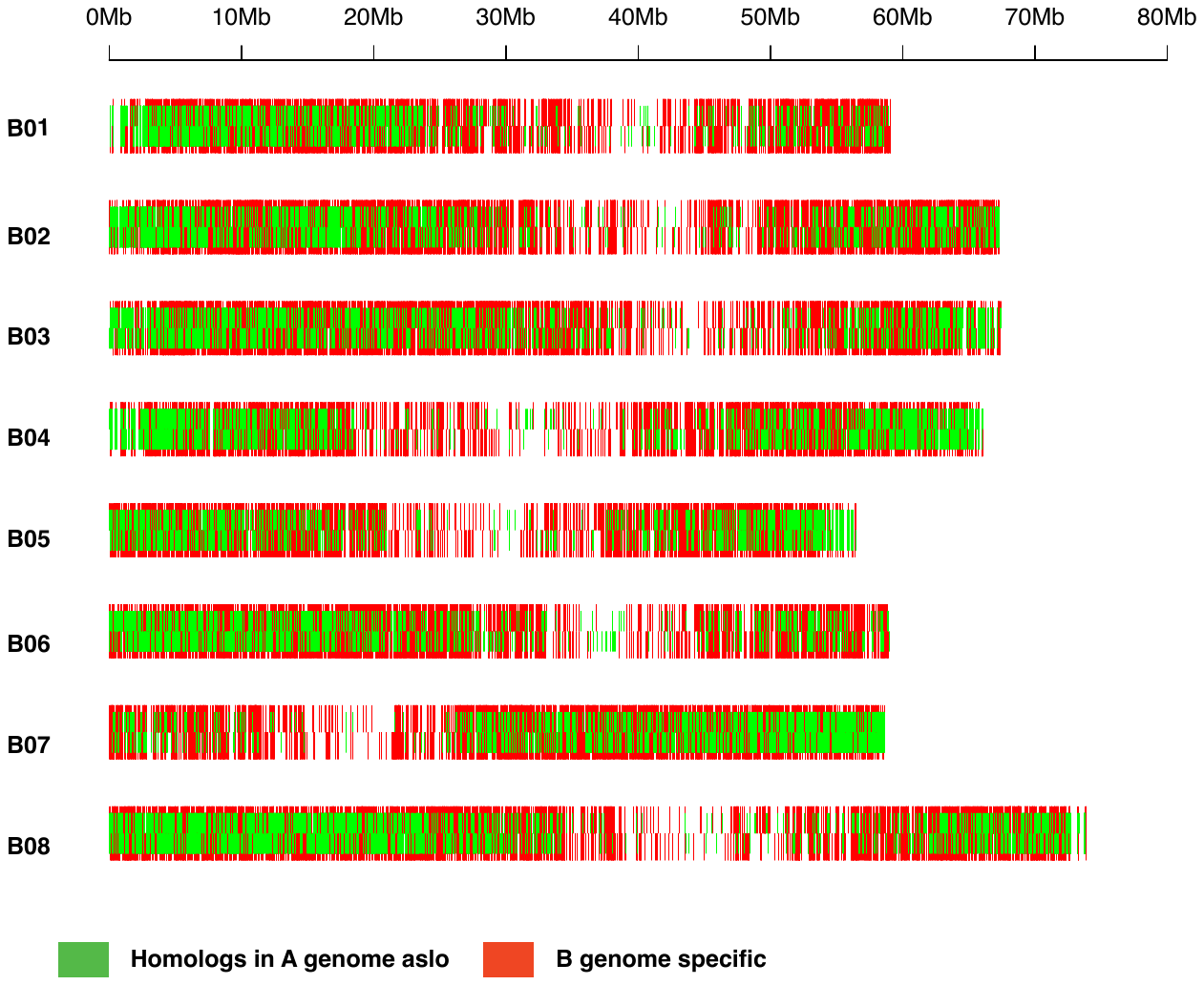

(a)

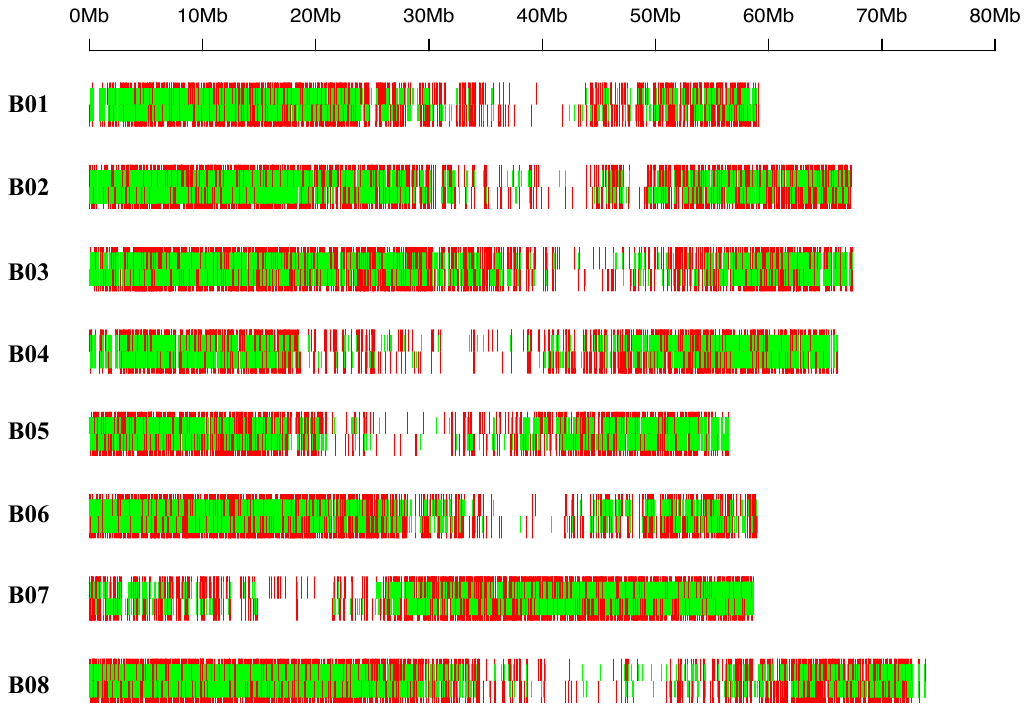

(b)

**Supplementary Figure 13 | Gene density on *B. juncea* B genome pseudochromosomes.** Green bars show genes that have orthologs in the A genome and *A. thaliana*. Red lines represent the B genome-specific genes – (a) total predicted genes (22,158) on B01-B08 pseudochromosomes, (b) genes that were found to express in the transcriptome data (total 12,262).

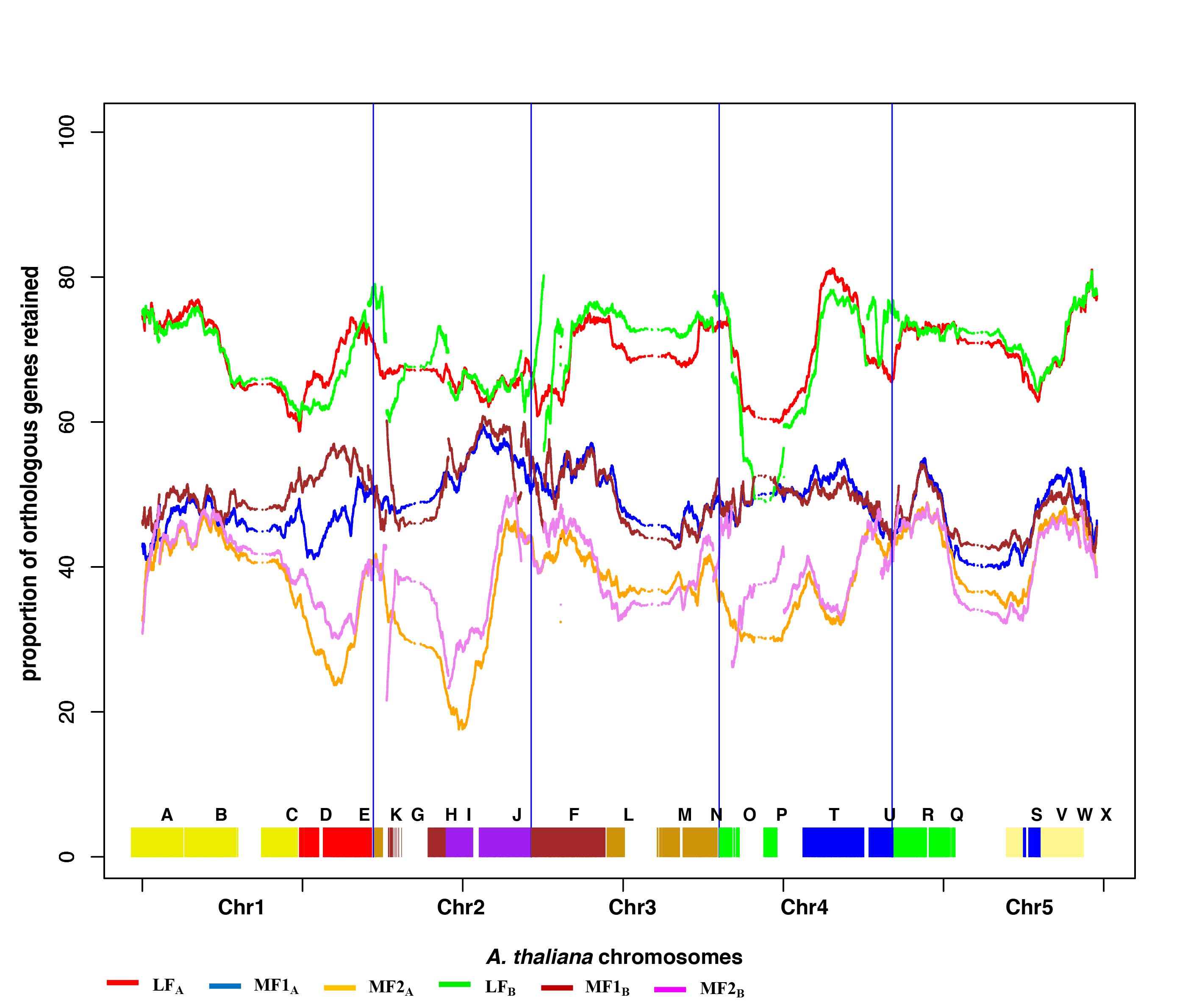

**Supplementary Figure 14 |** **Proportion of orthologous genes retained in the A and B genomes.** Gene retention/loss of the orthologous genes in the three paleogenomes (LF, MF1 and MF2) of A and B genomes of *B. juncea* corresponding to *A. thaliana* A to X gene blocks. The x axis shows the physical position of *A. thaliana* genes and gene blocks; the Y axis shows percentage orthologous genes retained in the three paleogenomes of the A and B genomes of *B. juncea*.

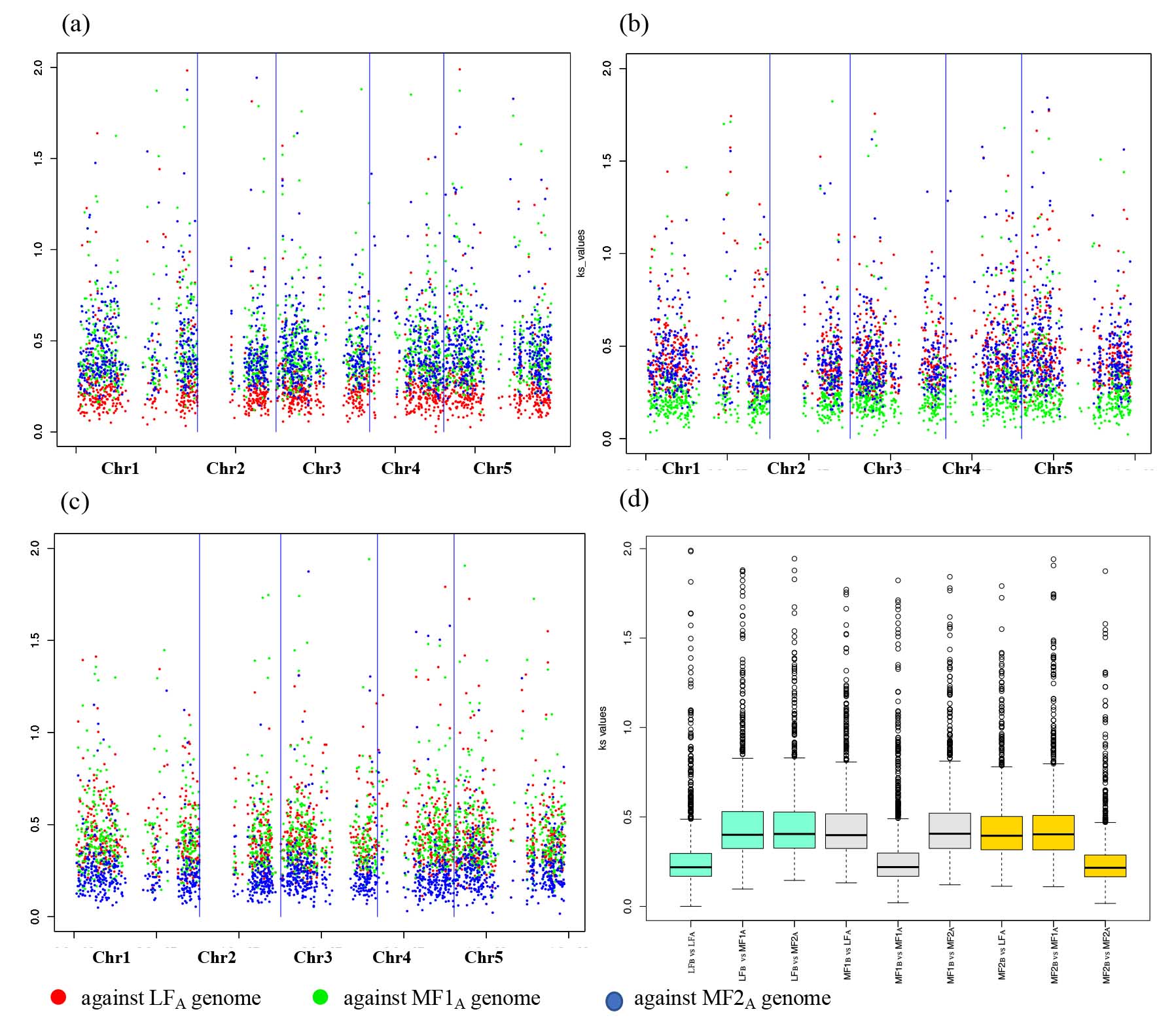

**Supplementary Figure 15 | Relationship of the three paleogenomes(LF, MF1 and MF2) of A and B genomes of *B. juncea* based on Ks values in 1428 genes present as a single copy in *A. thaliana* and three copies each in the A and the B genome.** Ks values were calculated between (a) LF (b) MF1 and (c) MF2 region genes of the B genome against the three syntenic genes in the A genome. The positions of the genes have been arranged on the basis of their physical position on the *A. thaliana* chromosomes (d) box and whisker plot showing inter-relationship of the three paleogenomes constituting the A and the B genome.

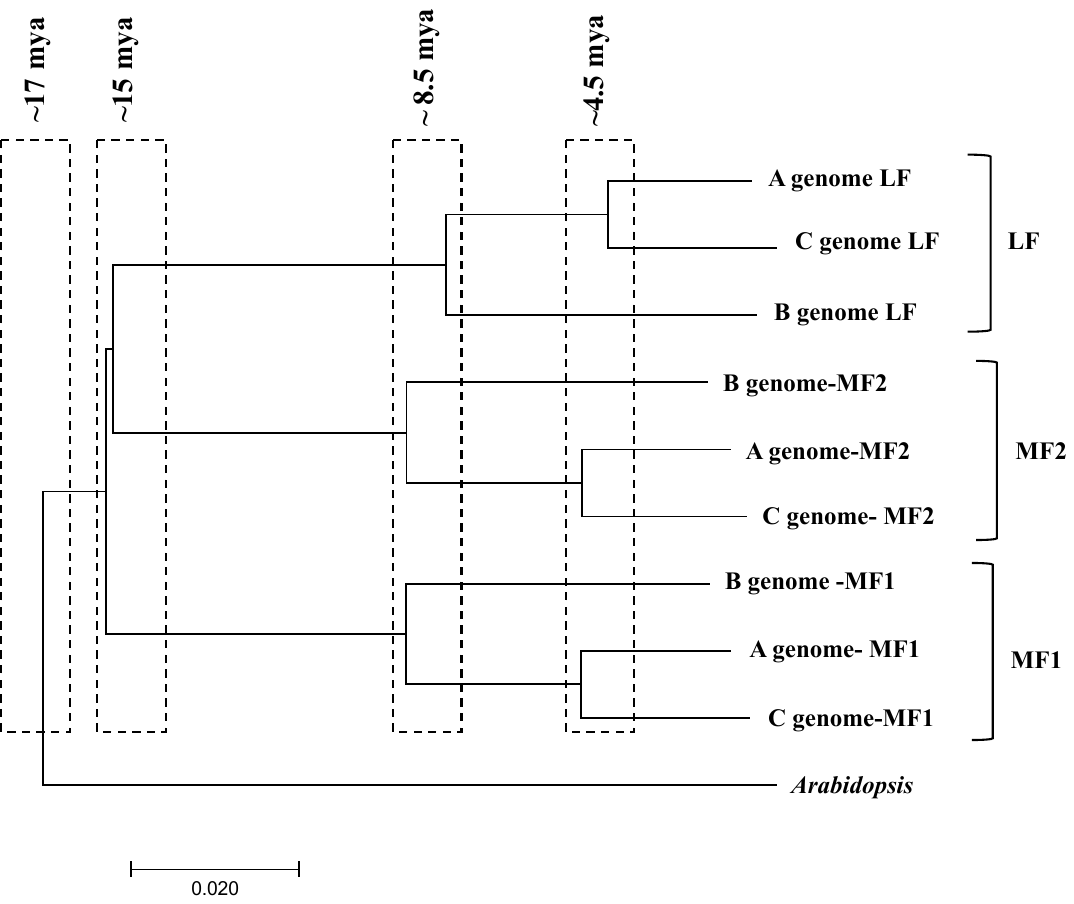

**Supplementary Figure 16 | Neighbour-joining tree based divergence analysis of the constituent paleogenomes of the A, B, and C genomes as compared to *A. thaliana*.** The tree was constructed on the basis of the distances estimated from synonymous substitutions in the concatenated CDS sequences of 1428 *Arabidopsis* genes that have been retained as three orthologs in the A and B genomes of *B. juncea* and the C genome of *B. oleracea*. A and B genome paralogs diverged at around 15MYA, whereas the homoeologs diverged ~8.5 MYA. A and C genome homoeologs separated around ~4.5 MYA.
